# On quantum computing and geometry optimization

**DOI:** 10.1101/2023.03.16.532929

**Authors:** Ashar J. Malik, Chandra S. Verma

## Abstract

Quantum computers have demonstrated advantage in tackling problems considered hard for classical computers and hold promise for tackling complex problems in molecular mechanics such as mapping the conformational landscapes of biomolecules. This work attempts to explore a few ways in which classical data, relating to the Cartesian space representation of biomolecules, can be encoded for interaction with empirical quantum circuits not demonstrating quantum advantage. Using the quantum circuit in a variational arrangement together with a classical optimizer, this work deals with the optimization of spatial geometries with potential application to molecular assemblies. Additionally this work uses quantum machine learning for protein side-chain rotamer classification and uses an empirical quantum circuit for random state generation for Monte Carlo simulation for side-chain conformation sampling. Altogether, this novel work suggests ways of bridging the gap between conventional problems in life sciences and how potential solutions can be obtained using quantum computers. It is hoped that this work will provide the necessary impetus for wide-scale adoption of quantum computing in life sciences.

## Introduction

Three-dimensional (3D) [1] protein structures, experimentally determined using biophysical techniques like nuclear magnetic resonance (NMR) and X-Ray crystallography [2], enable in-silico characterization of protein function. Routine techniques, like molecular dynamics (MD) simulations [3], Monte Carlo (MC) simulations [3] and molecular docking [4], build on 3D protein structure data and allow for determination of protein dynamics and substrate binding. This insight, achieved from in-silico methods [5, 6, 7], is crucial in setting the course of, what is usually expensive, biochemical characterization carried out using laboratory assays.

Given that in-silico characterization is now an established aspect in translational studies, going from molecular effect prediction to laboratory-based verification, it is all the more important that algorithms utilize the existing hardware to the maximum and evolve with the advancements in hardware. An example of this can be seen with the adoption of artificial intelligence/machine learning (AI/ML) based methods in nearly all computational areas, especially drug discovery; one of the primary drivers of which is the advancements in graphics processing units (GPUs) [8]. Development of new algorithms in non-AI/ML areas utilizing the parallel-compute capability of GPUs, has allowed for advancements in other areas e.g, exponential speed-up of MD simulations [9, 10, 11] which now allow for large molecular assemblies to be observed over longer time-scales, offering better insights into the dynamics of these biomolecules.

Based on the fundamental principles of quantum mechanics, quantum computing is an emerging area which demonstrates potential to tackle hard problems, which are beyond the current capabilities of classical computers [12, 13]. Current state-of-the-art includes efforts in the area of quantum chemistry simulations [14, 15], machine learning [16, 17] and finance [18], with new algorithms, continuously being added to extend current capabilities e.g., QPacker [19], tackling the protein design problem and others [20, 21] for protein folding. Unlike scalability in the classical computing ecosystem, where new technologies are easily integrated and algorithms easily scaled to utilize the advance e.g., in the case of the use of GPUs with classical algorithms in the area of molecular dynamics simulations and machine learning to name a few, integration of existing algorithms with quantum computing is non-trivial. While advancing at a significant rate, requiring a radical rethink about how algorithms are designed and deployed, quantum computing at present remains non-intuitive for the classical programming fraternity.

Quantum algorithms aim to achieve quantum advantage, which simply put, is the ability to solve problems that classical computers struggle with [22, 23, 24]. A quantum algorithm can broadly be decomposed into three parts, a) the data encoding step, b) the use of the quantum hardware to solve a particular problem of interest and lastly, c) converting the results into a form which is readily understandable, with steps “a” and “c” being inextricably linked. All aspects of quantum algorithms face challenges due to the non-intuitive nature of the hardware executing these.

In this work the area of molecular geometry optimization is explored. The work focuses on illustrating, without quantum advantage, encoding of classical molecular data for use with quantum hardware. Problem areas where classical data can interact with quantum hardware are demonstrated. In particular, this work demonstrates, using an empirical quantum circuit, optimization of positions of atoms and distances between them, a task frequently carried out in classical molecular mechanics, which demonstrates all three parts of a quantum algorithm. Dihedral data is used with a quantum support vector classifier to introduce machine learning capabilities. Additionally, empirical rotamer sampling is demonstrated using quantum Monte Carlo simulations.

To the best of our knowledge, this work is a first in presenting quantum models that work with data from the area of classical molecular mechanics. Although achieving quantum advantage is beyond the scope of this work currently, it is expected that this work will act as a primer for new users. By introducing a method to translate classical molecular data for use with quantum computers and giving examples of problem areas where classical data can interact with quantum hardware, it is assumed that this work will help achieve solutions to classical problems beyond the capabilities of classical computers.

## Method

This work makes use of the open-source software development kit (SDK), Qiskit [25], using Python 3.7 through Anaconda. All work was carried out using quantum simulators. VMD [26] and NAMD [27] are used for handling and analysis of protein structure data.

In this work a number of models of quantum computation are presented. Briefly, these models demonstrate how classical molecular data can be encoded for use with quantum computers and how certain problems can be explored. These models are briefly introduced below.

### Model 1: Encoding classical molecular data and molecular mechanics

Cartesian coordinates are commonly used when recording the positions of atoms, alone or constituting molecules. Three dimensional cartesian (*x, y, z*) coordinates can be readily converted to their equivalent spherical (*r, θ, φ*) coordinate forms, see Figure 1(a), using empirical mathematical transformations.

**Figure 1:**
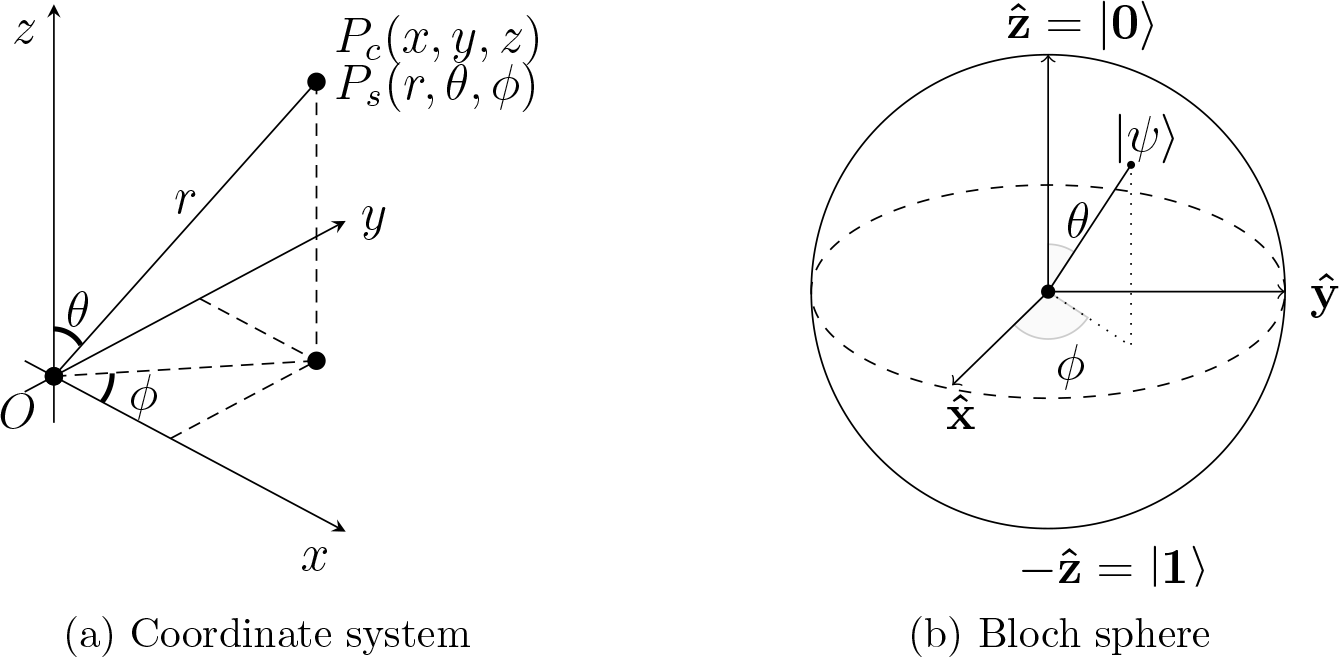
Coordinate systems and the Bloch sphere. a) Any cartesian coordinate (*x, y, z*) can be uniquely expressed as a spherical coordinate (*r, θ, φ*), where *r* is the length of the line segment, and *θ* and *φ* are angles measured from the *z* and *x*-axes respectively.b) The Bloch sphere showing the state of a qubit *ψ* given by Equation 1 and can be set by using the UGate gate in the Qiskit SDK by setting the respective angles (*θ* and *phi*).

A qubit state, represented by the Bloch sphere, see Figure 1 (b) is denoted by

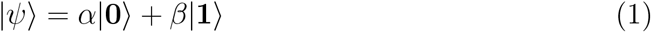

where *α* and *β* are probability amplitudes.

A qubit can be set to any arbitrary state, on the Bloch sphere using the generic single-qubit rotation gate, UGate, which accepts three Euler angles (*θ, φ, λ*) as inputs.

Classical molecular mechanical algorithms optimize molecular geometries by introducing variations in the observed state such that interatomic relationships converge to reference measures recorded, for the said atoms, in the force field [28]. These quantities comprise bonded terms, i.e., bonds, angles, dihedrals and non-bonded terms, i.e., van der Waal and electrostatic. This optimization can be reduced to a simple analogous problem where two vectors (a subject and a reference) are used to represent two points and both the magnitude and direction of the subject vector are modified to approach the magnitude and direction of the reference vector. For a multi-atom system represented by vectors, the convergence of the magnitude (distance between atoms) to a reference value alone (listed e.g., in the force field) approximates the optimization effect from the bond term or other quantities that require an ideal separation between atoms. Together with the vector direction, all interactions for three-atom or larger systems can be optimized.

For the simple vector magnitude and direction convergence problem introduced, this work makes use of the CSwapGate to calculate the dot product of two qubit states [1, 29]. While different circuit topologies are possible, this work uses a variational circuit setup. Figure 2 shows the quantum circuit used.

**Figure 2:**
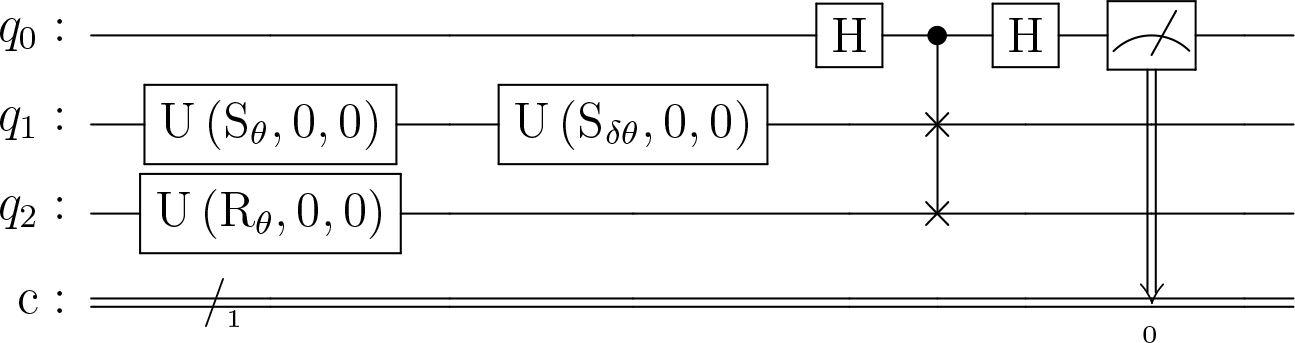
Quantum circuit using the CSwapGate. The circuit uses three qubits, where *q*_0_ is the control line employing two Hadamard gates and from which measurement is made. The encoded data is loaded as angles onto *S*_*θ*_ and *R*_*θ*_, where *S*_*θ*_ takes the subject value and *R*_*θ*_ takes the reference value. *S*_*δθ*_ acts as the variational parameter which is controlled by the optimizer.

The pre-processing step, which prepares the data, starts with computing the magnitude and direction of the subject and reference vectors. The calculated magnitude is normalized to the unit scale for encoding as an angle using the UGate. This work arbitrarily chooses to normalize the magnitude using the scheme shown below

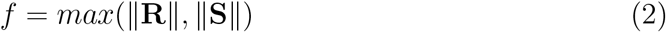

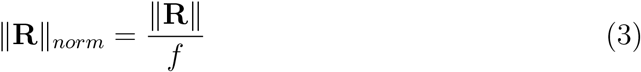

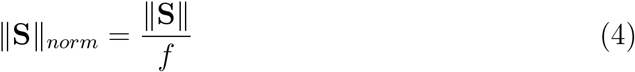

where ||**R**||, such that {**R** ∈ ℝ : **R** ≠ 0}, and ||**S**||, such that {**S** ∈ ℝ : **S** ≠ 0}, are magnitudes of the reference and subject vectors and ||**R**||_*norm*_ and ||**S**||_*norm*_ are their normalized counterparts which are then directly used, in radians, as angle inputs for the UGate. In the case of direction, the angle quantity is directly used as input to the UGate without normalization.

As the circuit is used in a variational form, the SPSA [30] optimizer available in Qiskit is used to perturb the variational parameter with the minimizer acting on the transformed dot product, see below. As introduced [1, 29], the equation to calculate the dot product is given by

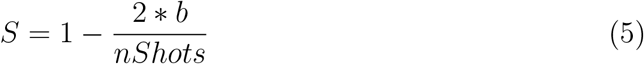

where “b” is the number of shots that result in the state “1” and nShots is the total number of shots attempted. The quantity “S” will approach “1” when the qubits are in the same state. This value is transformed by subtracting from one, see Equation 6, for use with the minimizer.

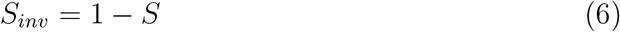

### Model 2: Vector alignment for planar molecular geometry optimization

The previously discussed method for magnitude optimization was used to refine the irregular sides of an arbitrary hexagon such that they become regular. To this end, an irregular-hexagon was randomly generated, and an arbitrary reference value was used to transform the shape into a regular hexagon.

### Model 3: Vectors and protein structure alignment

The method based on direction convergence discussed previously is used to align two structures of the protein ubiquitin (RCSB PDB ID: 1ubq), where one is the crystal structure and the other has undergone an arbitrary rotation in three dimensional space using VMD. Using the orient package in VMD, principal axes are calculated for both the original and rotated protein structures. The principal axes provide three unit vectors for each structure, the directions of which are systematically aligned across both structures. After direction convergence, new vectors and required transformations are computed and applied to the rotated structure to map it to the original structure. An all-atom root mean square deviation (RMSD) is reported to assess the quality of the protein structure alignment.

### Model 4: Using variational quantum classifier for side chain rotamer classification

In order to test the variational quantum classifier, the ubiquitin protein structure (RCSB PDB ID: 6l0l) was used to create a dataset. For this, the sidechain of each amino acid was rotated, using VMD, about the bond CA-CB bond axis, excluding the amino acids glycine, alanine and proline. The rotations were carried out in 0.1 degree increments, creating 3,600 rotamer conformations (observations) per amino acid which were then saved as PDB files, with the only difference between the crystal structure and the new PDB structure being the single amino acid rotamer change. For each new conformation, a reference atom from the sidechain of the rotated amino acid was chosen, see supplementary Table 1 for a list of amino acids and their reference atoms. The number of atoms within certain select cutoffs (namely 5Å, 4Å, 3Å, 2Å) proximity were enumerated. These numbers act as features for the machine learning models used in this work. For each of the 3,600 observations, per amino acid, total potential energy of the system was calculated making use of the NAMDEnergy plugin in NAMD with the charmm36 force field. The requirement of potential energy for this model required the protein structure to have hydrogens which were missing in ubiquitin structure previously selected. A solution-NMR (nuclear magnetic resonance) based structure which already had the hydrogens (PDB ID: 6l0l) was therefore used.

For use with a classification model, the energy values were divided into two classes, stable and unstable, with energy values at or below “0” kcal/mol being stable and those above “0” kcal/mol being unstable. The label for each class was “1” if a particular rotamer conformed to that class or “0” otherwise. In all instances, both labels were assigned. For model training, to ensure a balanced dataset, only amino acids where 50 values for each class were available, were chosen and correspondingly a 100-observation balanced dataset was generated.

The model used the ZZFeaturemap together with the EfficientSU2 ansatz with the four features mentioned above making use of the “rx” and “y” gates with circular entanglement, and employing the COBYLA optimizer. The 100-observation balanced dataset was split with 70% data used for training and 30% for testing. The same training and testing data was also used with a classical support vector classifier and the classification accuracy was determined for both the variational quantum classifier and the classical support vector classifier. As another performance measure the entire dataset of 3,600 observations per amino acid was used for prediction and the accuracy scores for both classical and quantum algorithms were computed.

To ascertain if the model training was robust, three trials each were carried out, per amino acid, and the accuracy determined for both the quantum and classical classifiers.

### Model 5: Quantum Monte Carlo simulation for rotamer energy landscape profiling

The Monte Carlo method was used to sample the rotamer energy landscape. A six-qubit string was used to represent a total of 64 (2^6^) states, each corresponding to a rotamer conformation of an amino acid, with consecutive states representing a difference of ∼ 5.6 *deg*. The rotamers for each amino acid of ubiquitin (RCSB PDB ID 6l0l) were generated using the method stated earlier. The quantum circuit was empirically driven using six Hadamard gates, for the random generation of each of the six-qubits to represent a new state.

The transition of the states were accepted with a probability of “1” if the new state was more stable, that is the change in energy (*Newstate − Oldstate*) is lesser than or equal to “0” (*δE ≤* 0), or using the rule:

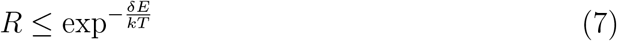

where *kT* was set to “1” and *R* is a random number, in the event that the *δE* > 0. The simulations were carried out for each amino acid for a total of 1000 Monte

Carlo moves. The states and their corresponding energies were recorded. For illustration purposes energy values were transformed using

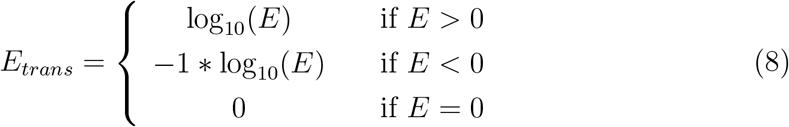

where “*E*” is the energy and “*E*_*trans*_” is the transformed energy used for generating plots.

## Results

### Model 1: Encoding classical molecular data and molecular mechanics

Vector magnitudes and directions were compared across two vectors, a subject and a reference vector. Figure 3 shows a particular instance of this where the difference (*abs*(|*reference*| *−* |*subject*|) = 13.29*AU*) between the magnitudes of the subject (|*subject*| = 25.98*AU*) and reference (|*reference*| = 12.69*AU*) vectors minimized to zero as the quantity (*S*_*inv*_) in Equation 6 is minimized using the quantum circuit illustrated in Figure 2.

**Figure 3:**
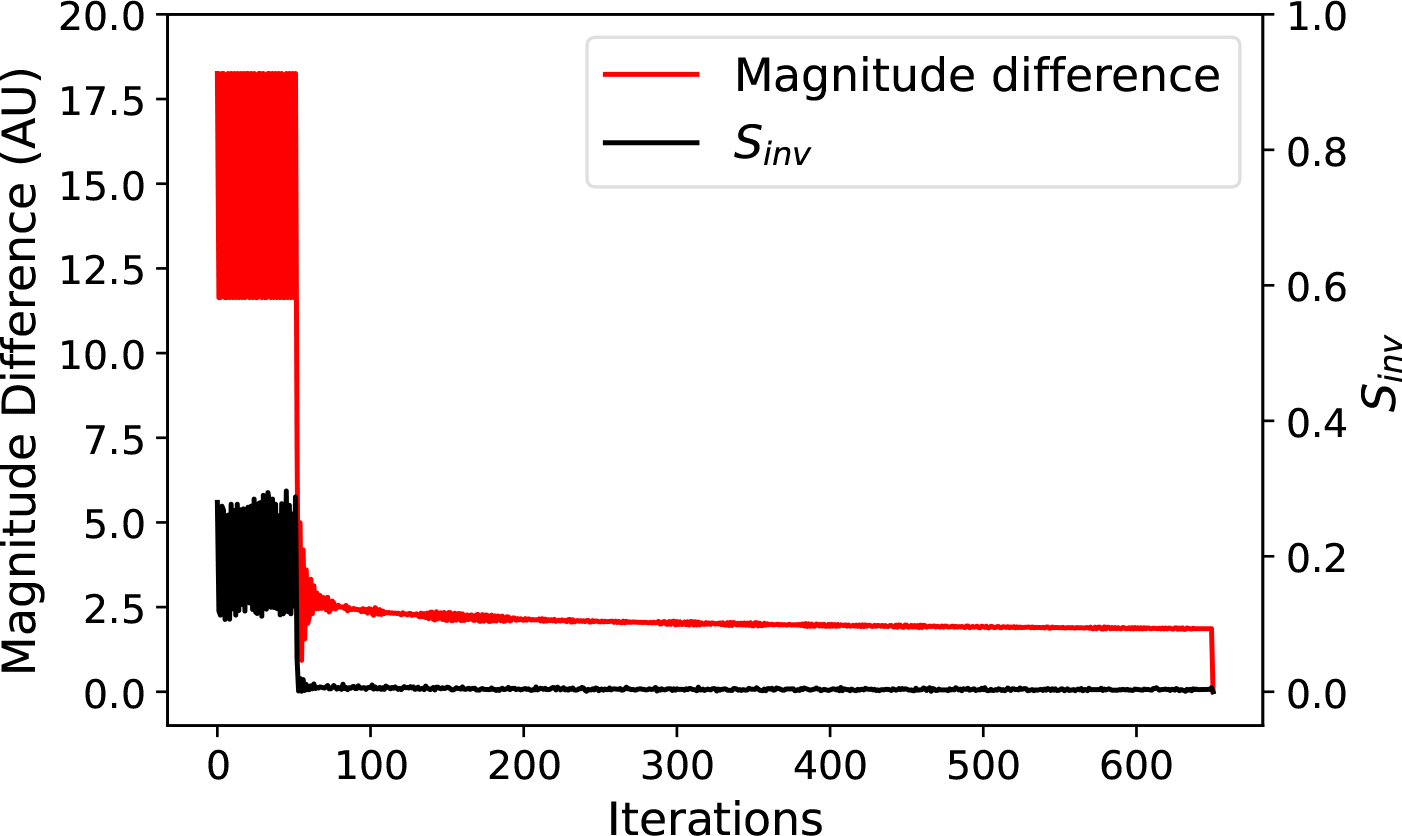
Minimization of magnitude difference and *S*_*inv*_. The difference in the magnitudes of the subject and reference vector approaches “0” as the minimizer reduces *S*_*inv*_. From the quantum circuit in Figure 2, the minimizer alters *S*_*δθ*_ such that the state of the qubit carrying the magnitude of the subject vector approaches the state of the reference qubit, in turn resulting in magnitude of the support vector converging to the magnitude of the reference vector.

Two angles (*θ, φ*) are required to uniquely represent the direction of a vector in the spherical coordinate system. Figure 4 demonstrates results of using the same circuit as used for the magnitude to ensure directions align between the subject (*θ* = 1.15rad, *φ* = 2.03rad) and the reference (*θ* = 0.73rad, *φ* = 0.98rad) vectors by minimizing the difference between them.

**Figure 4:**
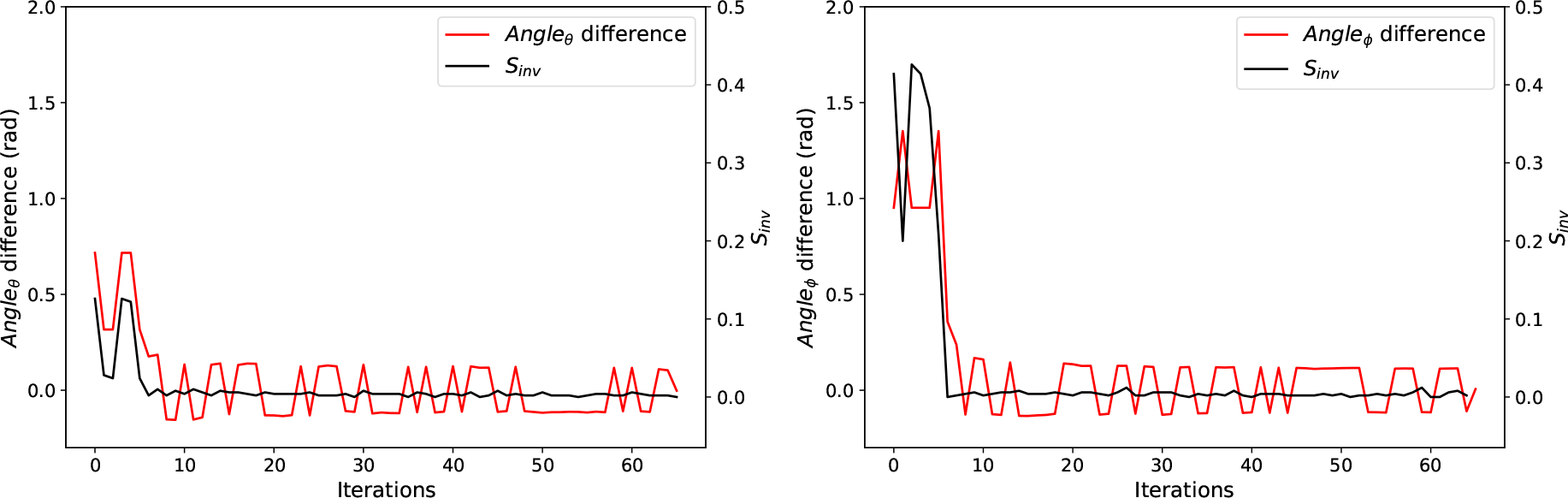
Minimization of angle difference and *S*_*inv*_. To uniquely represent a vector, two angles (*θ, φ*) are needed. Using the same quantum circuit as shown in Figure 2, the difference in the individual angles of the subject and reference vector are minimized as the minimizer reduces *S*_*inv*_. For ease of illustration, only every 10th iteration is plotted.

While in this work only one case each for magnitude and direction is illustrated, the method can readily be adapted to explore other cases.

### Model 2: Vector magnitude for planar molecular geometry optimization

This model presents optimizing the sides of a hexagon as a use case for the quantum circuit and the classical data encoding method discussed above. This problem bears similarity with refinement of bond length of planar molecules, e.g., benzene. In this example, all sides are chosen to have an arbitrary length of 1.5*AU*. Figure 5 shows the starting (Blue) and final (Red) states of the system, and illustrates that the final state adopts a regular hexagonal topology.

**Figure 5:**
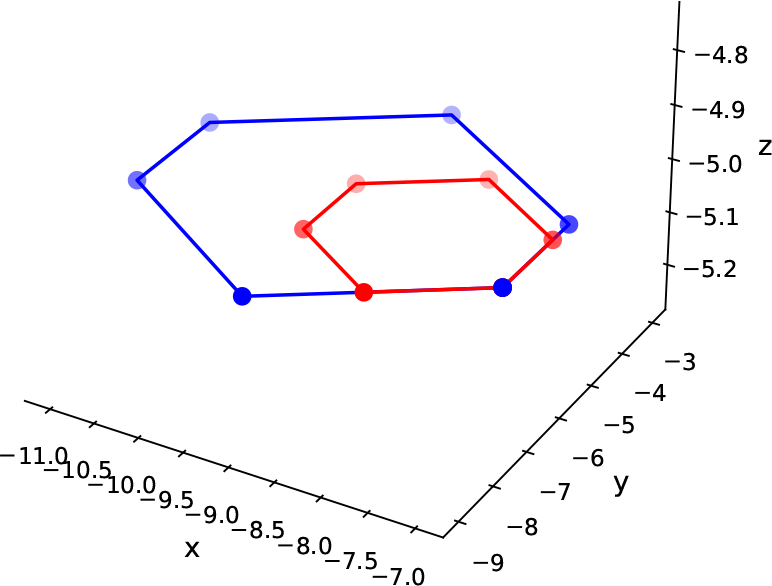
Optimization of the sides of a hexagon. The quantum model demonstrated above is used to optimize the sides of a hexagon. The blue figure represents a randomly generated hexagon, whose sides are modified to be regular (1.5*AU*), using the magnitude difference minimization method detailed earlier.

### Model 3: Vectors direction alignment and protein structure alignment

Protein structure alignment is presented as a use case of aligning vector directions. Figure 6 shows the rigid transform of a protein structure, with the transform comprising 3D rotations about the axes (*x* = 45*deg, y* = 35*deg, z* = 25*deg*). To achieve an alignment between the transformed and original structure, the direction alignment model is used. Figure 6 shows the result of the alignment with an RMSD value of ∼ 0.4Å.

**Figure 6:**
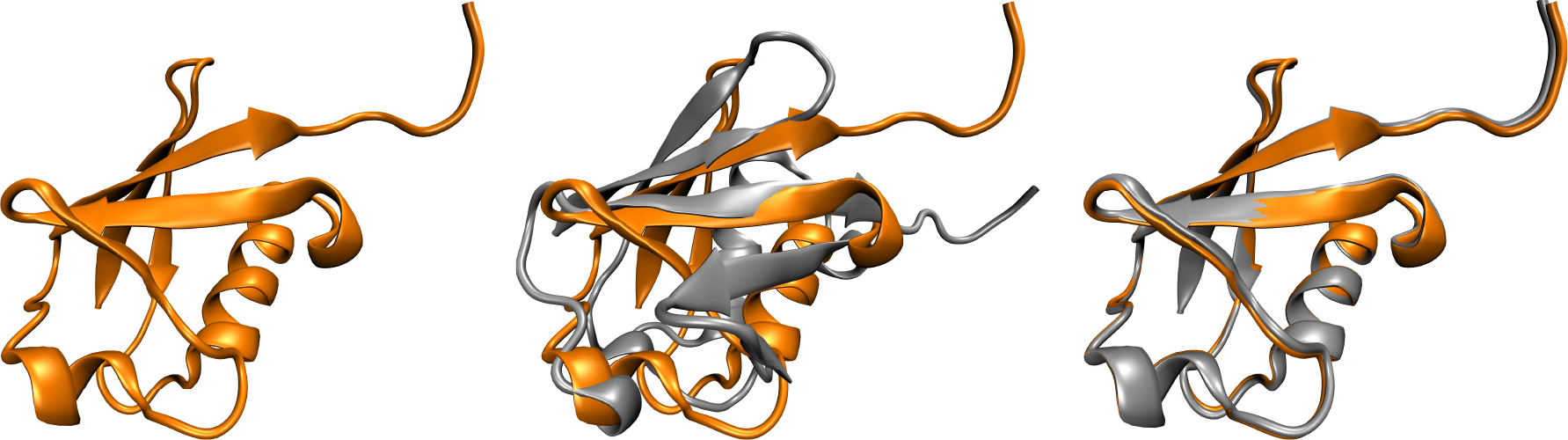
Protein structure alignment. The crystal structure of ubiquitin from RCSB PDB (left; orange) was arbitrarily rotated about *x, y* and *z*-axes (middle), with the rotated structure shown in gray. Using the vector direction alignment model, the rotated structure was aligned with the original structure (right). The alignment resulted in an all-atom RMSD of 0.4 Å.

As stated earlier, while only one case is demonstrated in the work, the method can be used for rigid alignments between transformed structures.

### Model 4: Using variational quantum classifier for side chain rotamer classification

To generate the dataset for this model, the ubiquitin protein structure, comprising 82 amino acids was used and 3,600 sidechain rotamer conformations (observations) were generated for all amino acids excluding the glycine, alanine and proline as listed in the methods section. Subsequently only 45 amino acids were used as only these allowed creation of balanced 100-observation datasets, of which 50-observations had the classification “stable” and the other 50-observations “unstable”. Remaining amino acids had less than 50 out of 3,600-observations in either “stable” or “unstable” classes.

Figure 7 shows the comparison of the classification accuracy for the 45 amino acids using both the quantum and the classical classifiers. Additionally the classification accuracy achieved during the model training stage is shown alongside the classification accuracy achieved after using the same trained model to classify all 3,600 rotamers which included 100-observations from the training data and 3,500 observations which were novel for the model.

**Figure 7:**
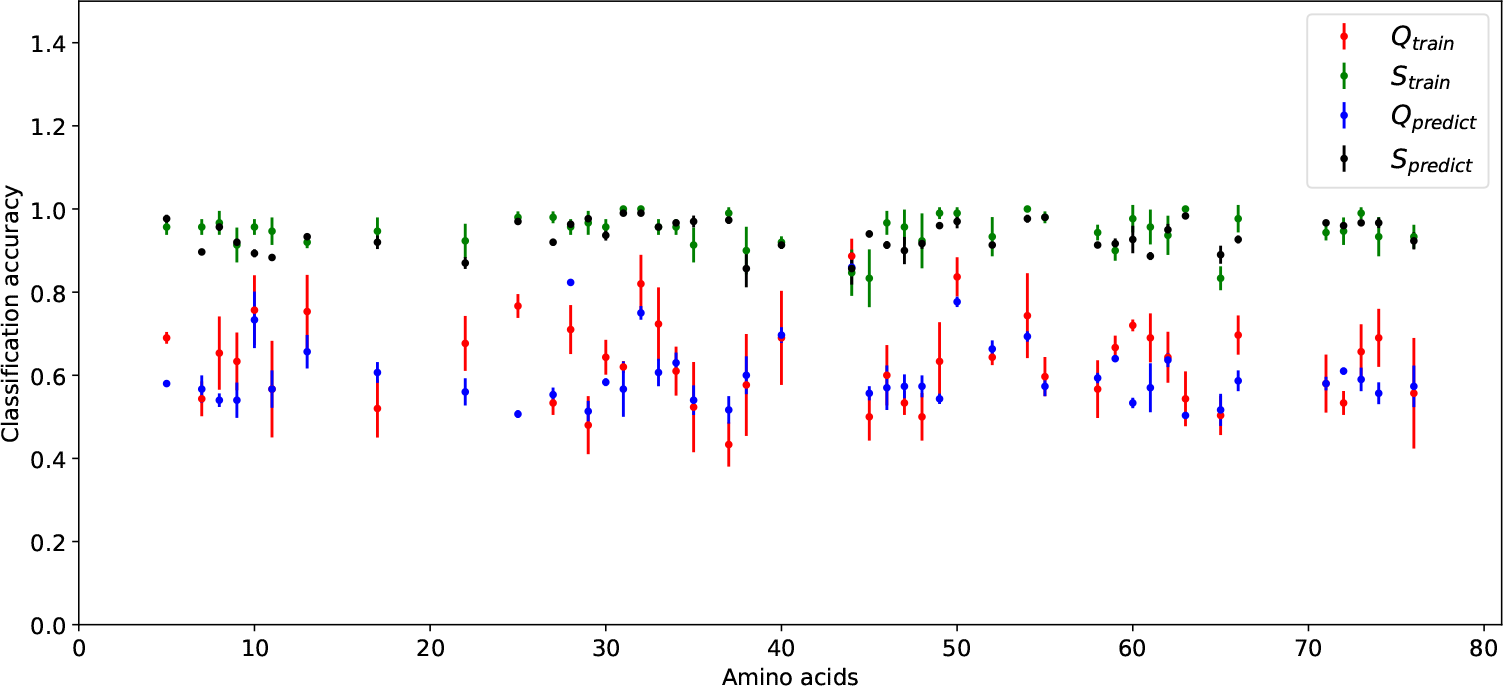
Comparison of classical and quantum classifiers. Overall the classical classifier performs better (*S*_*train*_, *S*_*pred*_) than the quantum classifier (*Q*_*train*_, *Q*_*pred*_) both on the training and prediction datasets. Three runs are carried out for training and prediction, with the quantum-based classification showing a much higher spread of accuracy across trials both for training (red) and prediction (blue).

For each dataset, both training and prediction, three trials were conducted to gauge robustness of the classifiers compared.

Overall the classical classification model outperformed the quantum classifier. With an average scoring accuracy of ∼ 60% across both training and prediction datasets, the quantum classifier shows a much higher spread of prediction accuracy. (c) Amino acid # 64

### Model 5: Quantum Monte Carlo simulation for rotamer energy landscape profiling

Using a 6-qubit string, 64 rotamer states were represented for each amino acid, with each state ∼ 5.6 *deg* apart from its next subsequent state. The energy of each new state was computed and the new state accepted or rejected using the rules listed in the methods section. Figure 8(a,b) show the case of two amino acids (66 and 72), both of which clearly show an inverse relationship between states visited (Figure 8: left panel) and their corresponding energetic signatures (Figure 8: right panel). For the 1000 Monte Carlo step simulation carried out for each amino acid, high occupancy (Frequency) is seen for low energy states and vice versa. Another case of amino acid 64 is shown in Figure 8 (c), where all energy values (*E*_*trans*_) sampled are below “0”, resulting in uniform occupancy for all rotamer states. The results for all amino acids are included in the supplementary material which reproduces similar expected behavior.

**Figure 8:**
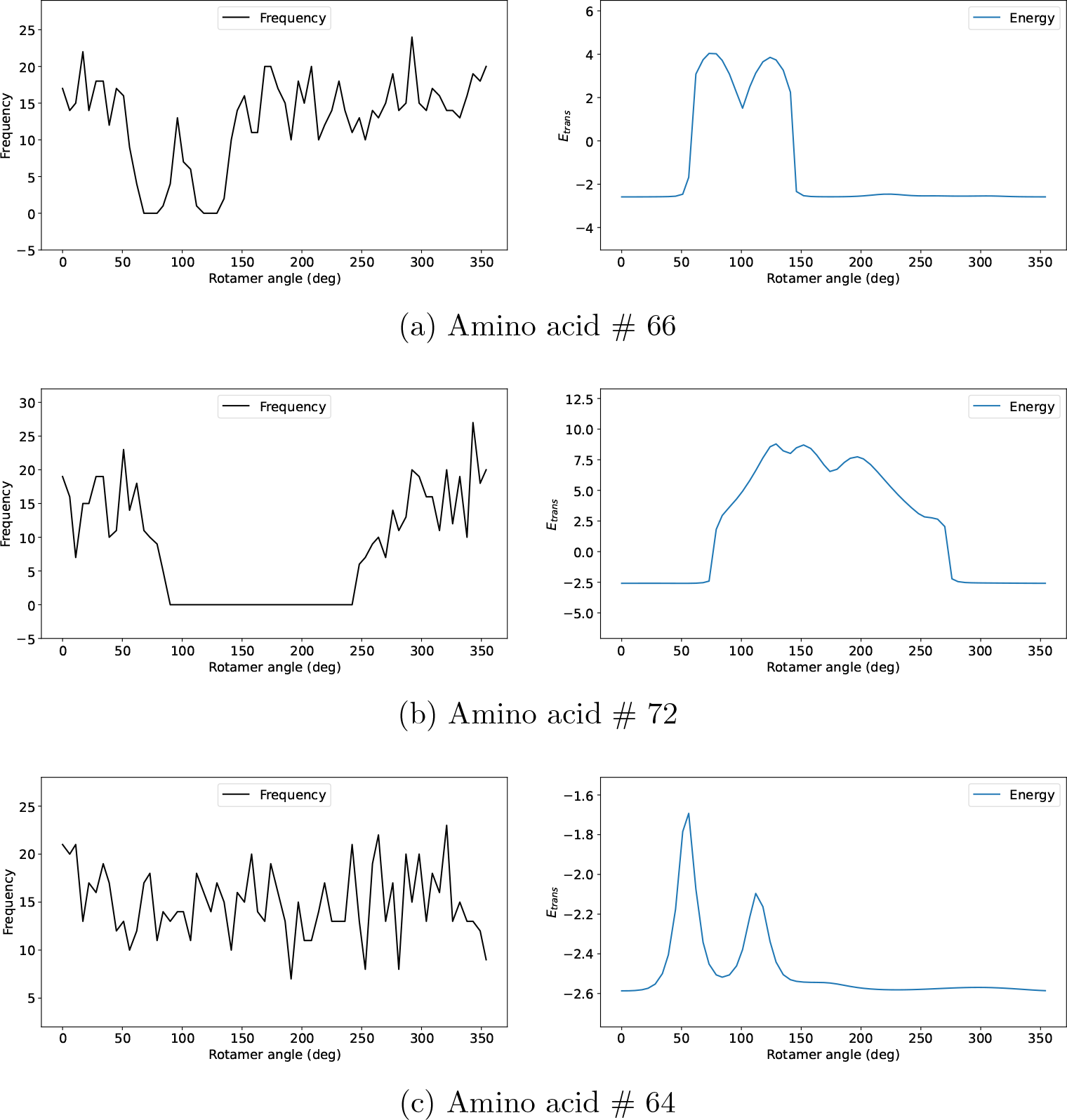
Monte Carlo simulation for rotamer energy profiling. Results of the states sampled using Monte Carlo simulation are shown for three amino acids. Amino acids 66 and 72, show an inverse correlation between rotamer energy and the occupancy profile as expected. For amino acid 64, given all rotamer states have *E*_*trans*_ *<* 0, (a) all states are uniformly visited.

## Discussion

This work, using empirical models, demonstrates how classical data can be used with quantum computers. In classical molecular mechanics atomic coordinates are perturbed to achieve a desired state, representing some energy minimum. The reference state is usually encoded in force fields. The first use case, presented in this work, demonstrates an analogous case where two points in 3D (*x, y, z*) coordinates are expressed using spherical coordinates (*r, θ, φ*). To ensure that the subject converges to the reference state, the spherical coordinates (*r, θ, φ*) are directly loaded onto qubit using the UGate. Together with the CSwapGate, the dot product is computed allowing for the direct comparison of the two qubits and indirectly the same for the encoded coordinates and any resulting difference then minimized.

This model and its variations used to compare both vector magnitudes and directions has significant implications in the area of molecular mechanics as it can be tied together with various optimization routines to achieve desired states of molecular systems. This work demonstrates, without quantum advantage, and using empirical systems utility of this basic model to optimize geometries of molecules using a hybrid classical-quantum computing workflow.

Apart from the above model, this work also shows the use of quantum machine learning with biological data. Although the example does not demonstrate quantum advantage, and the classical classifier outperforms the quantum classifier, the example demonstrates the use of classical data from geometry optimization problem-space with a quantum machine learning algorithm. In addition to the above, the same data is used to carry out Monte Carlo simulations.

As stated earlier, encoding classical data for use with quantum algorithms presents a significant challenge in this area. This work presents some ideas for encoding data that can further help with the development of quantum algorithms and eventually achieving quantum advantage.

## Conclusion

Using conventional examples of molecular geometry optimization, this work acts as a primer to familiarize the wider community working in the area of molecular mechanics with quantum computing. While quantum advantage is not demonstrated in this work, it successfully demonstrates encoding of classical molecular data for use with quantum computers. By presenting several models working in different ways, this work is expected to draw attention from the wider community and hopefully future work will demonstrate quantum advantage which in turn will benefit the area of molecular mechanics and consequently drug discovery.

## Data and Software Availability

Python notebooks and the data used in this work can be made available on request.

