## Supplementary figures and images for "On quantum computing and geometry optimization"

### Profile_4.pdf

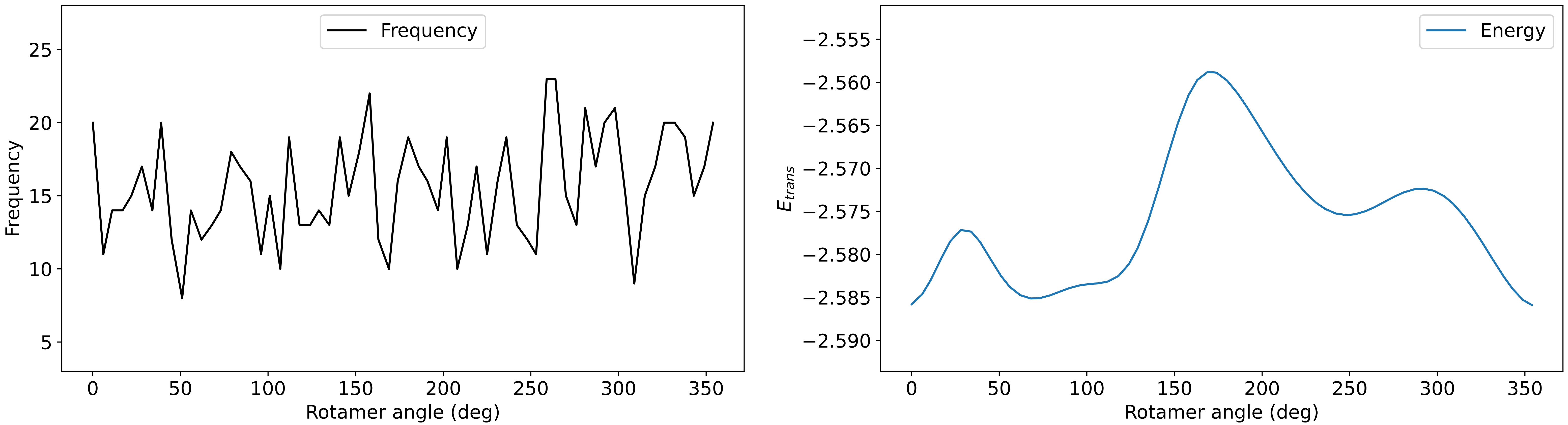

### Profile_5.pdf

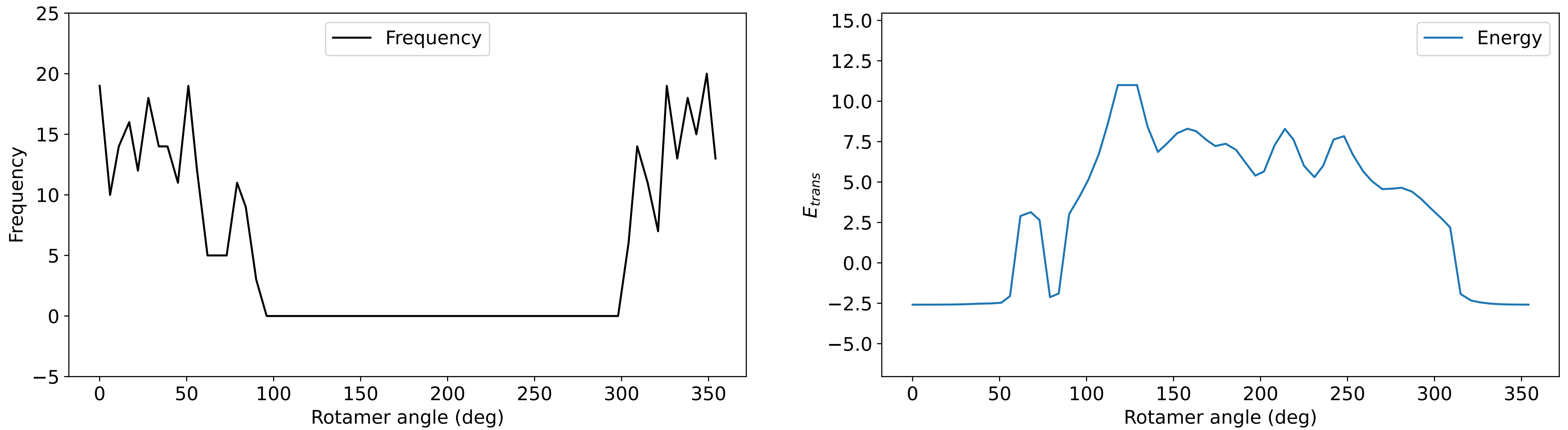

### Profile_6.pdf

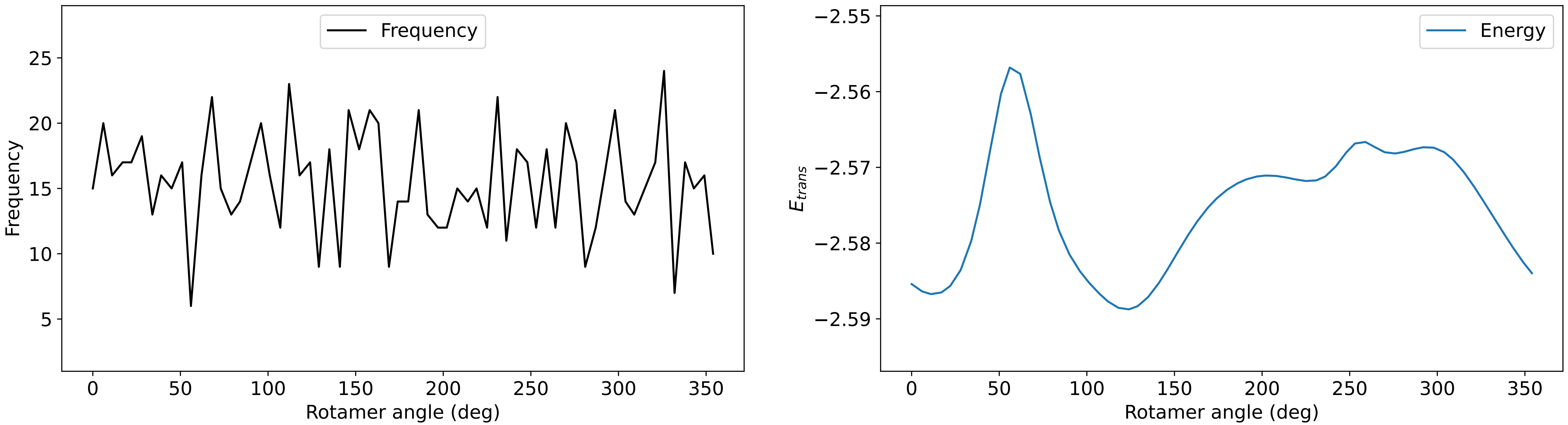

### Profile_7.pdf

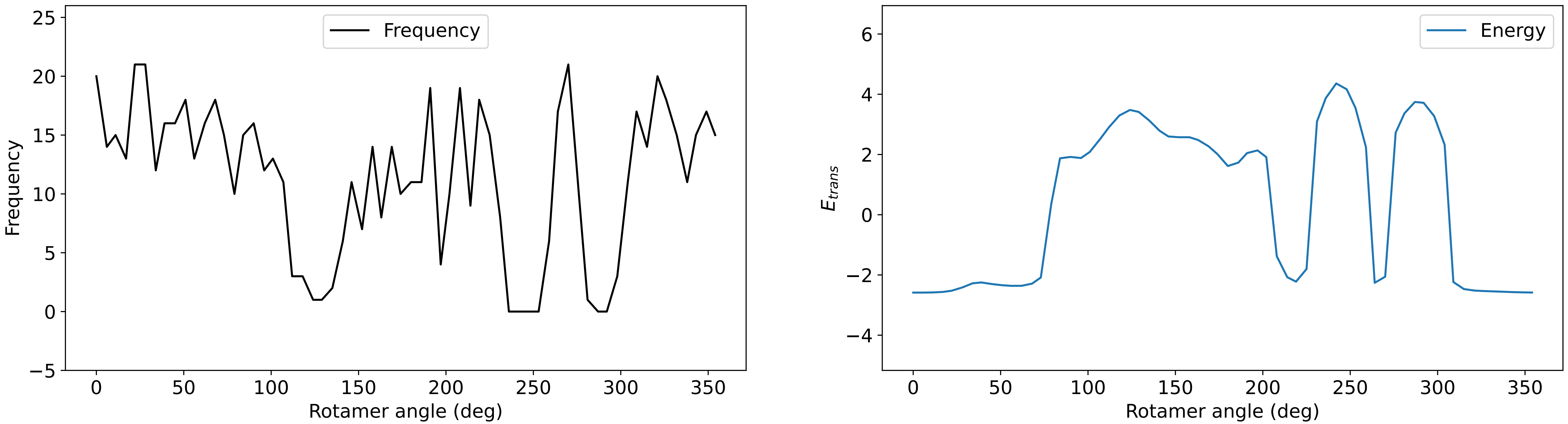

### Profile_8.pdf

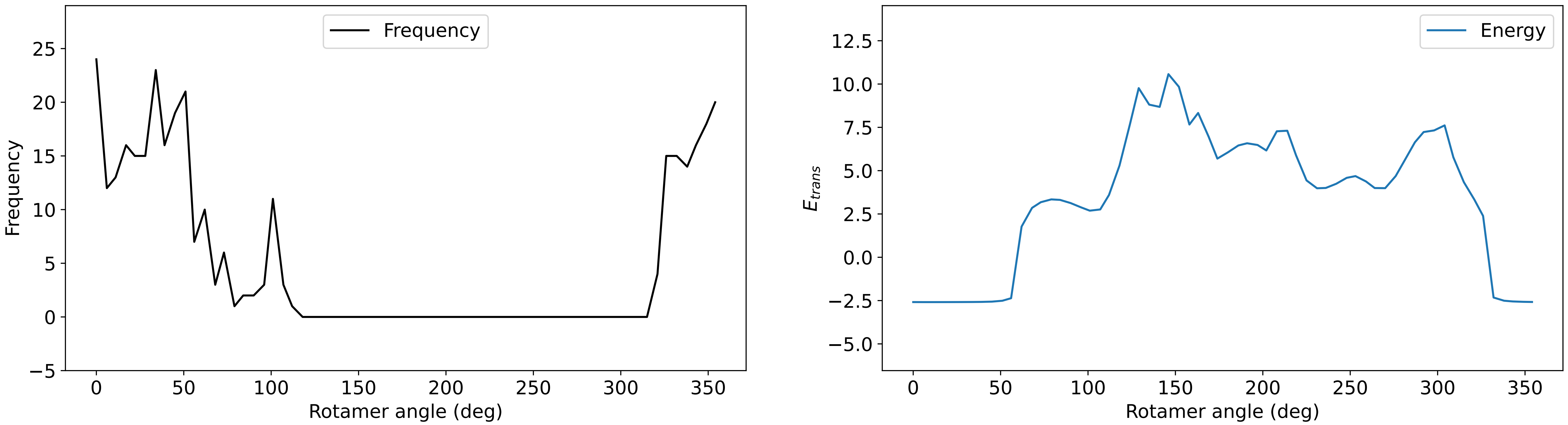

### Profile_9.pdf

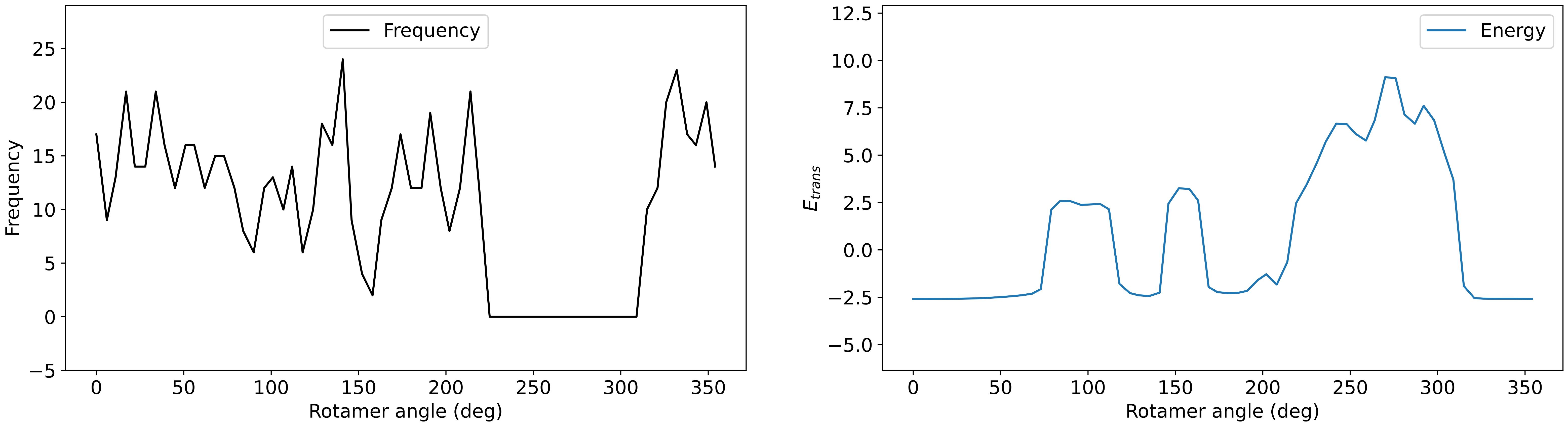

### Profile_10.pdf

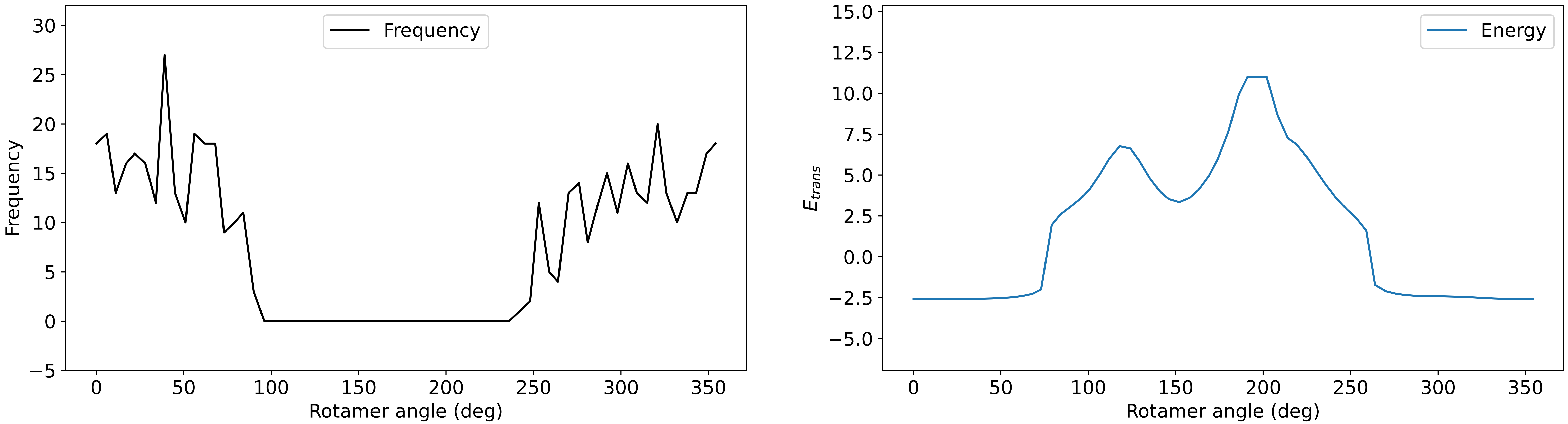

### Profile_11.pdf

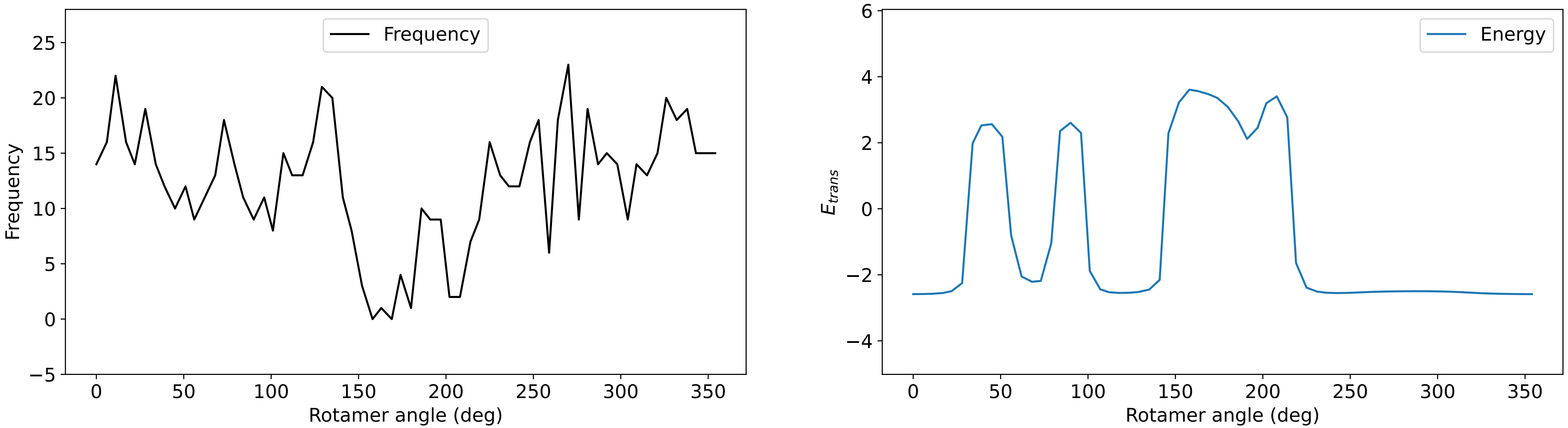

### Profile_12.pdf

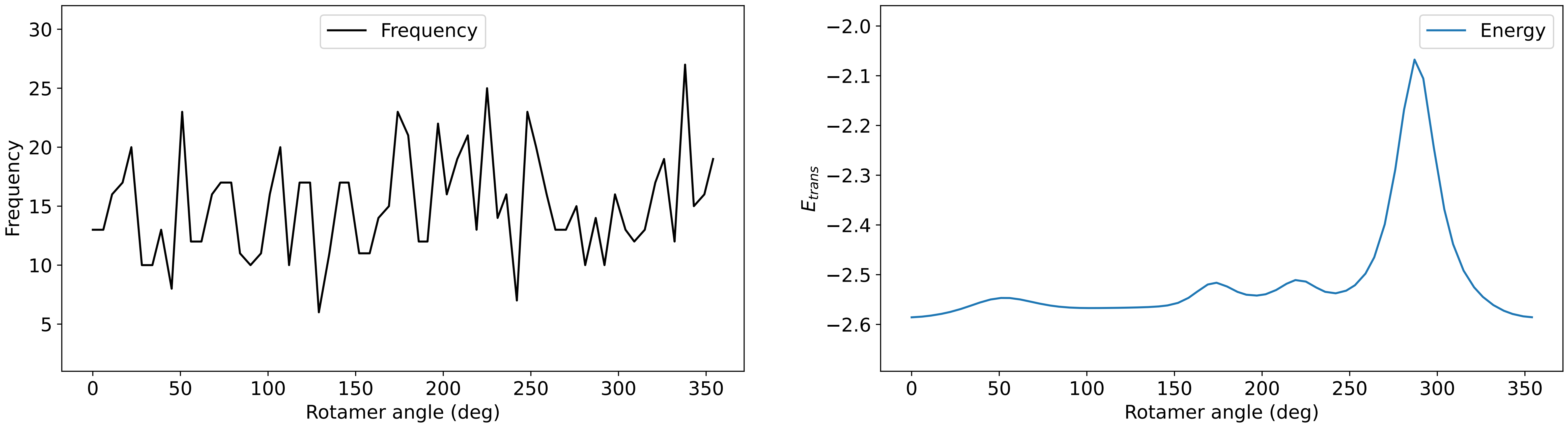

### Profile_13.pdf

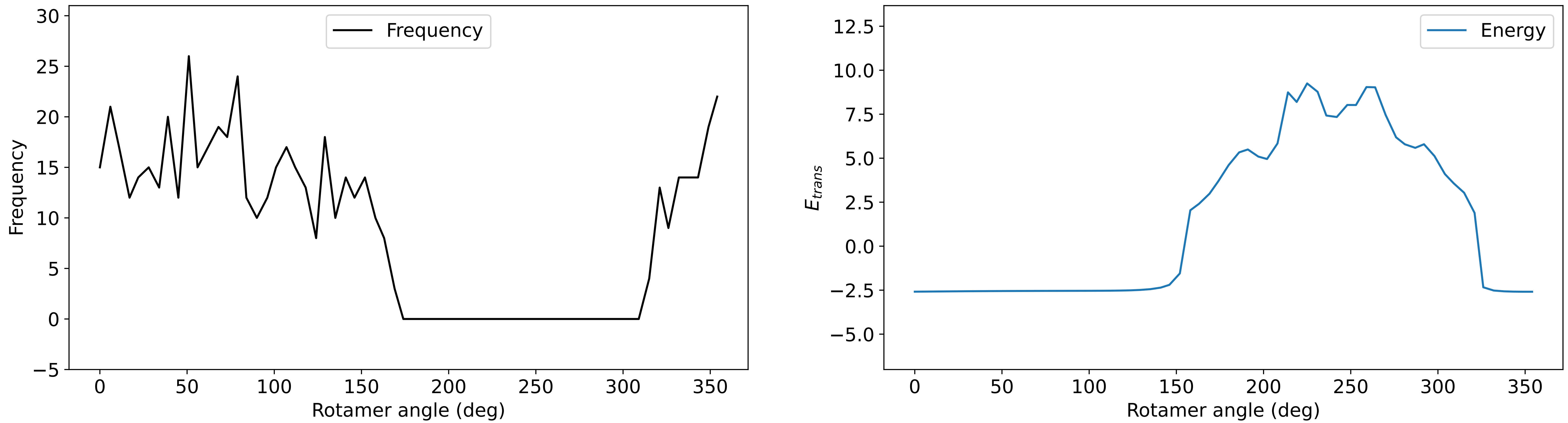

### Profile_15.pdf

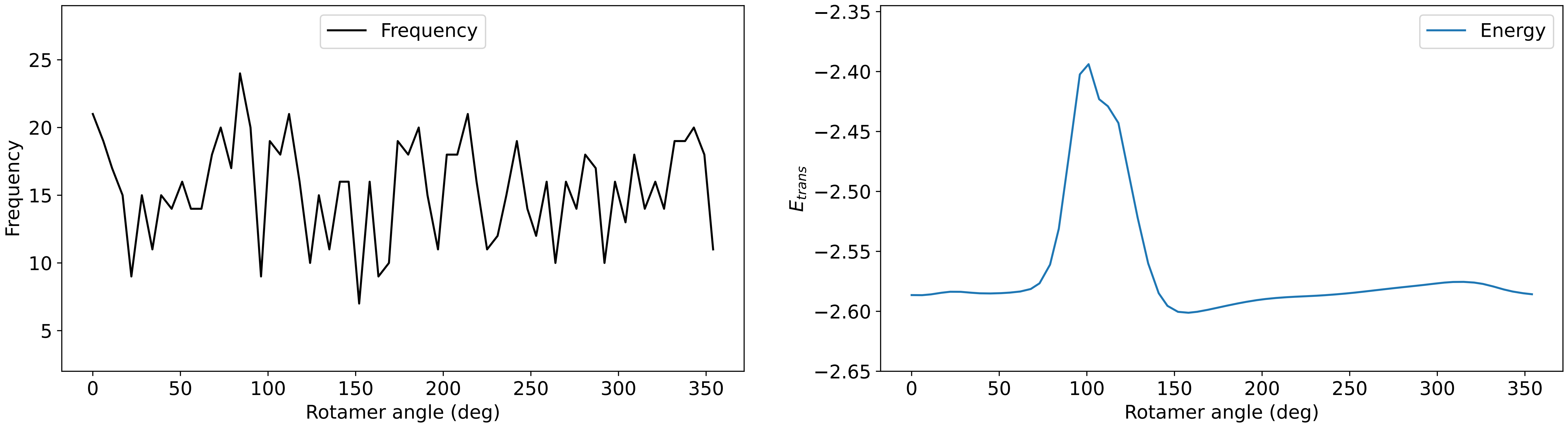

### Profile_16.pdf

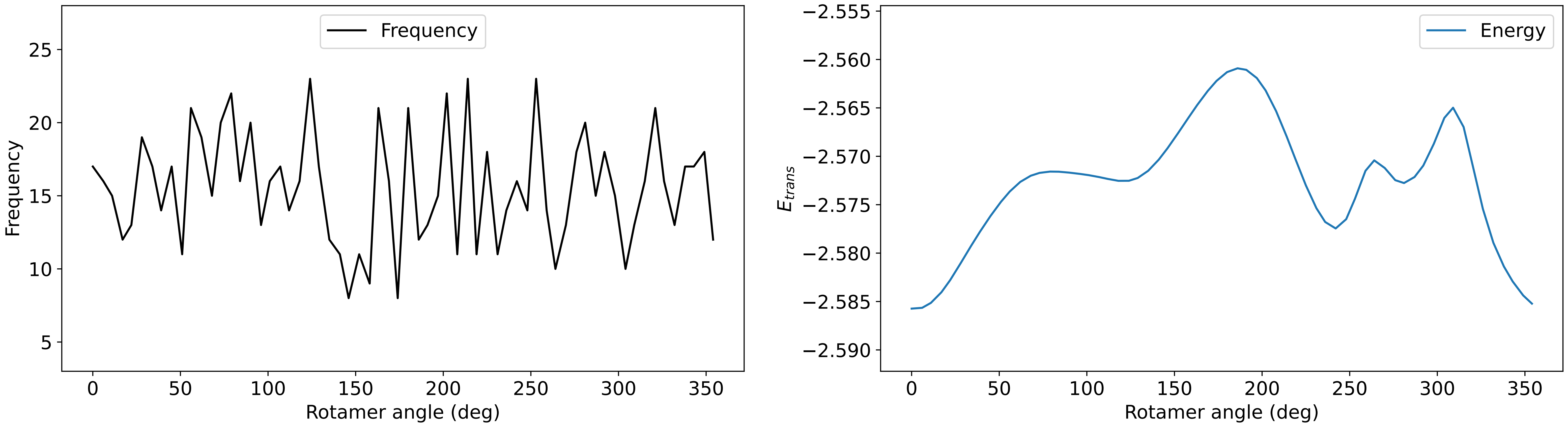

### Profile_17.pdf

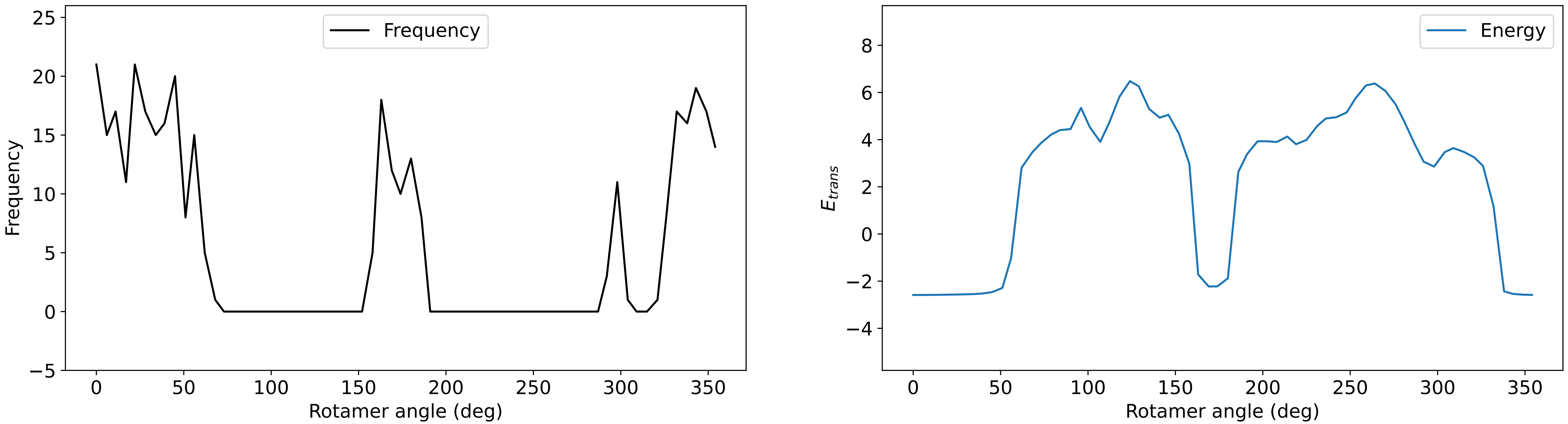

### Profile_18.pdf

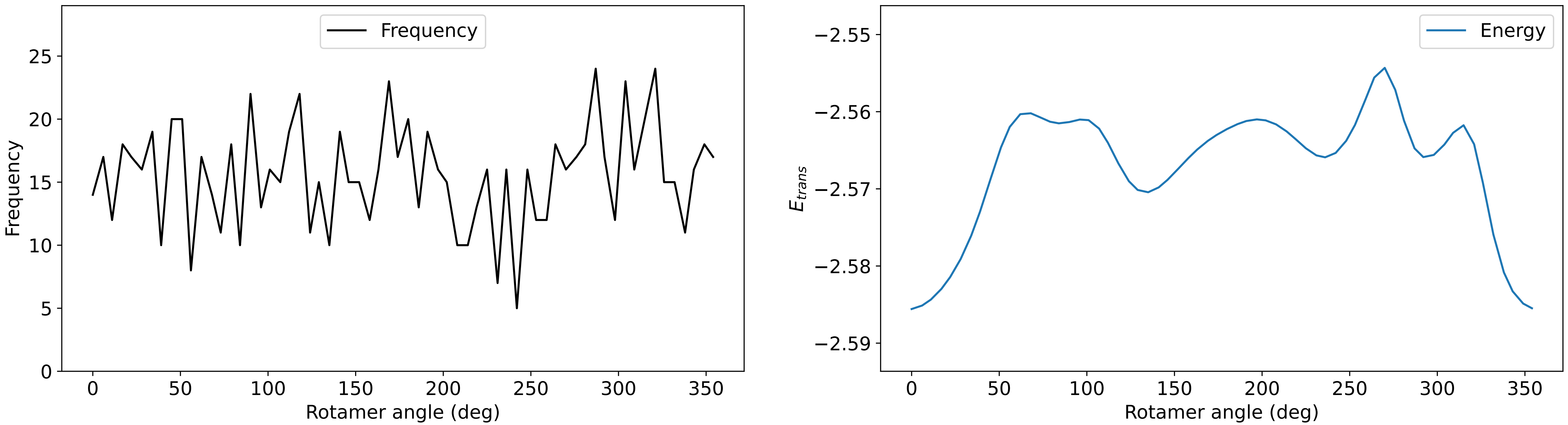

### Profile_19.pdf

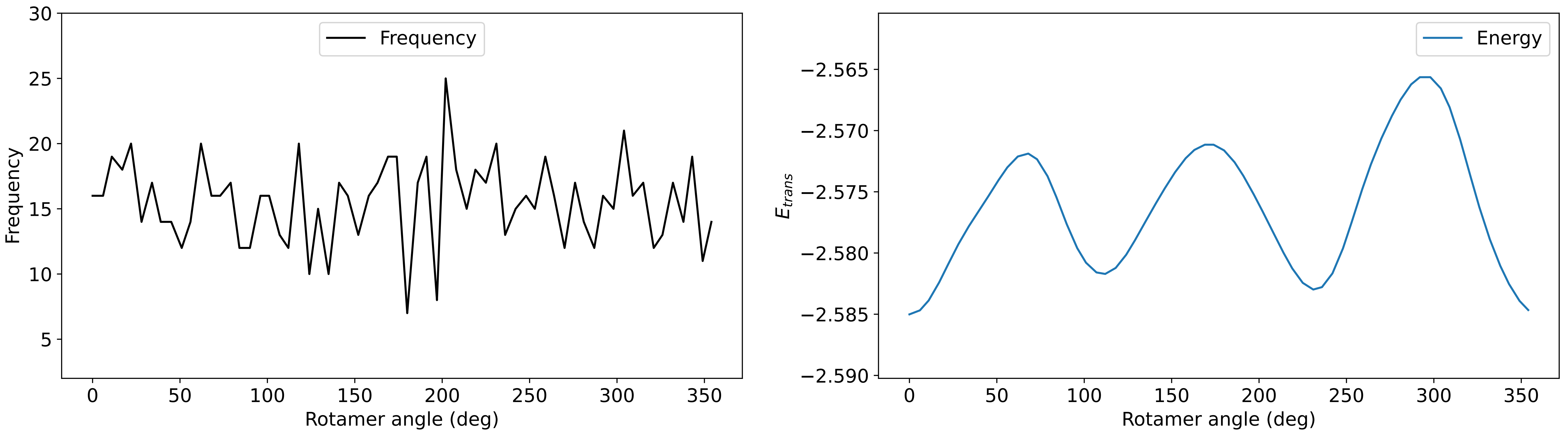

### Profile_20.pdf

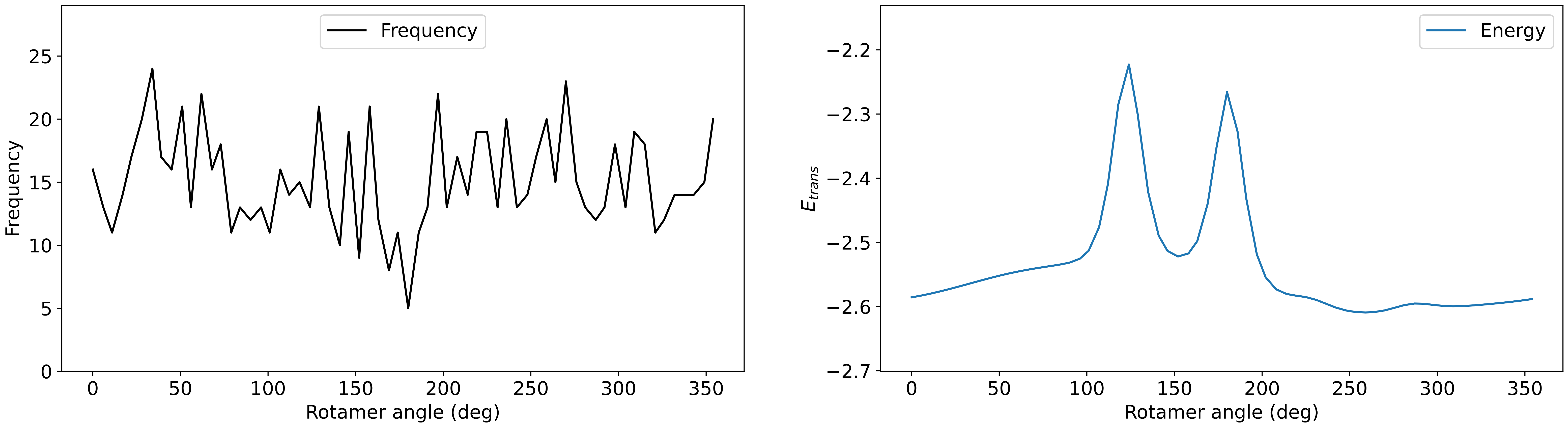

### Profile_21.pdf

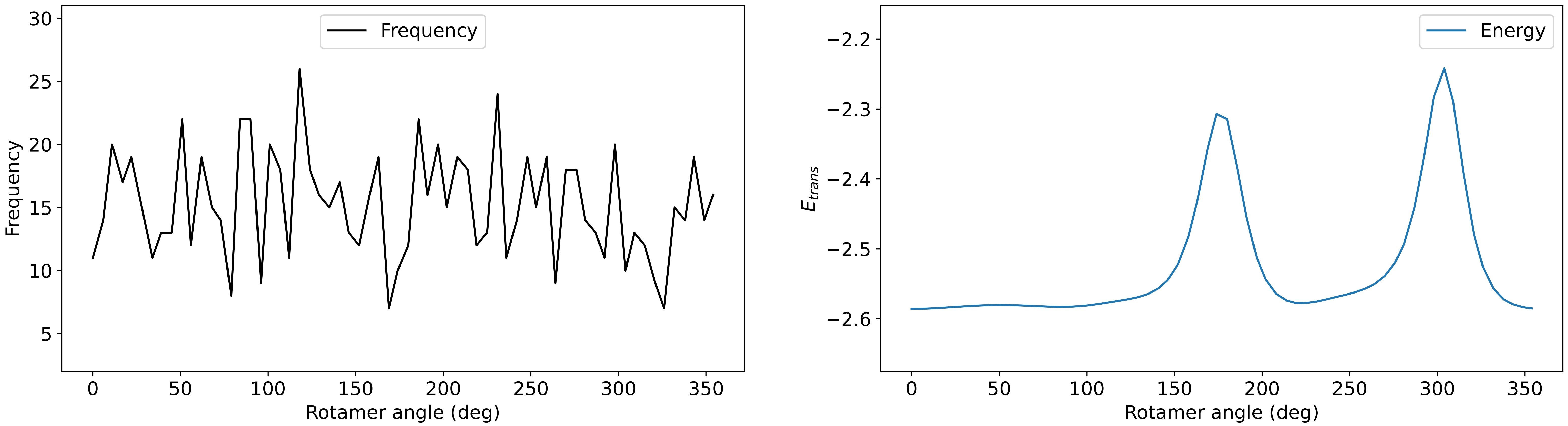

### Profile_22.pdf

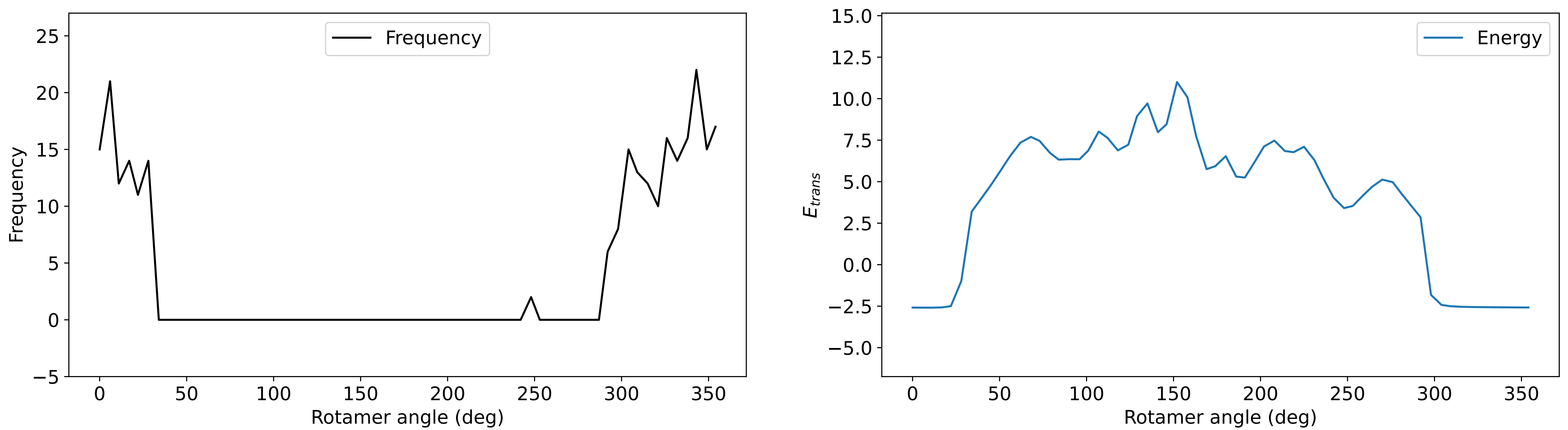

### Profile_24.pdf

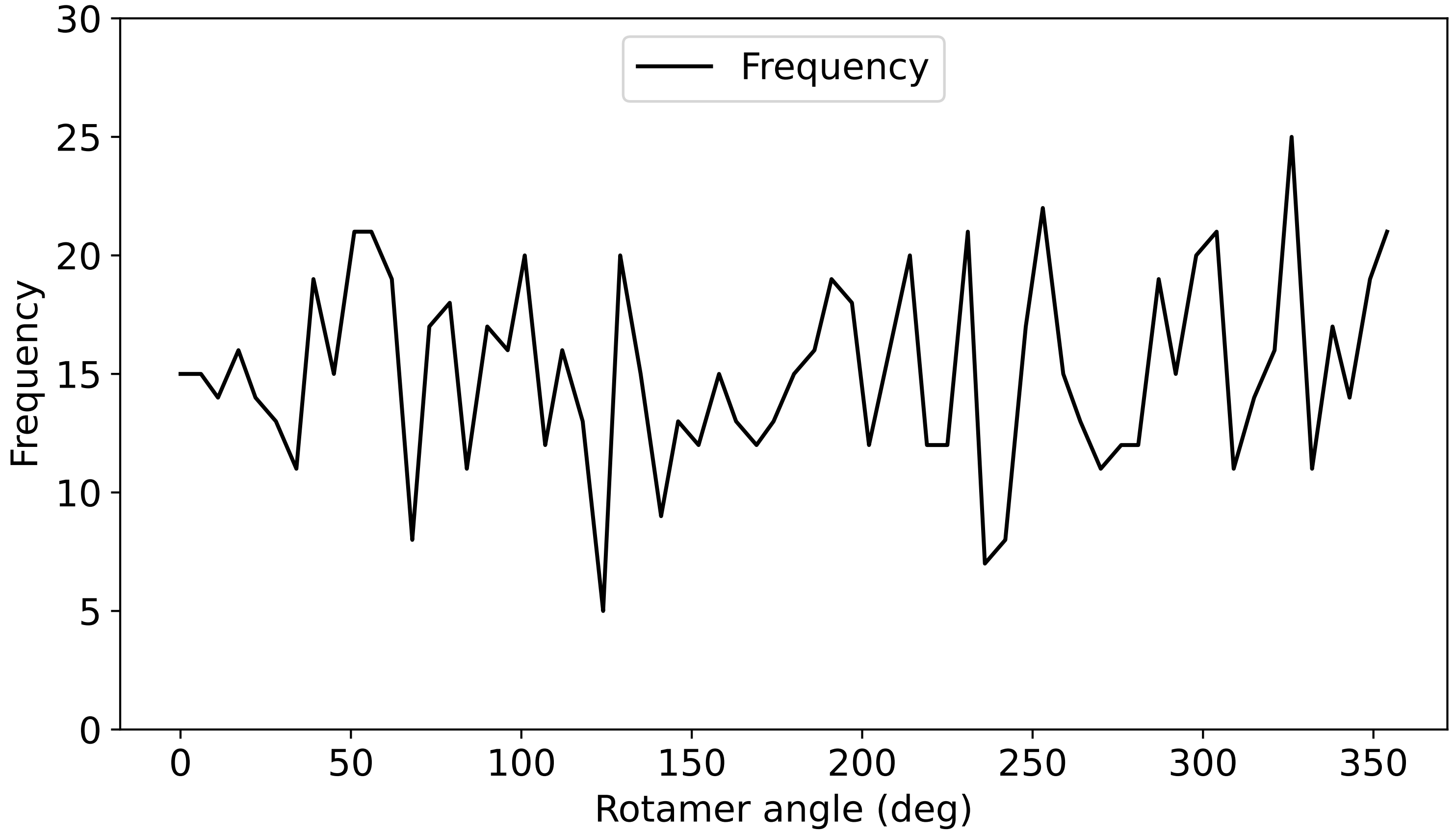

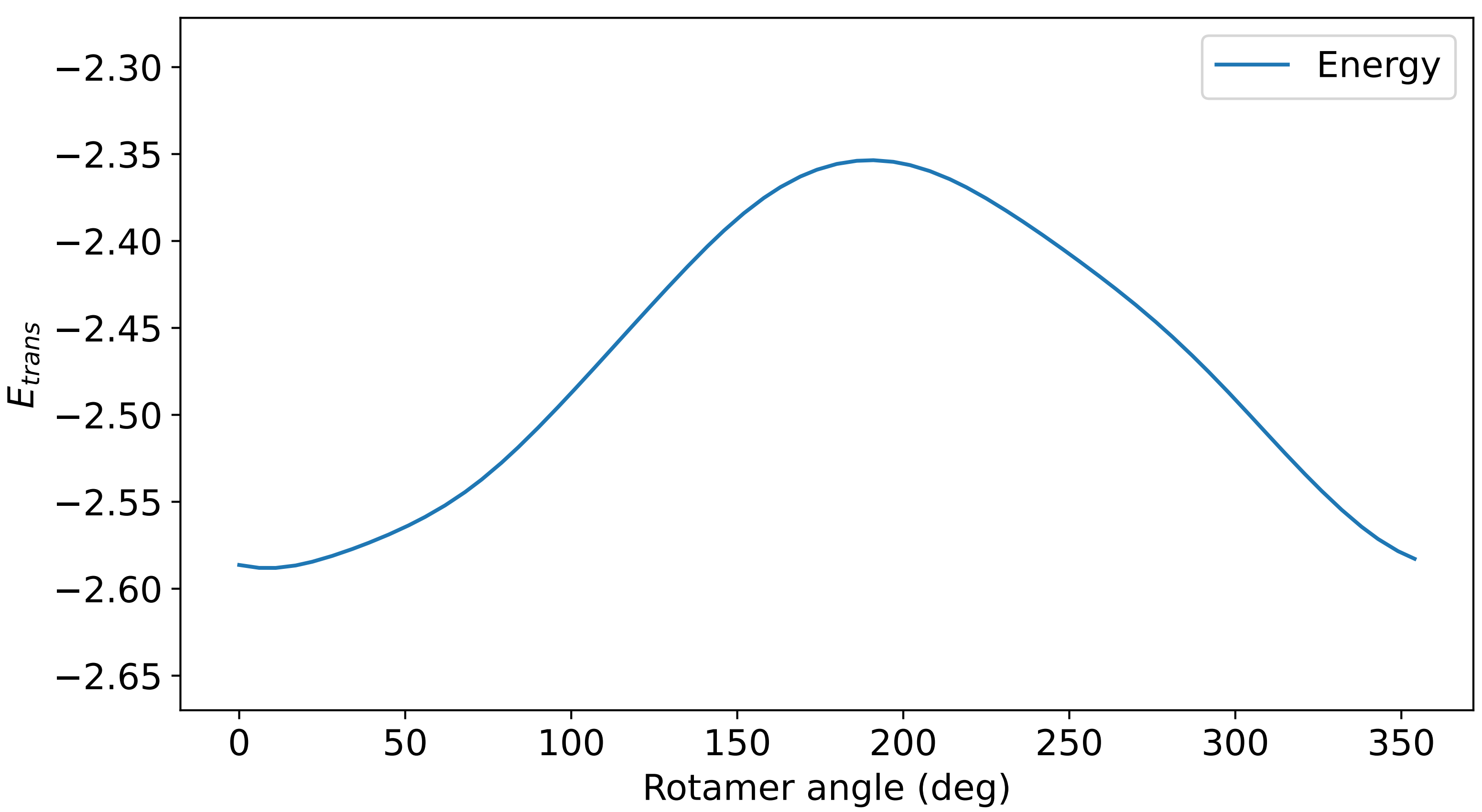

### Profile_25.pdf

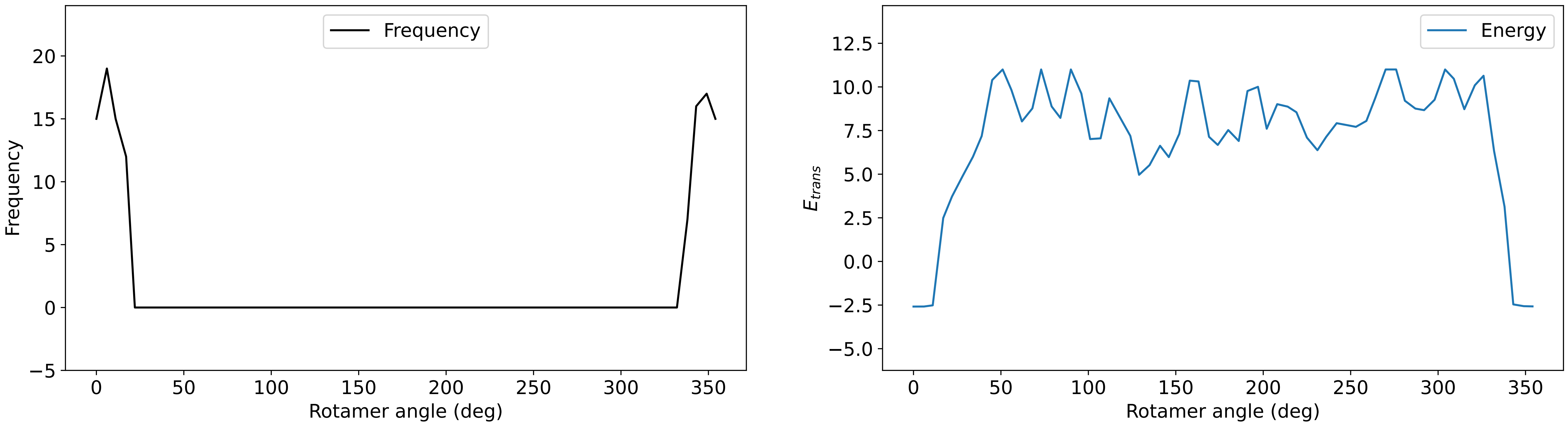

### Profile_26.pdf

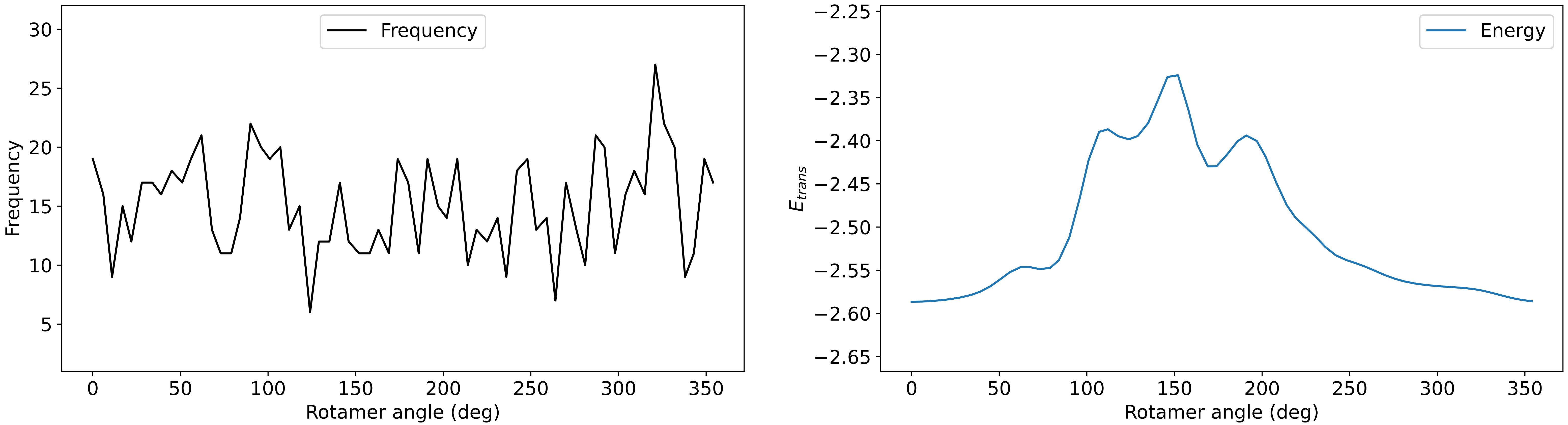

### Profile_27.pdf

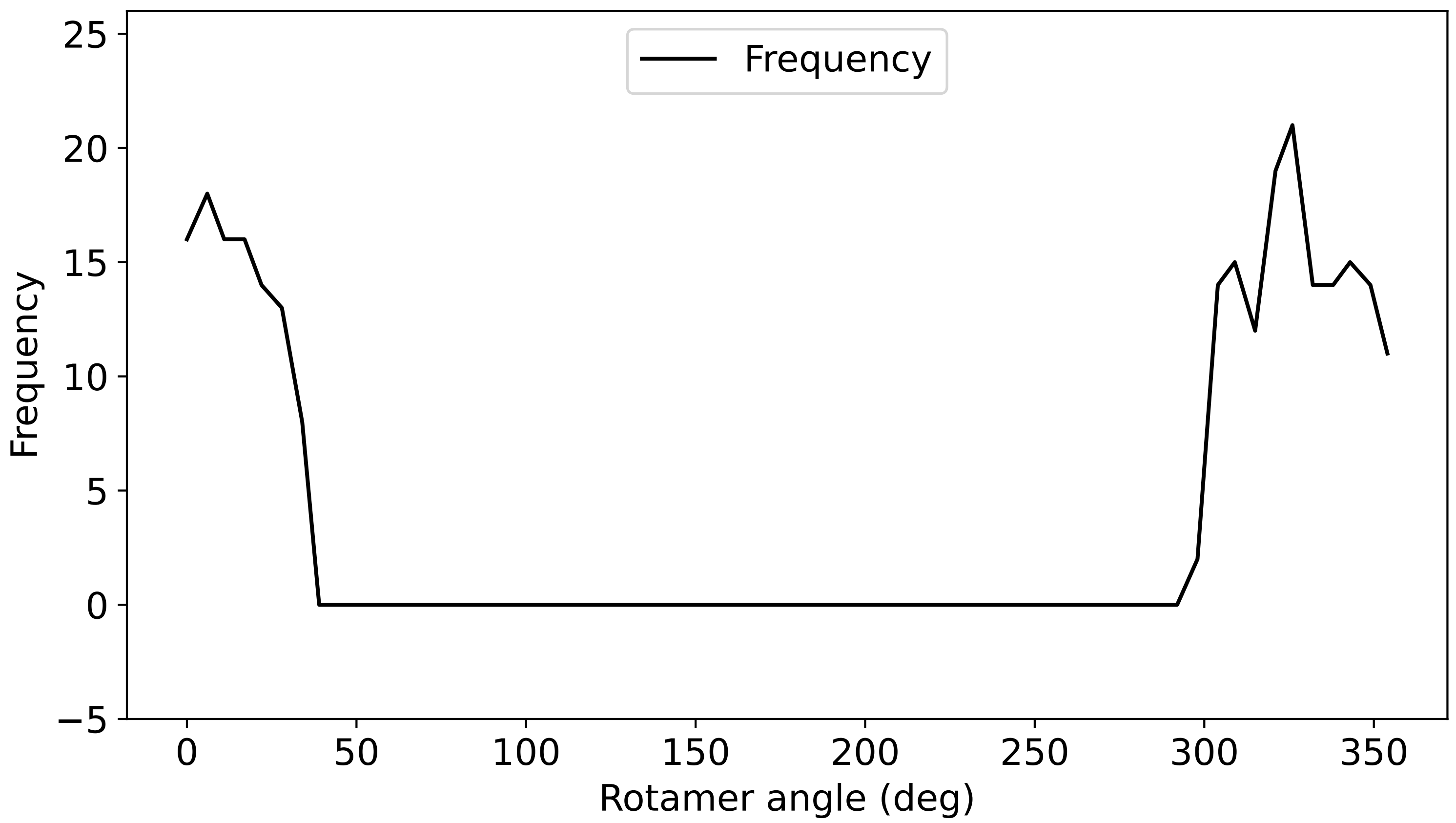

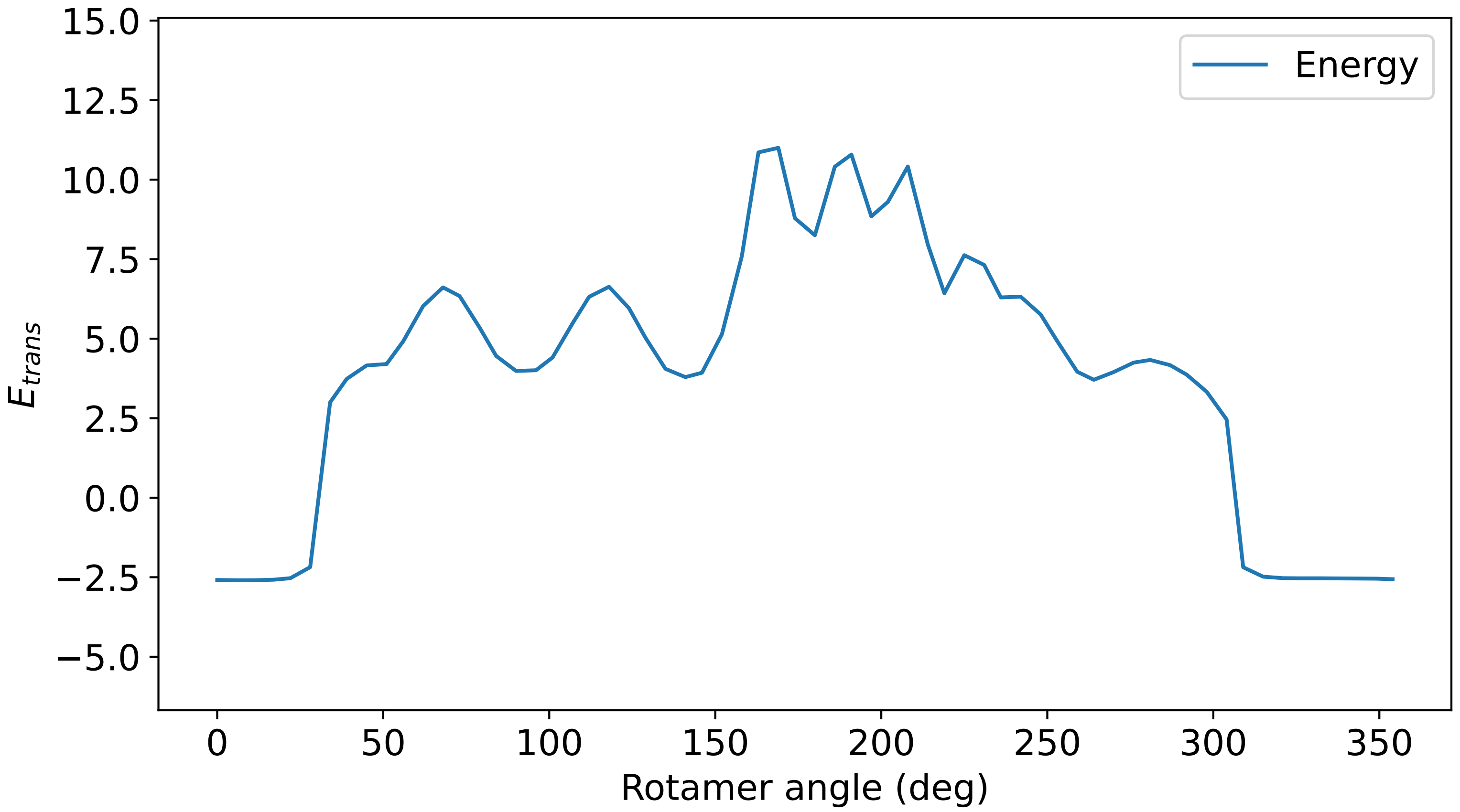

### Profile_28.pdf

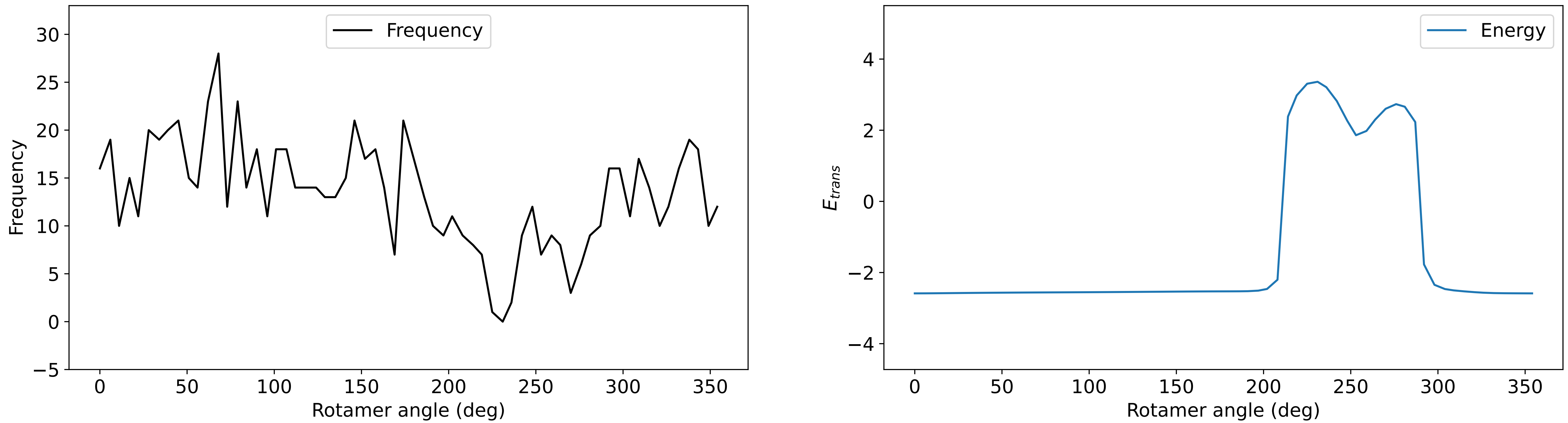

### Profile_29.pdf

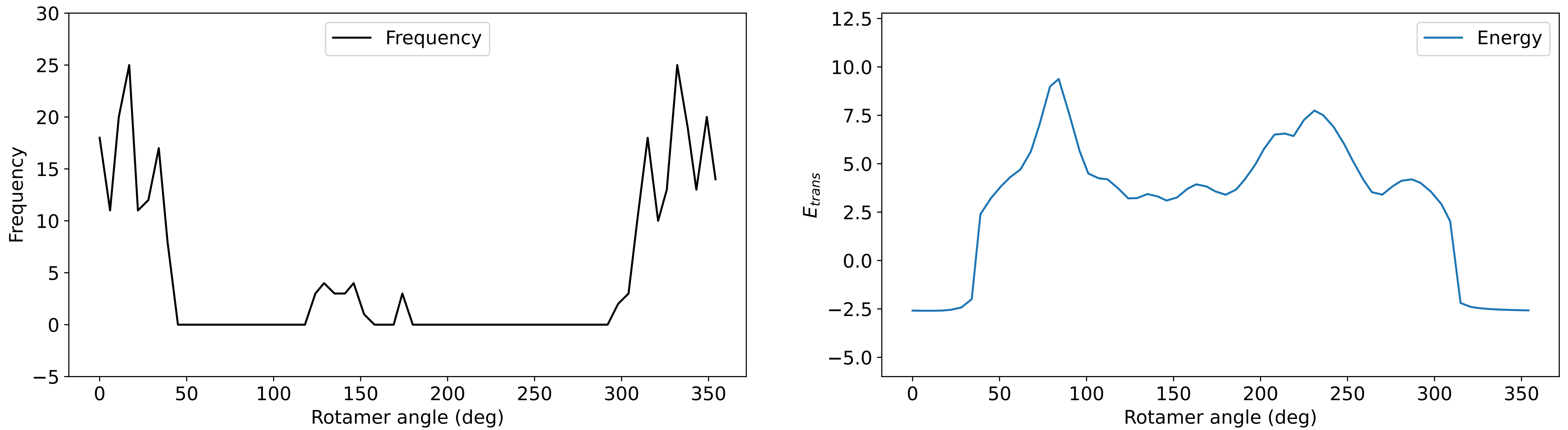

### Profile_30.pdf

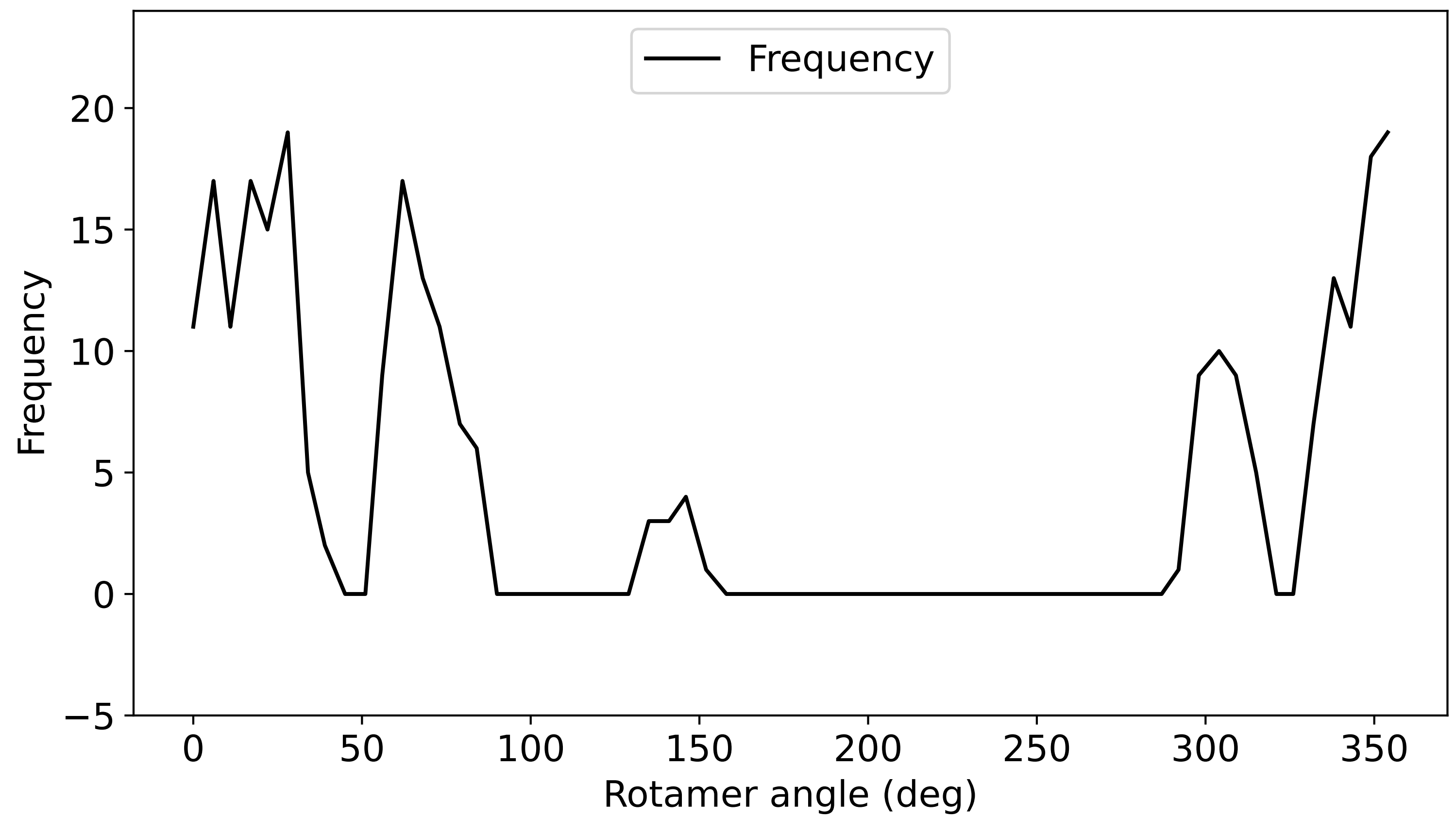

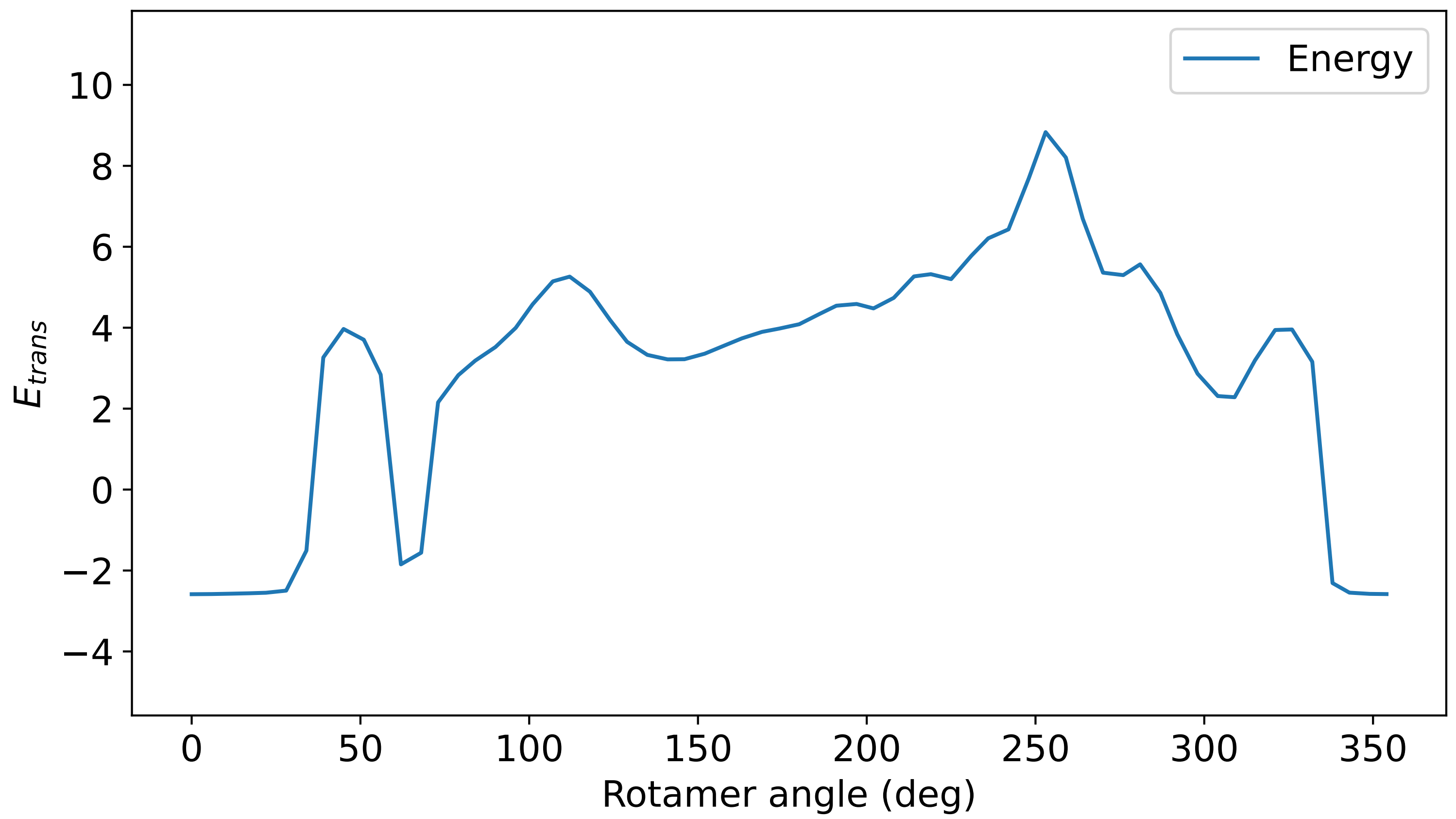

### Profile_31.pdf

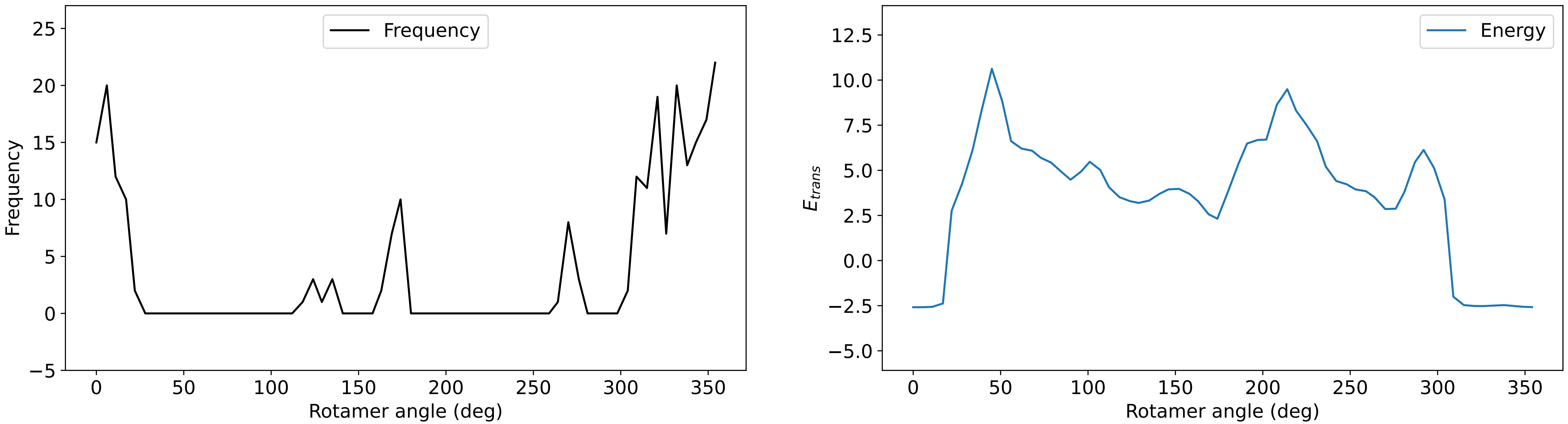

### Profile_32.pdf

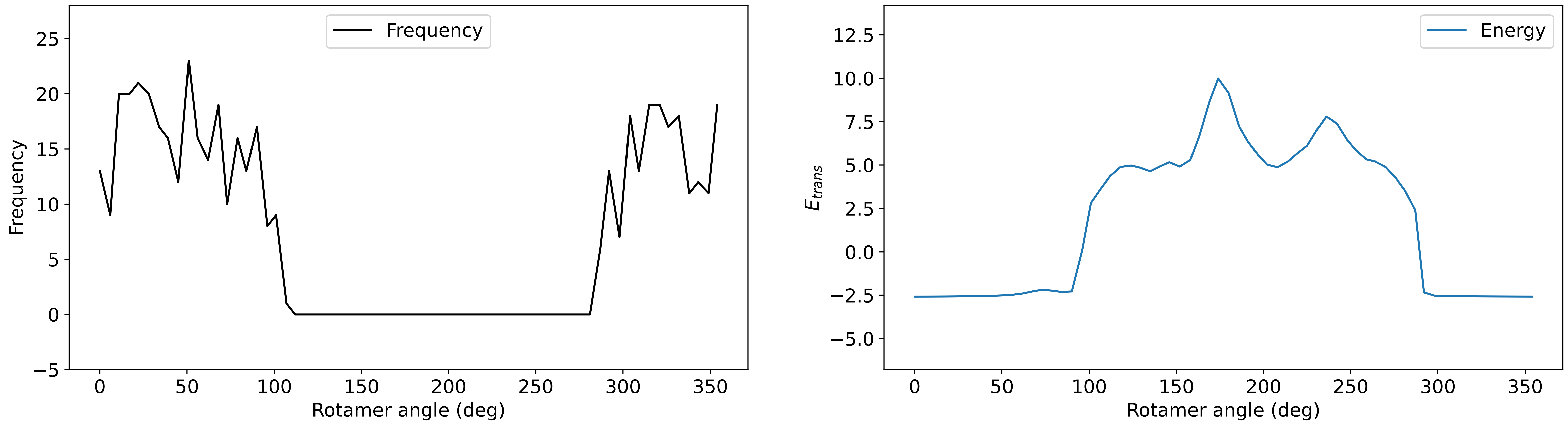

### Profile_33.pdf

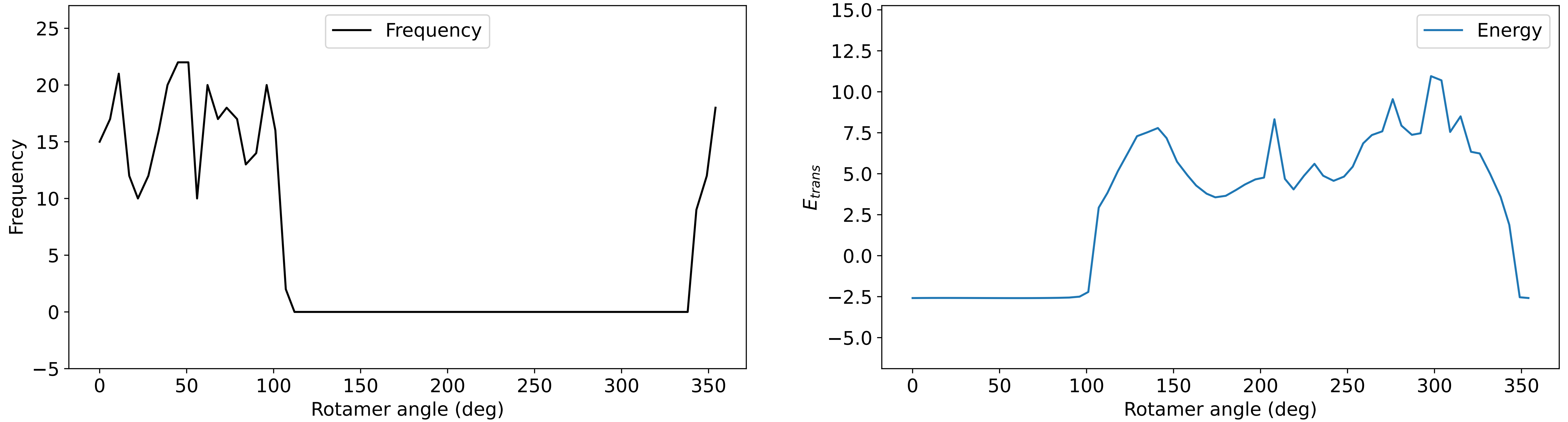

### Profile_34.pdf

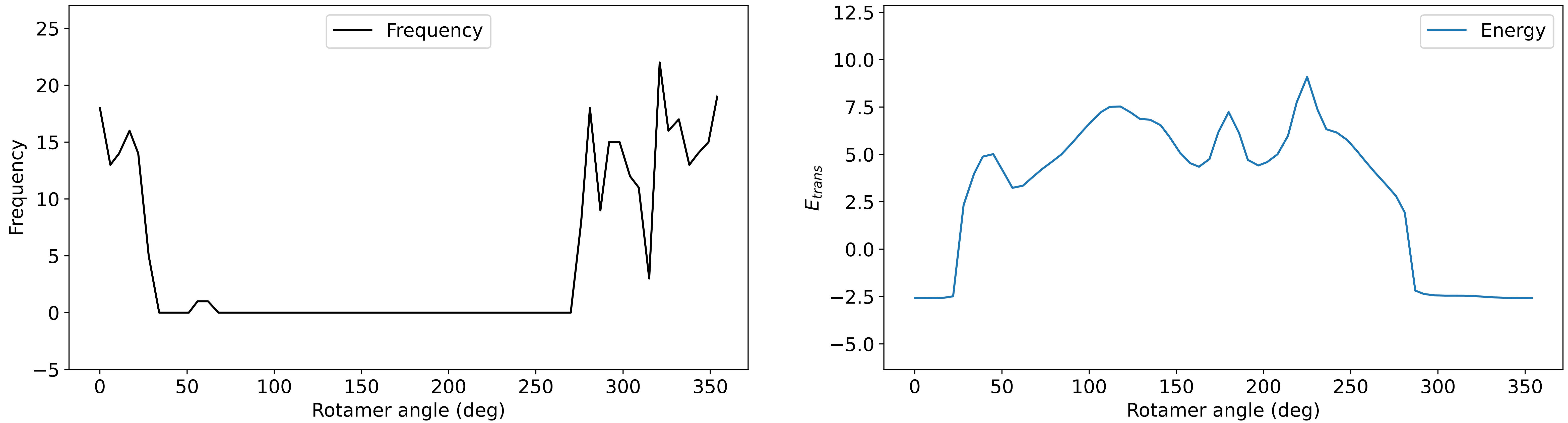

### Profile_35.pdf

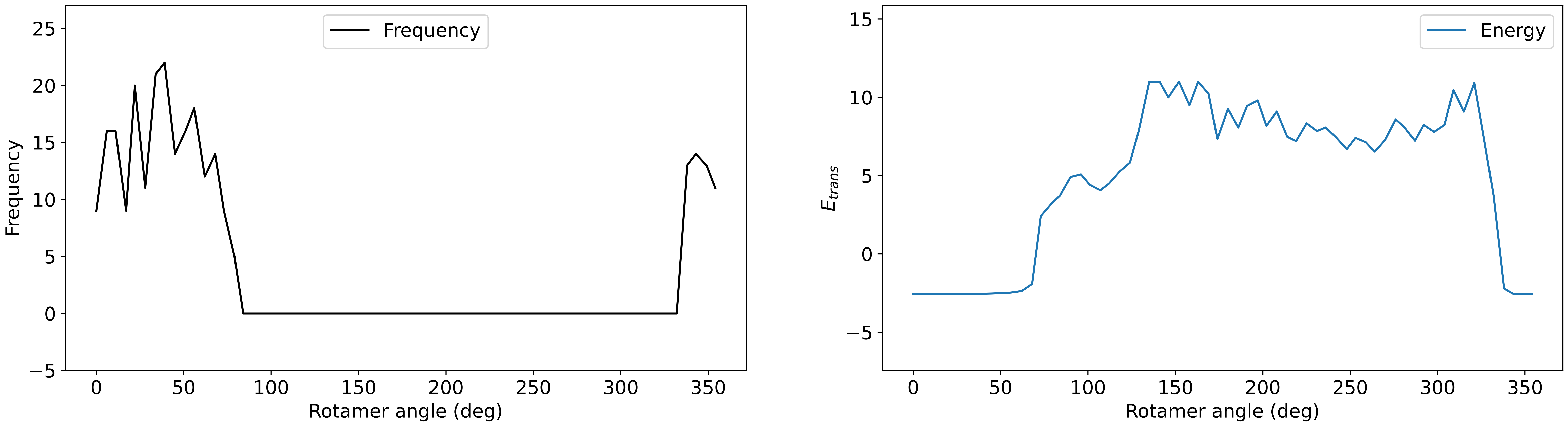

### Profile_36.pdf

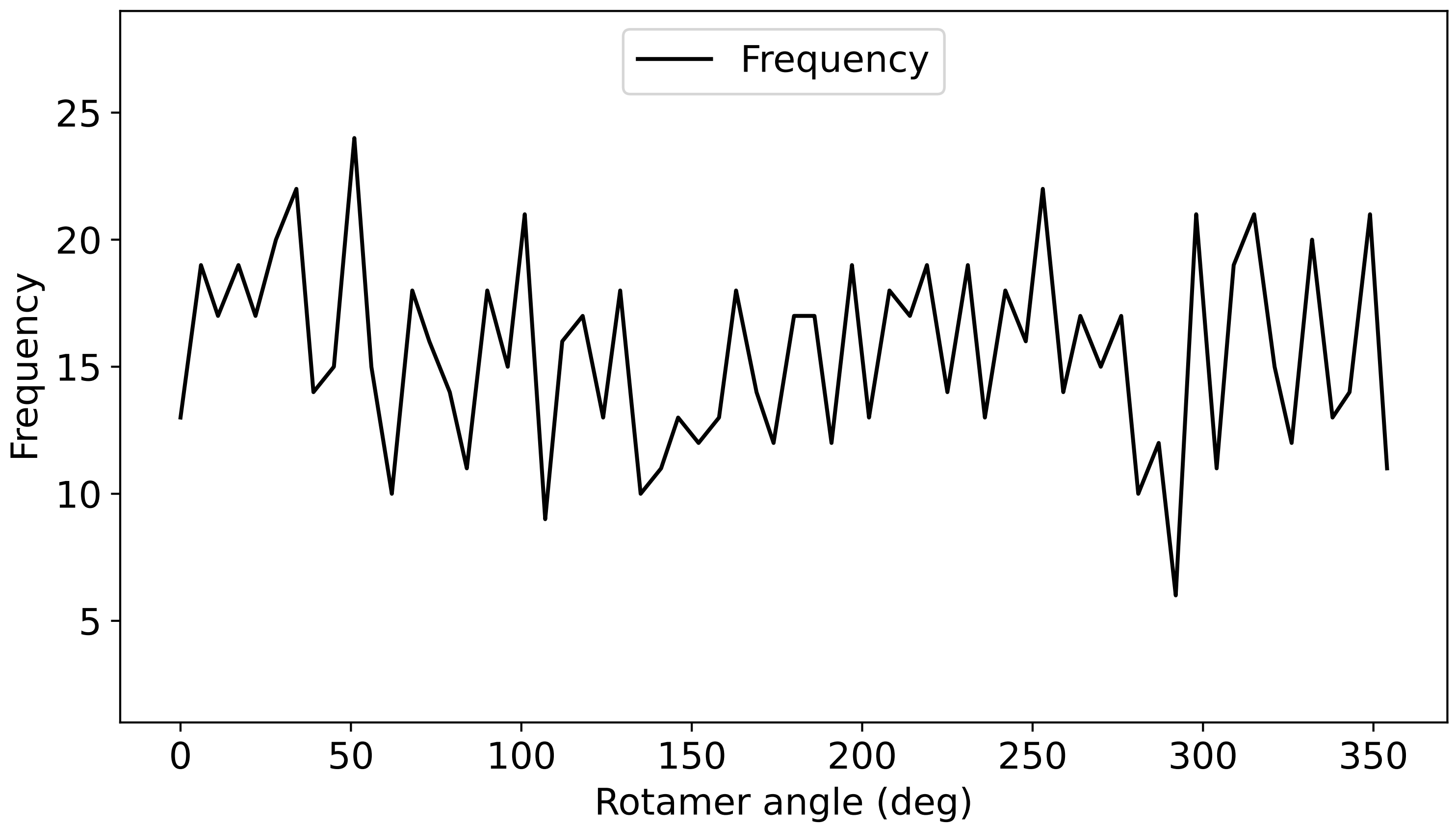

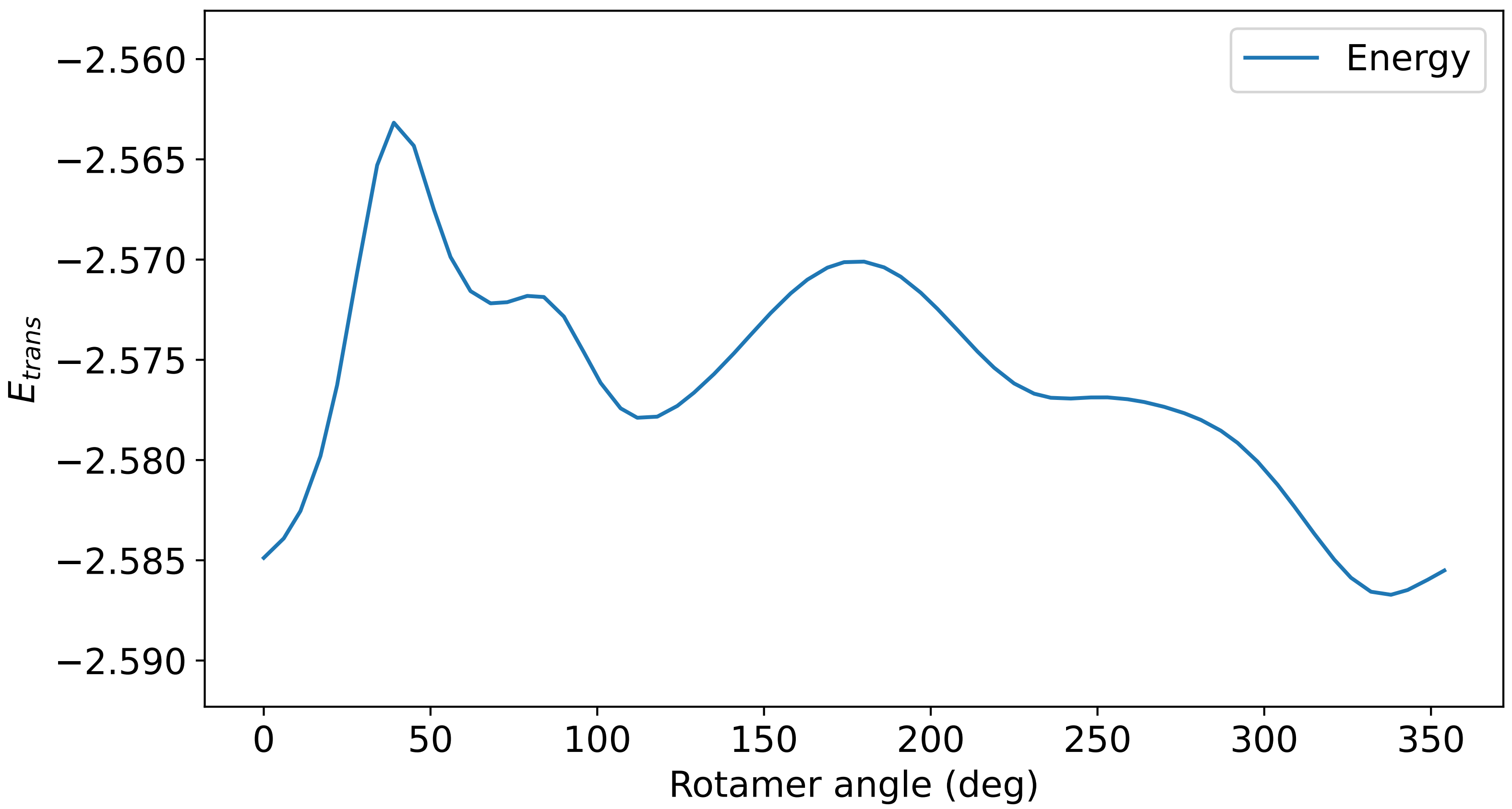

### Profile_37.pdf

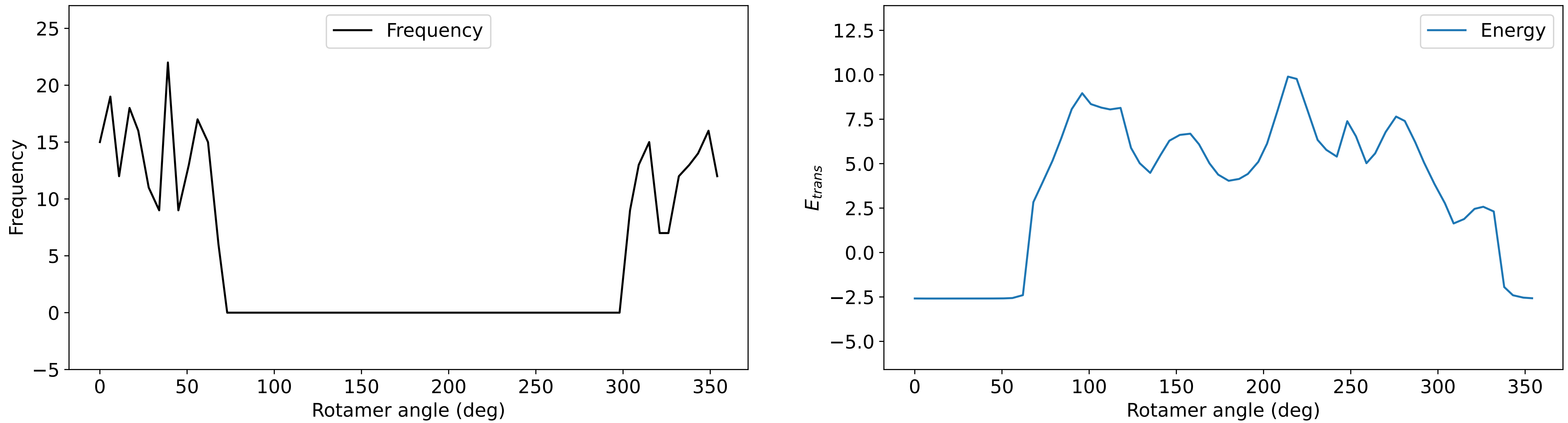

### Profile_38.pdf

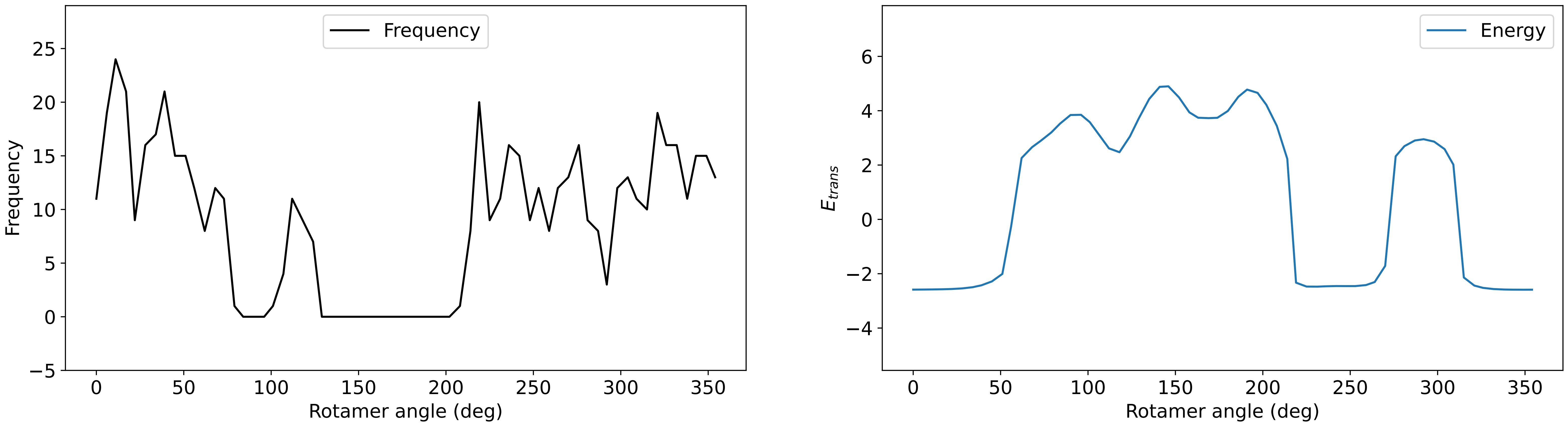

### Profile_40.pdf

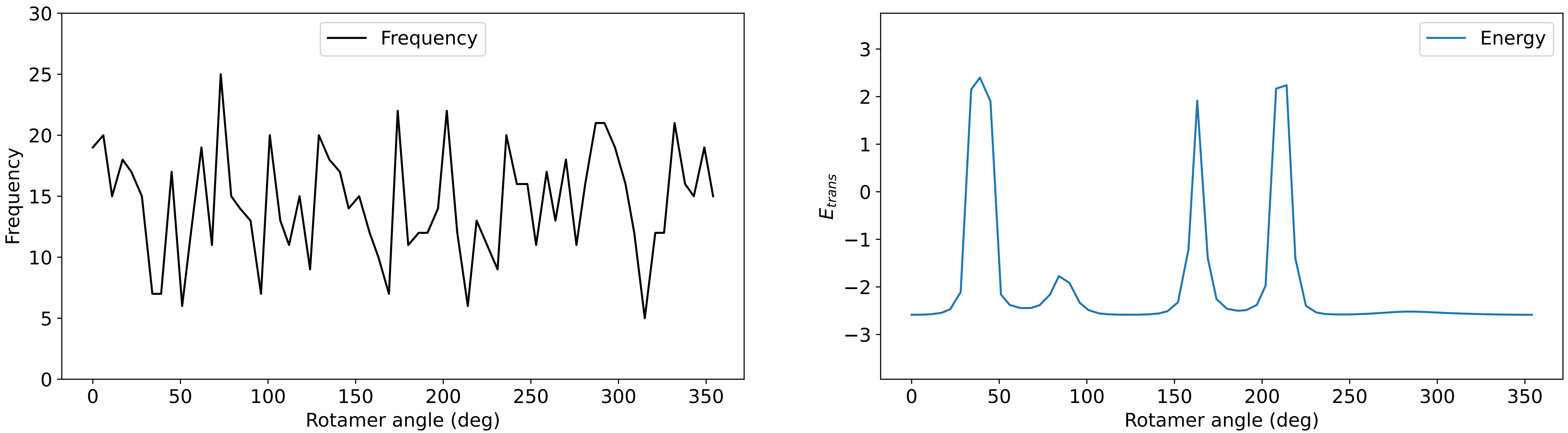

### Profile_43.pdf

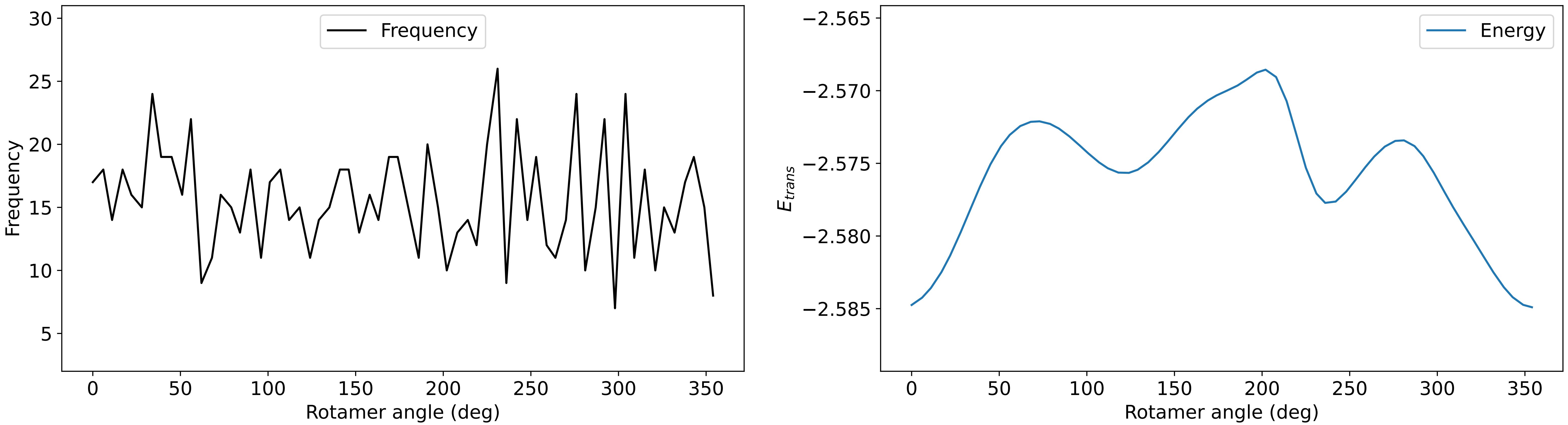

### Profile_44.pdf

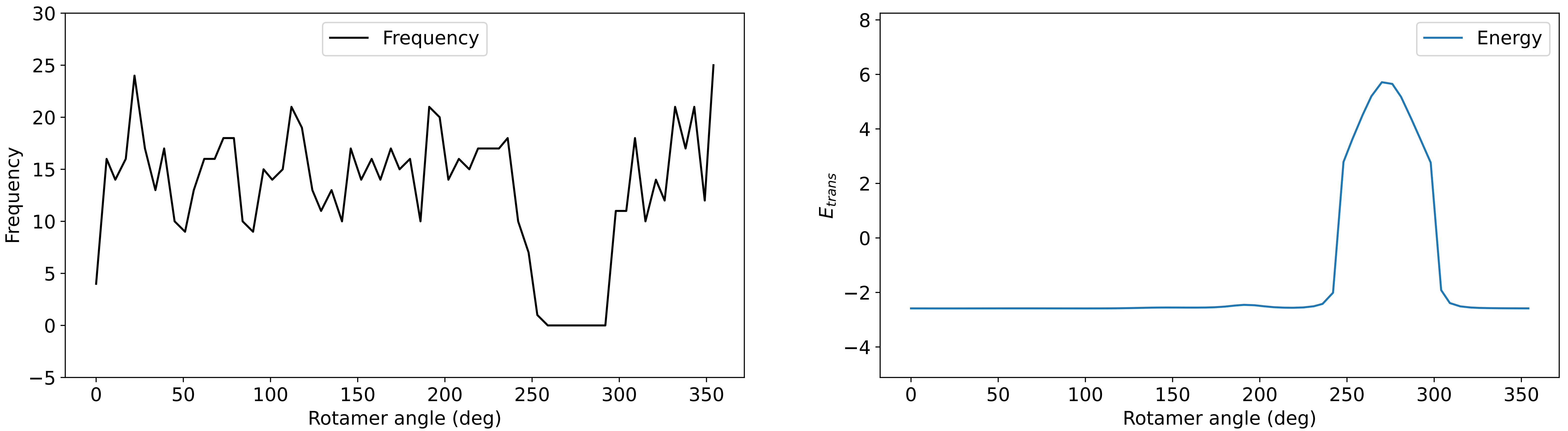

### Profile_45.pdf

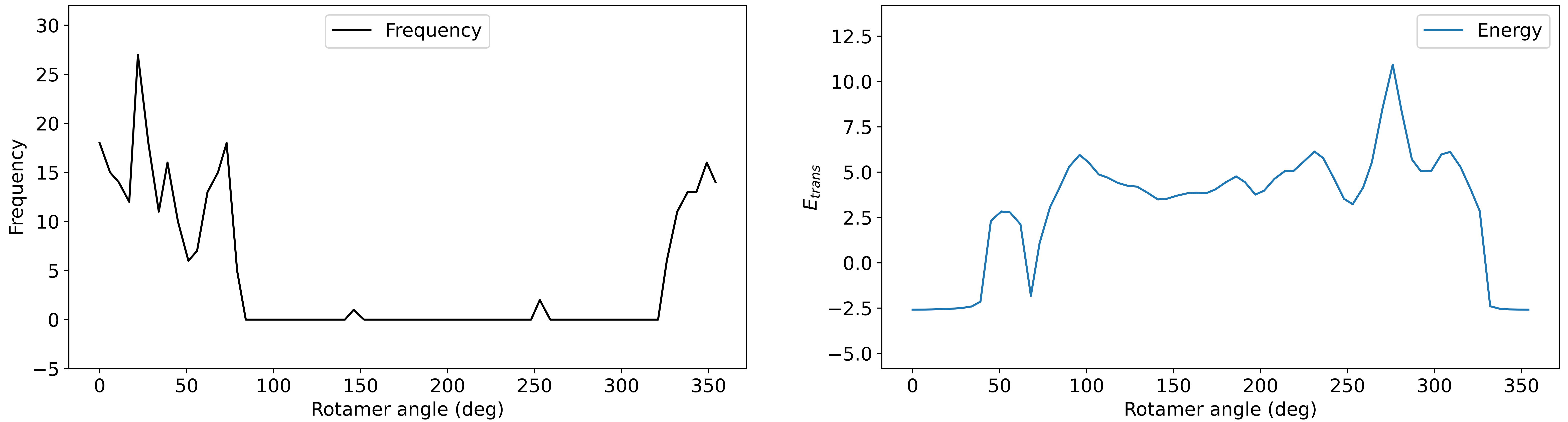

### Profile_46.pdf

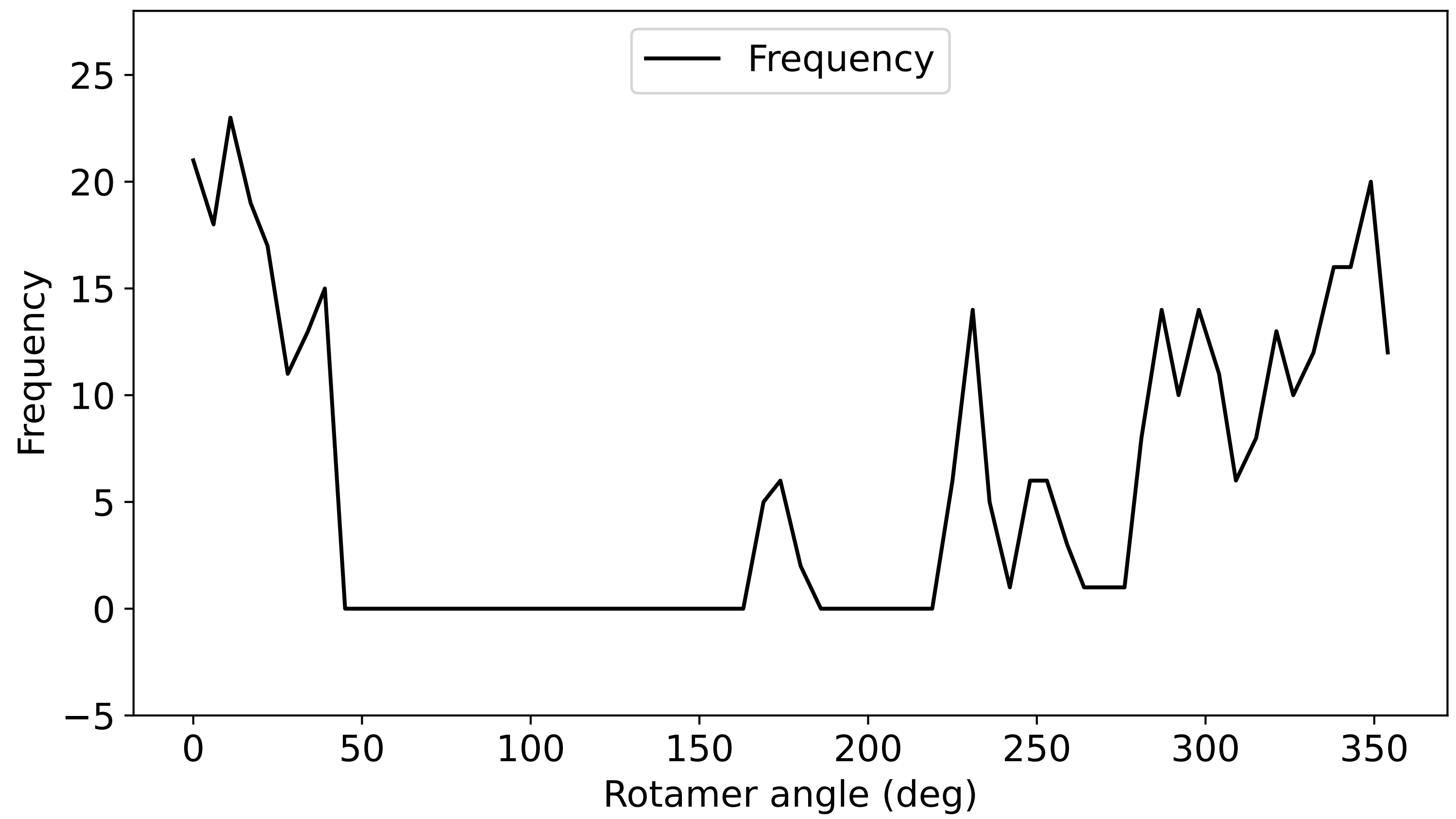

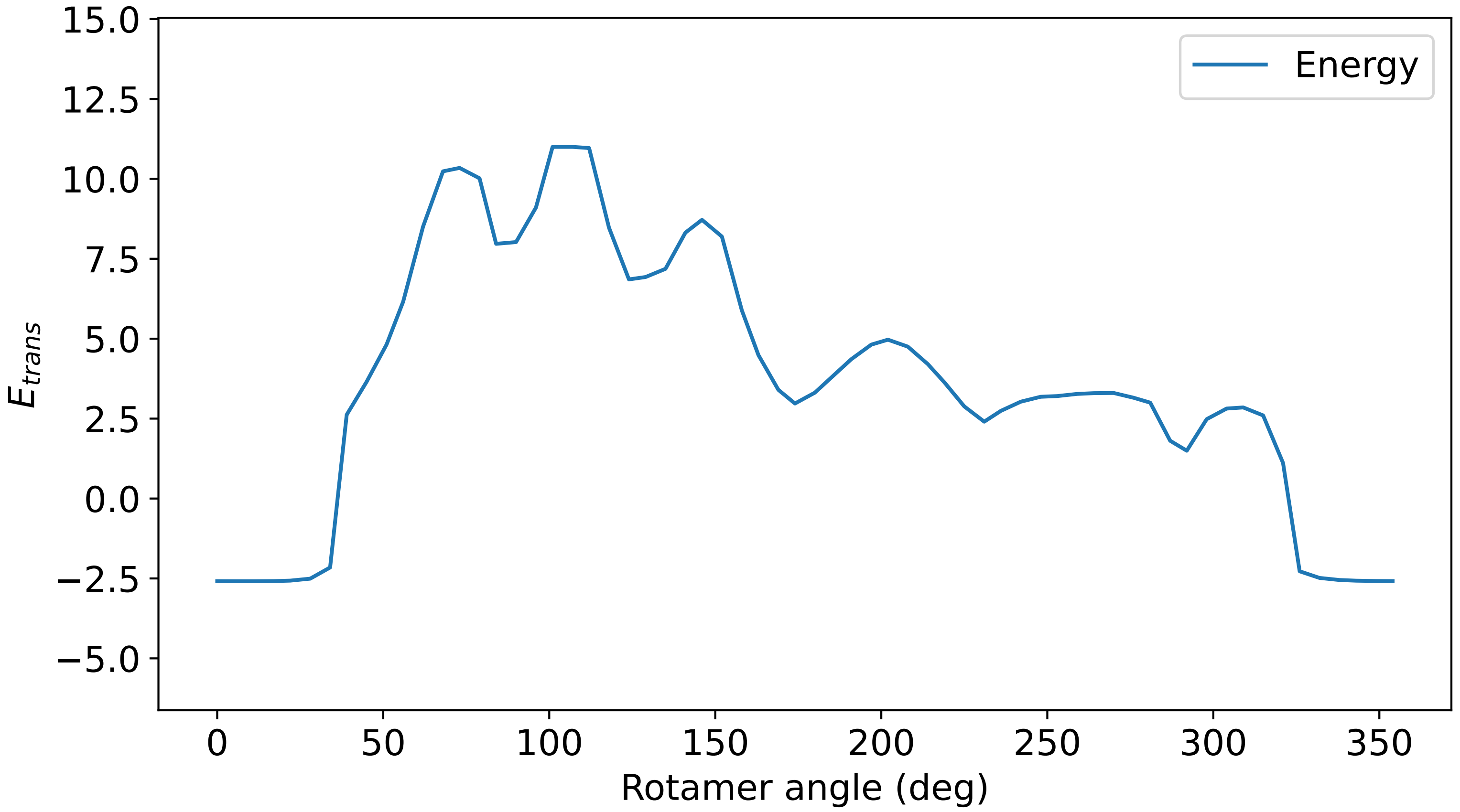

### Profile_47.pdf

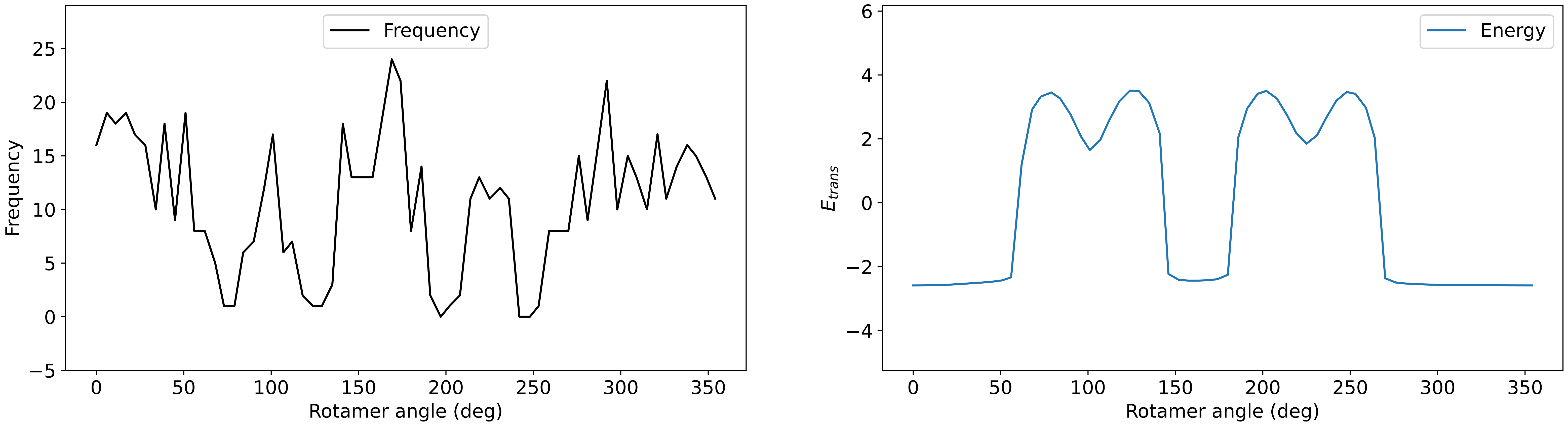

### Profile_48.pdf

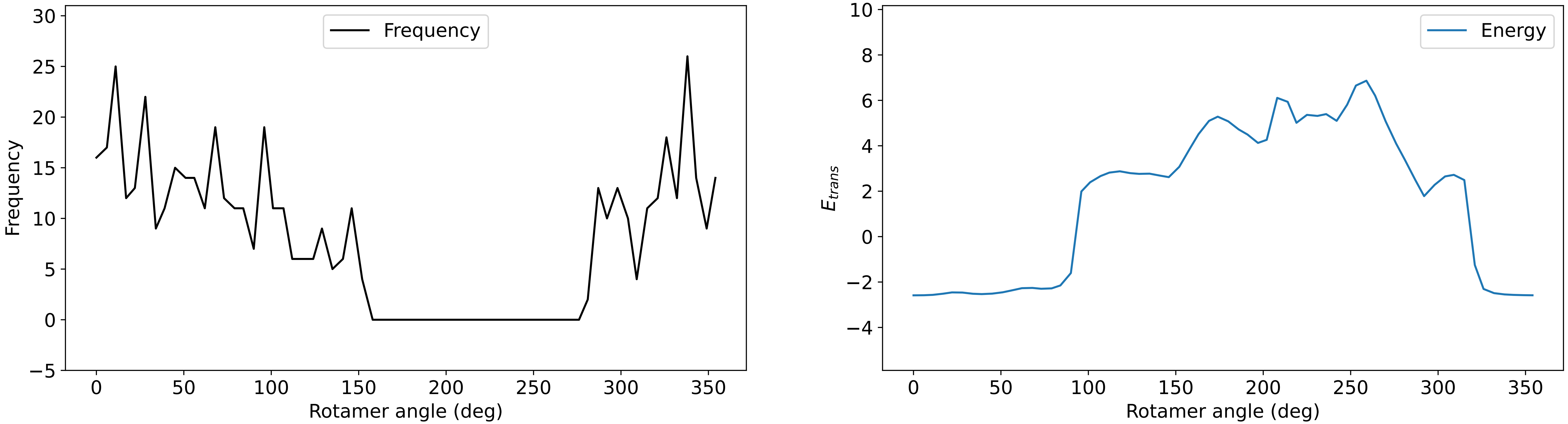

### Profile_49.pdf

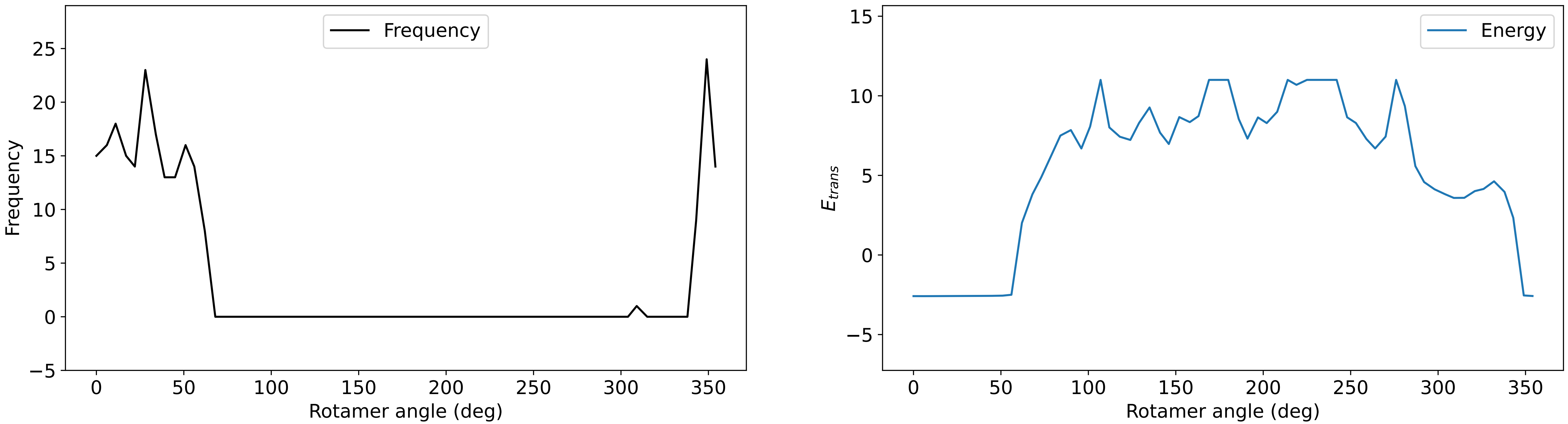

### Profile_50.pdf

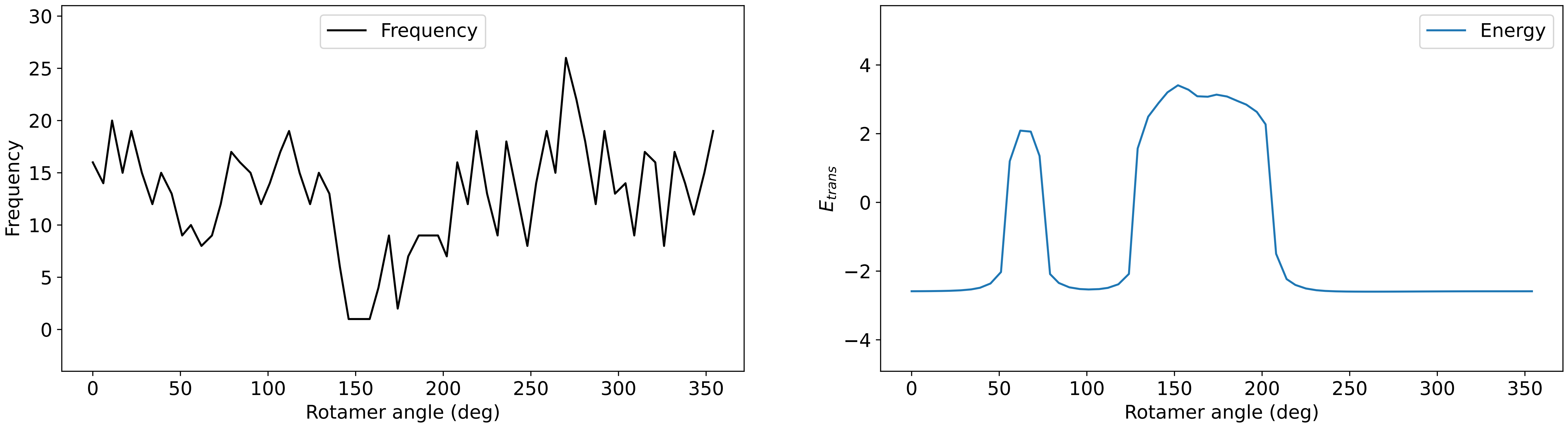

### Profile_52.pdf

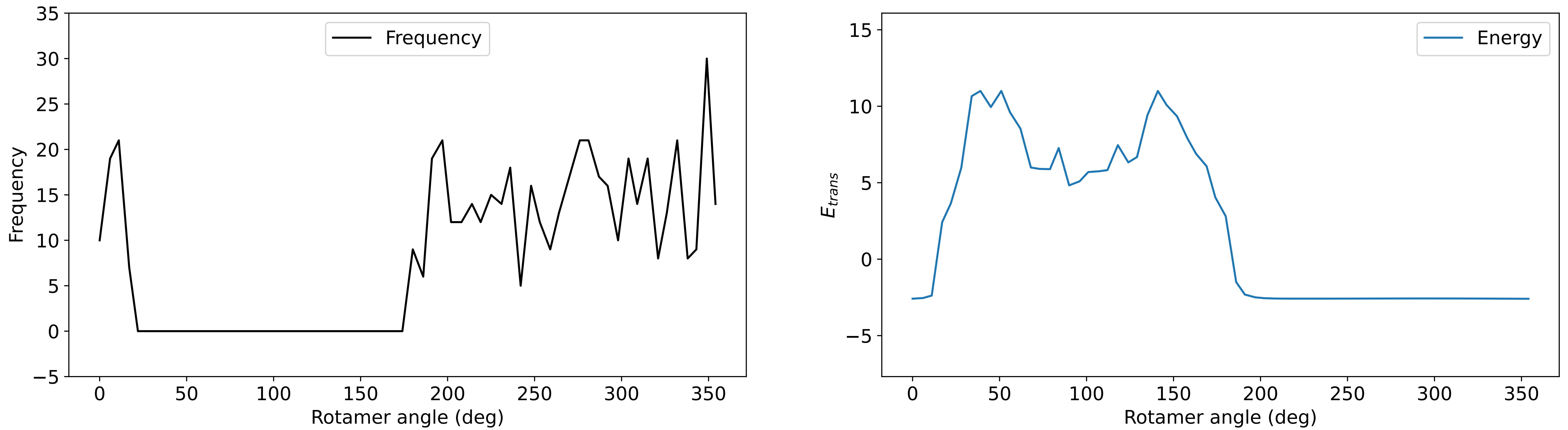

### Profile_53.pdf

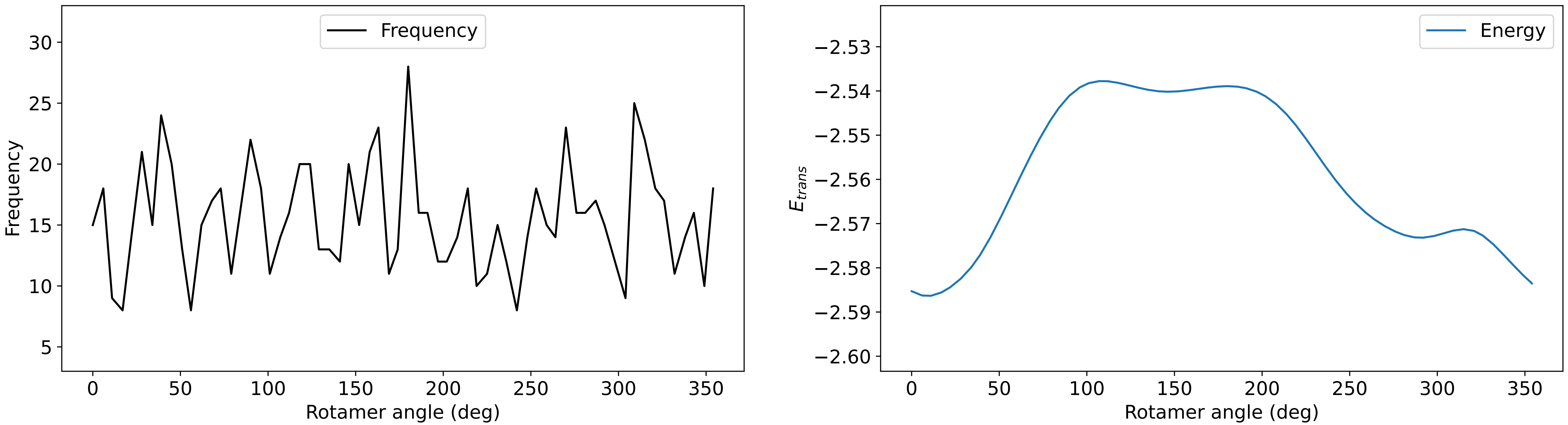

### Profile_54.pdf

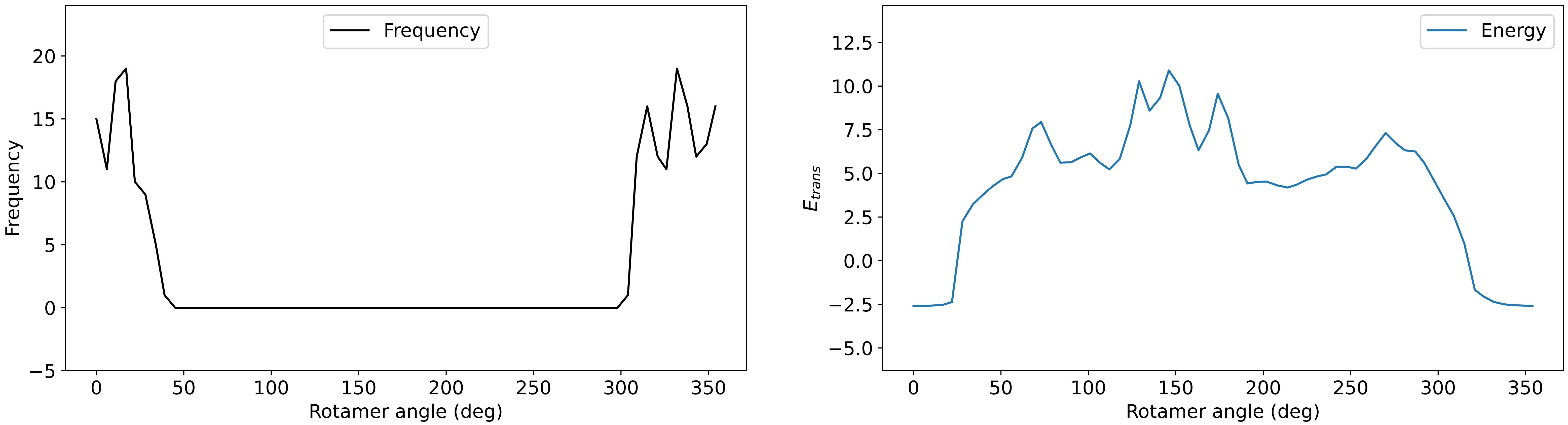

### Profile_55.pdf

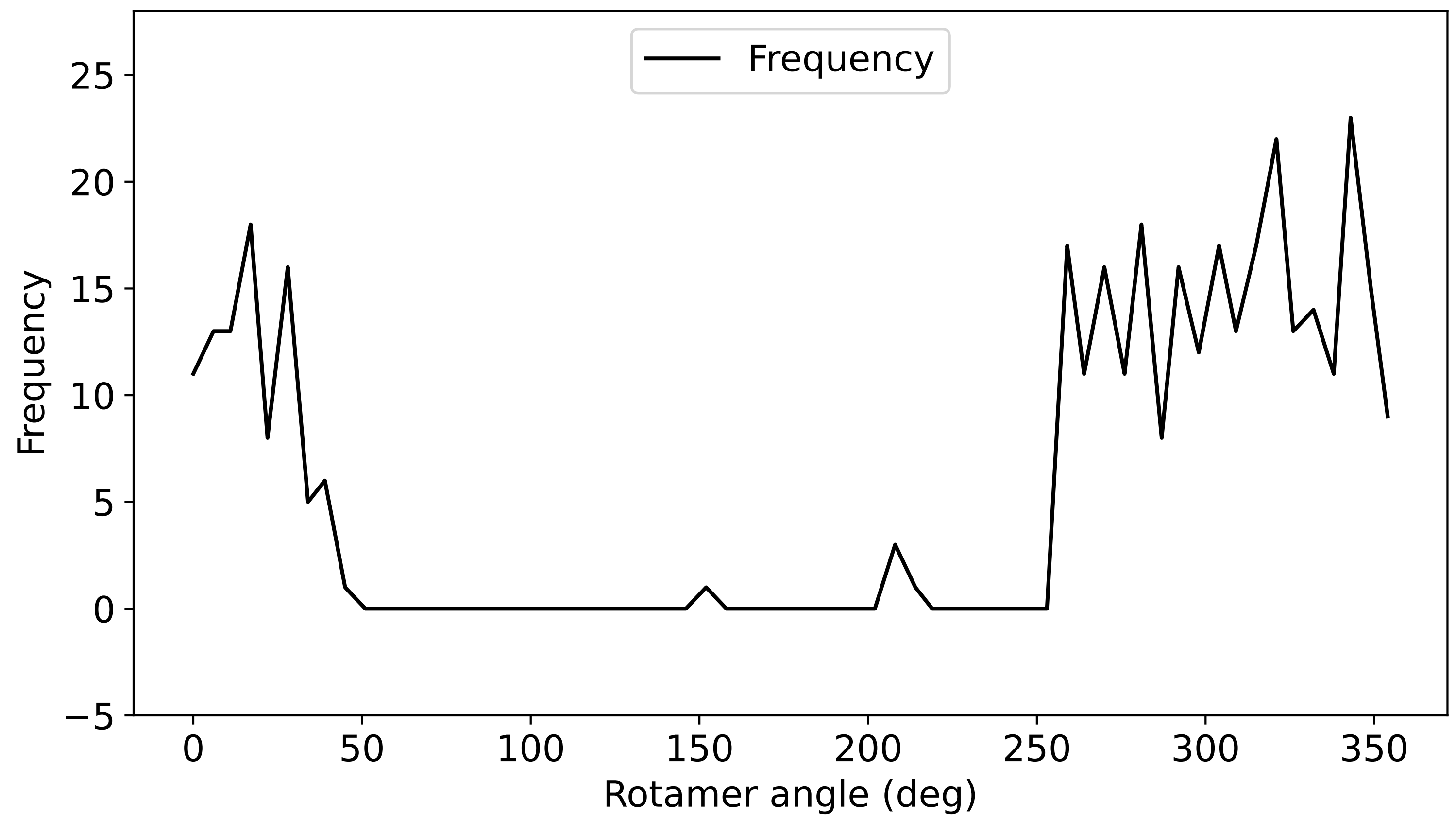

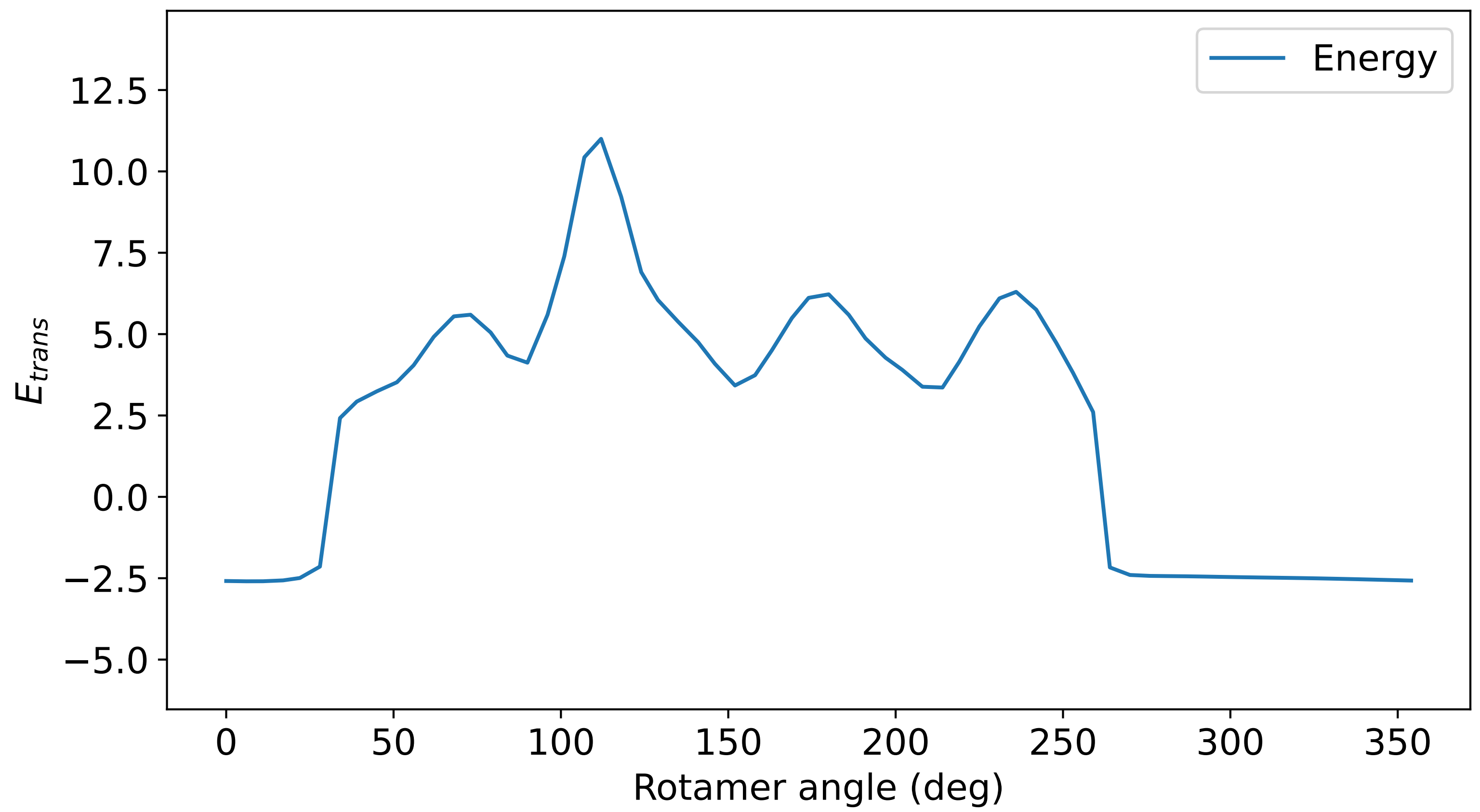

### Profile_57.pdf

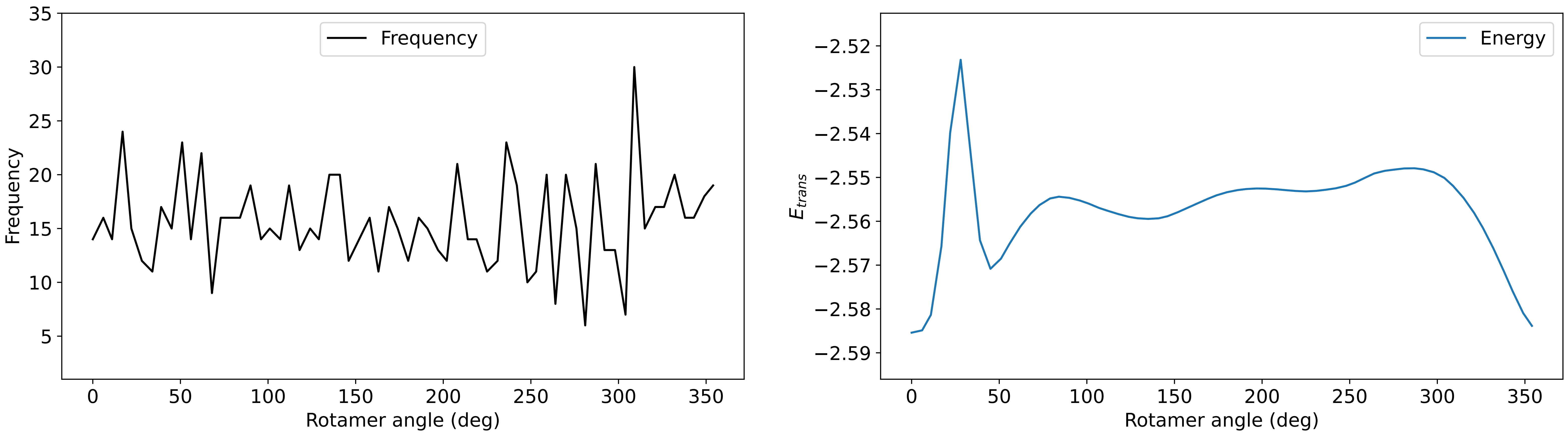

### Profile_58.pdf

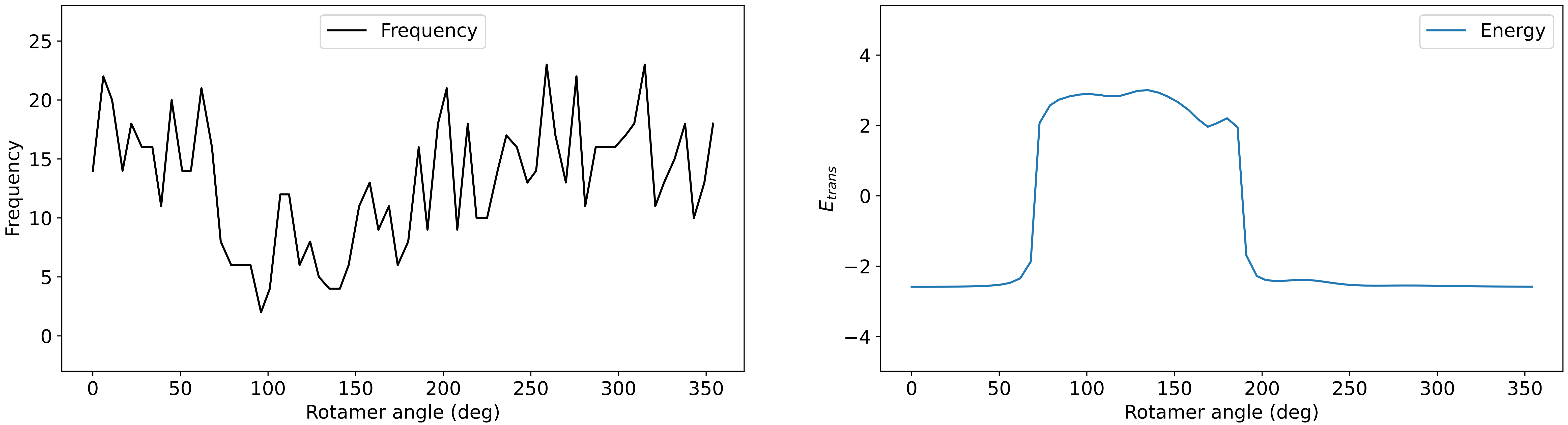

### Profile_59.pdf

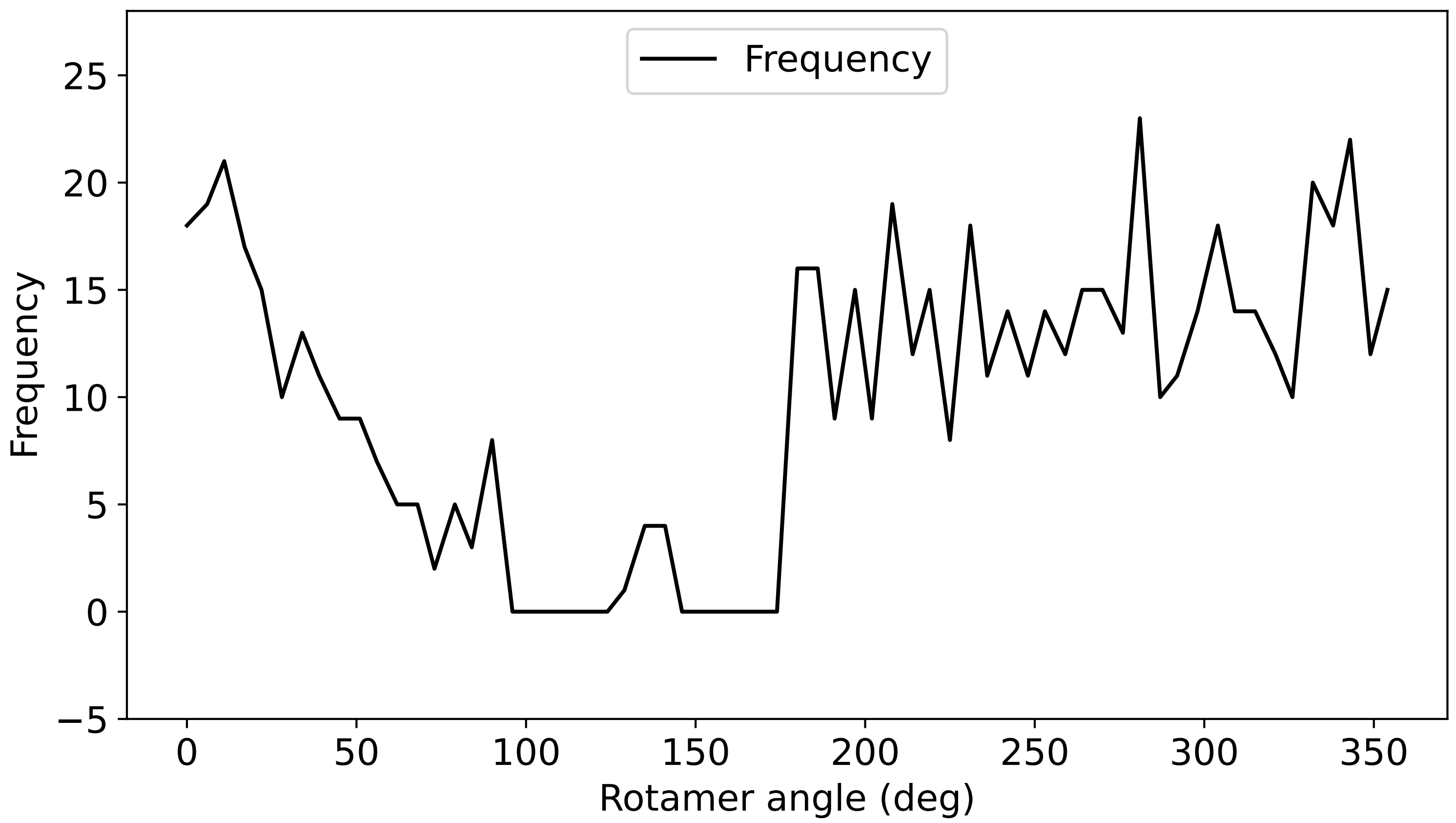

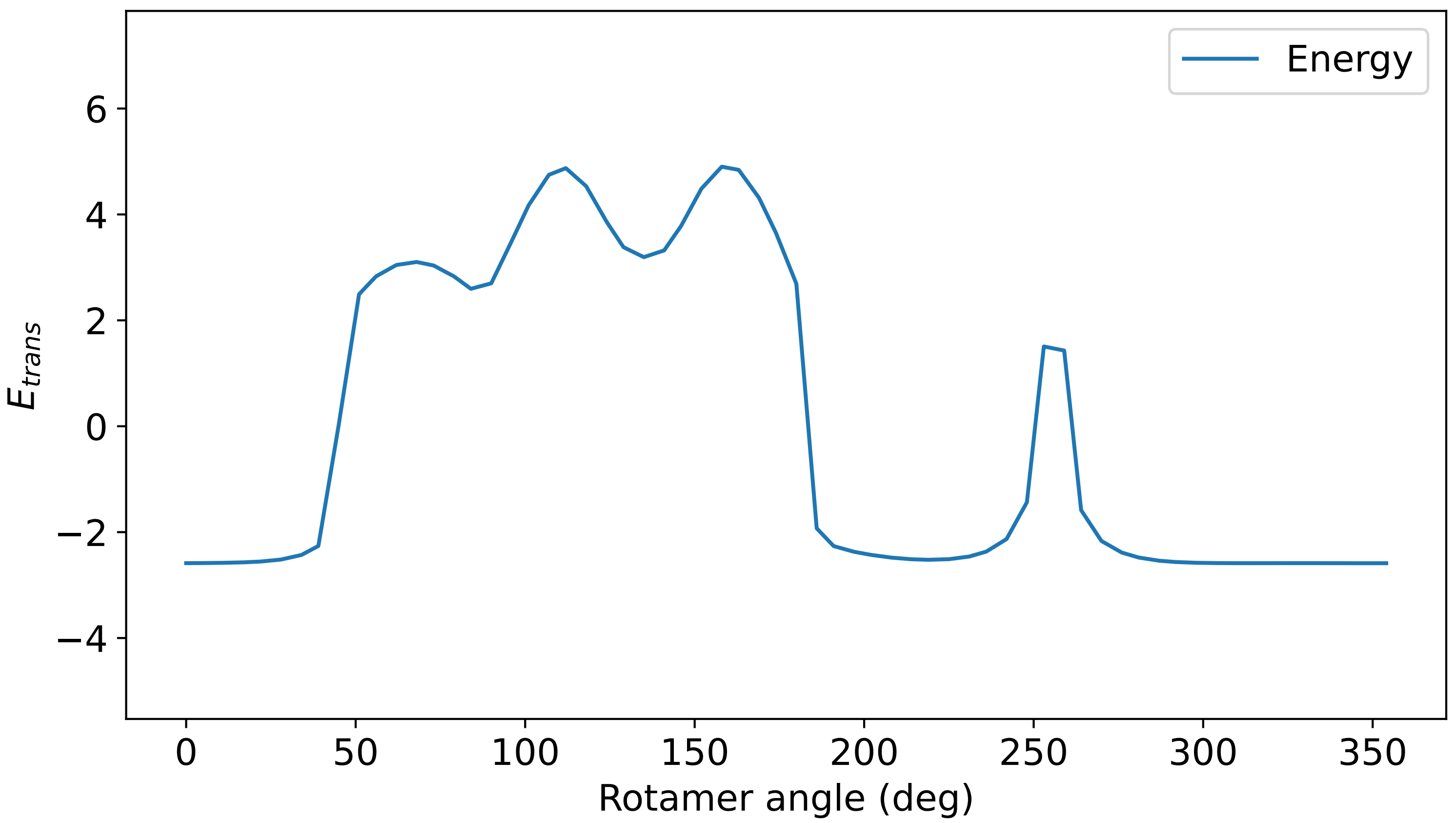

### Profile_60.pdf

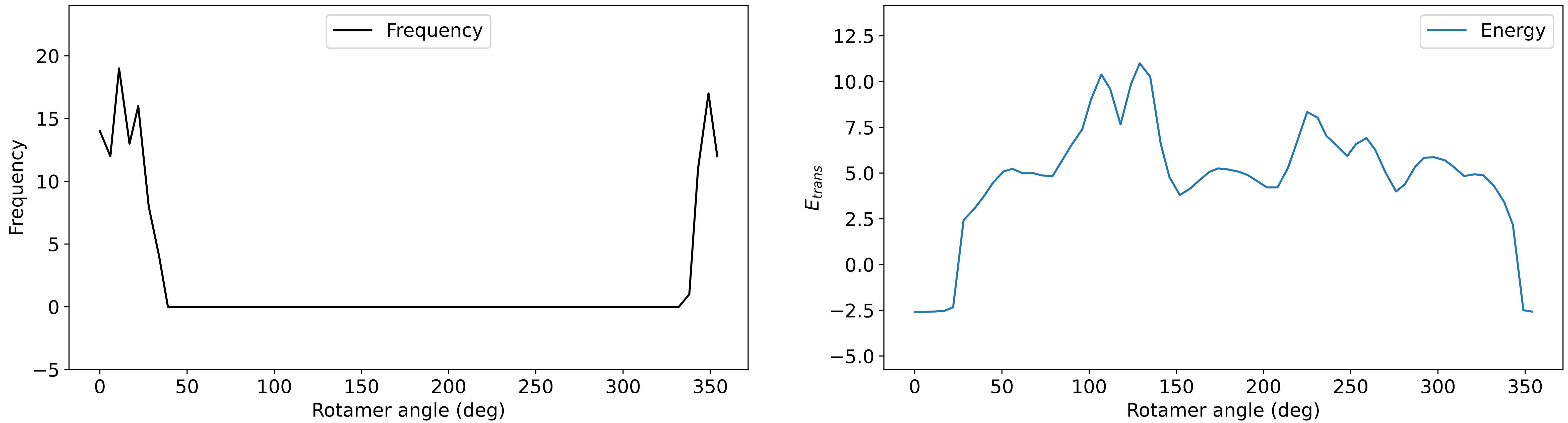

### Profile_61.pdf

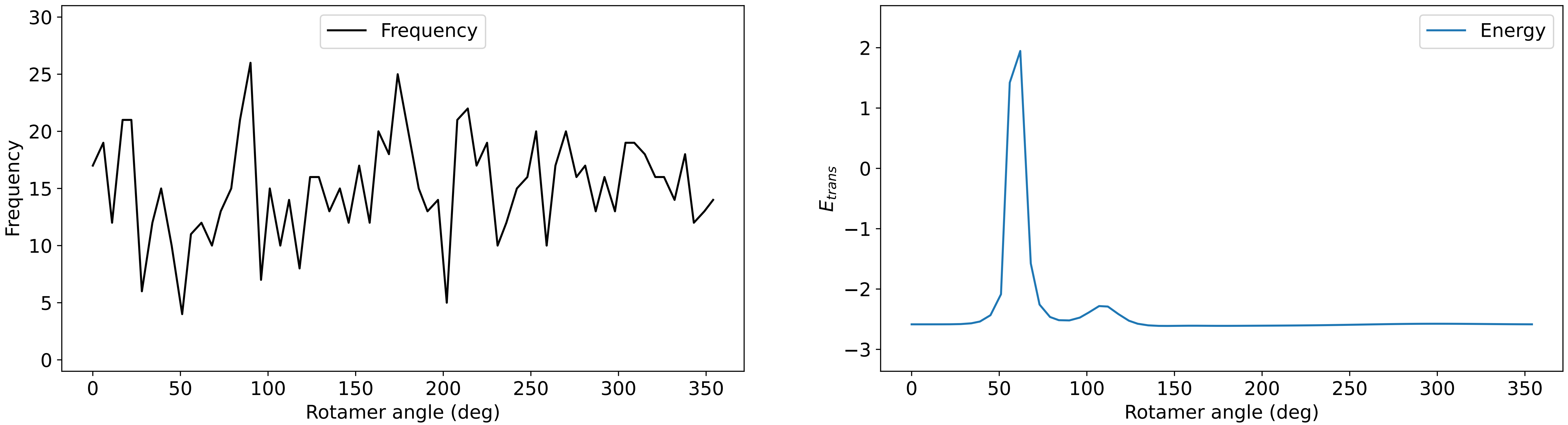

### Profile_62.pdf

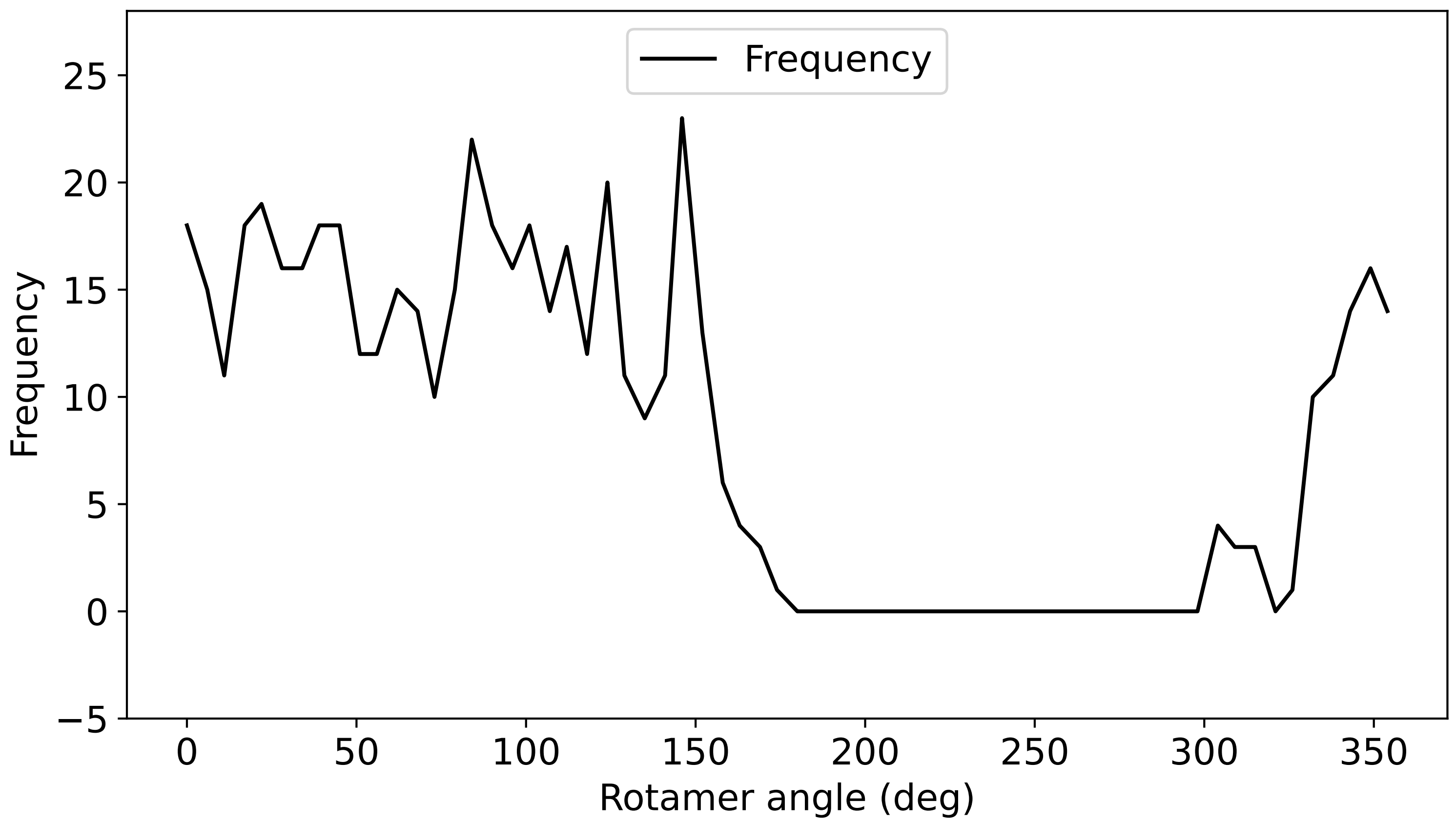

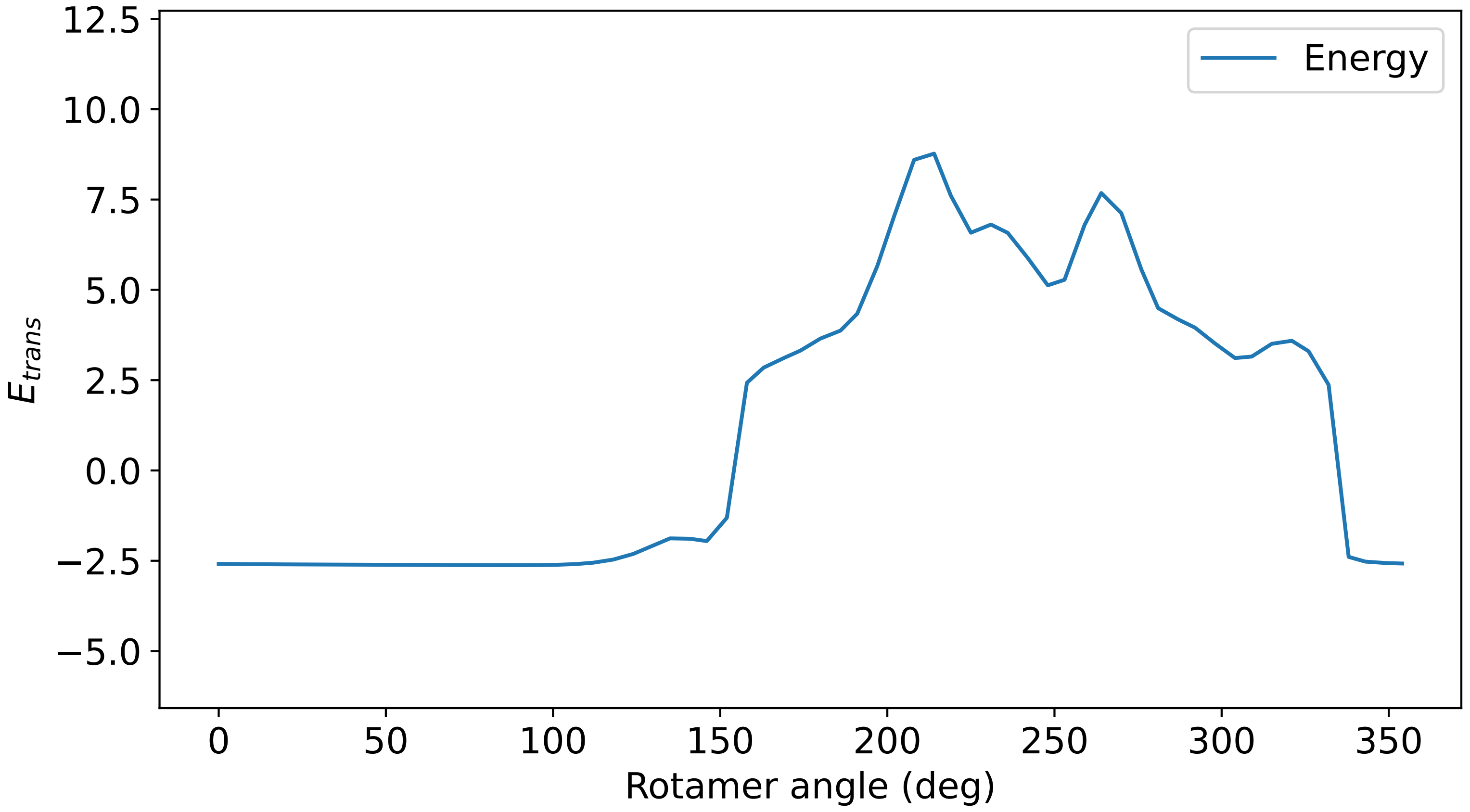

### Profile_63.pdf

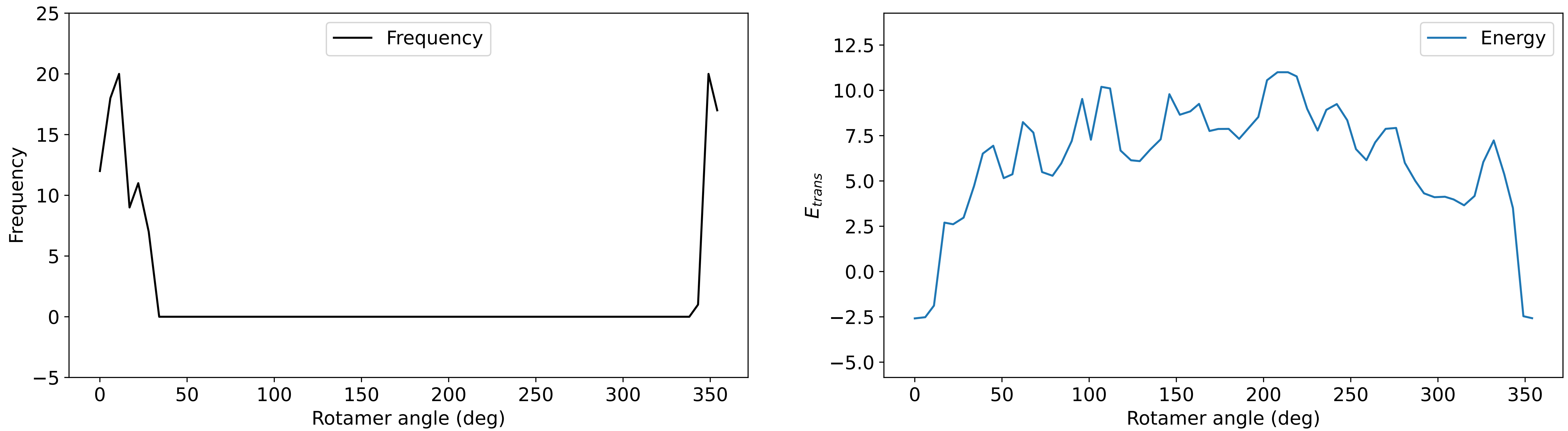

### Profile_64.pdf

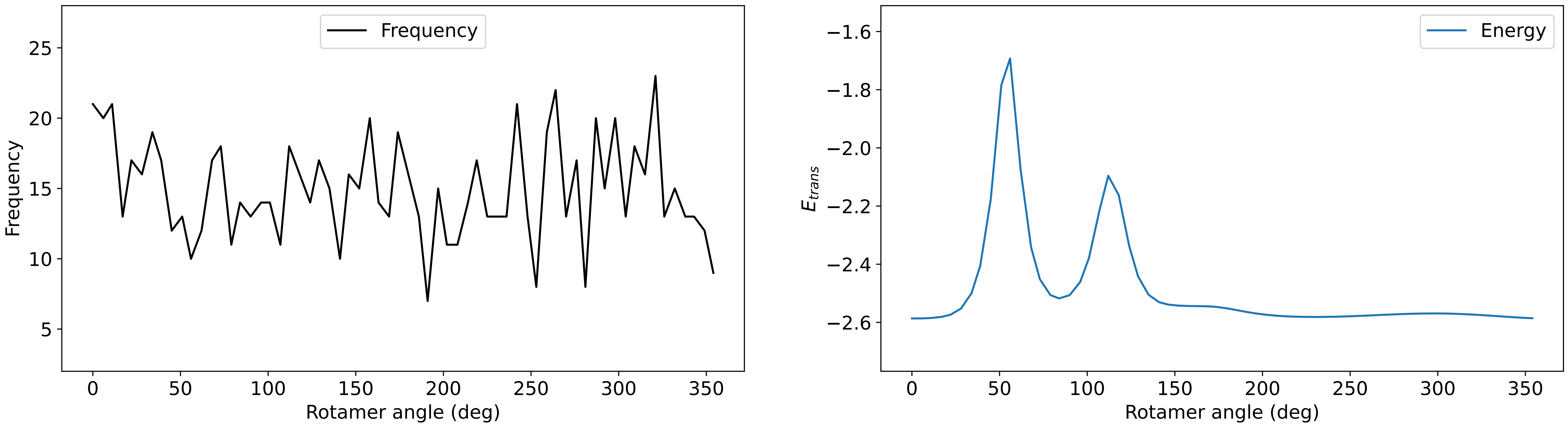

### Profile_65.pdf

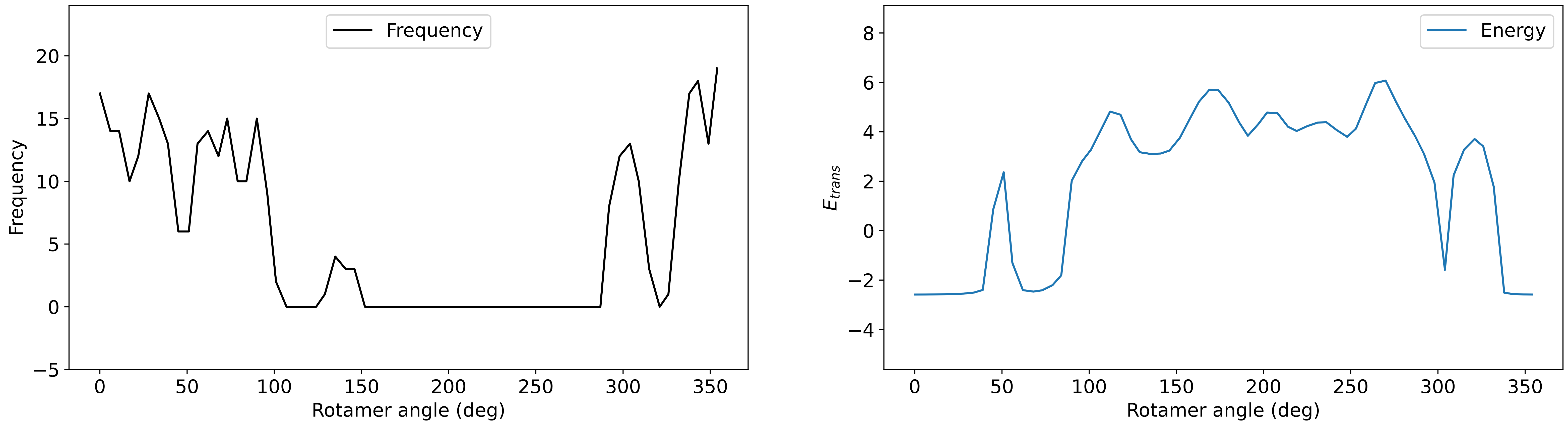

### Profile_66.pdf

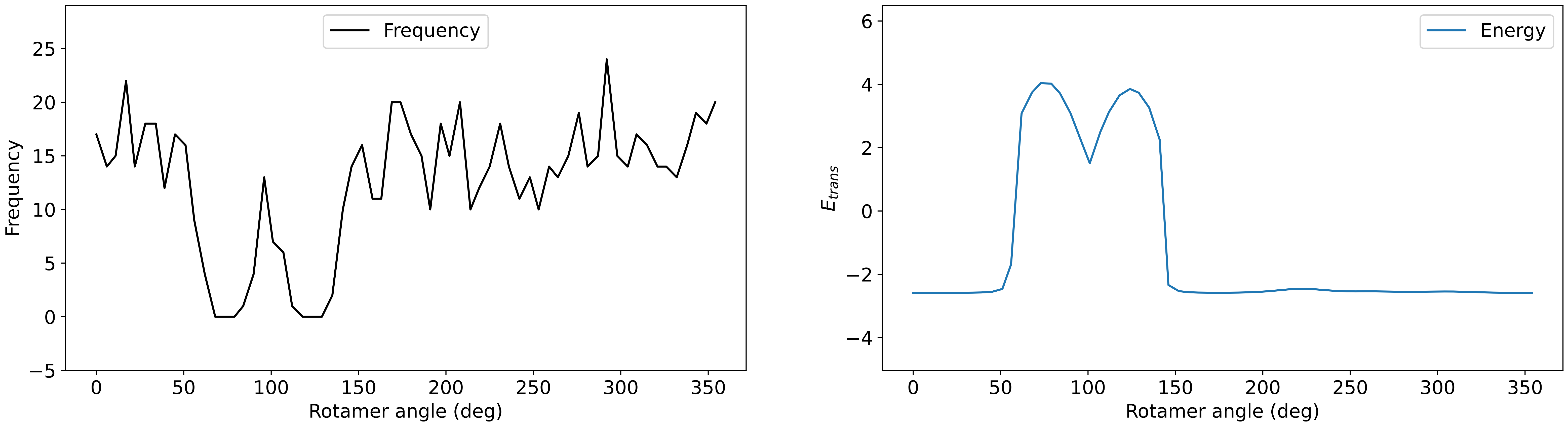

### Profile_69.pdf

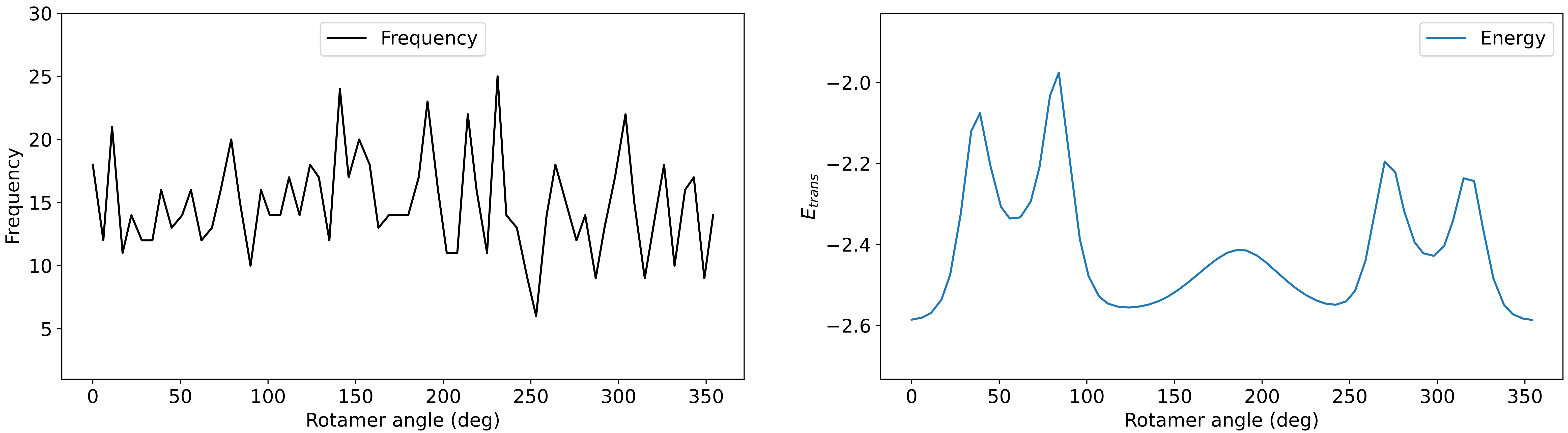

### Profile_70.pdf

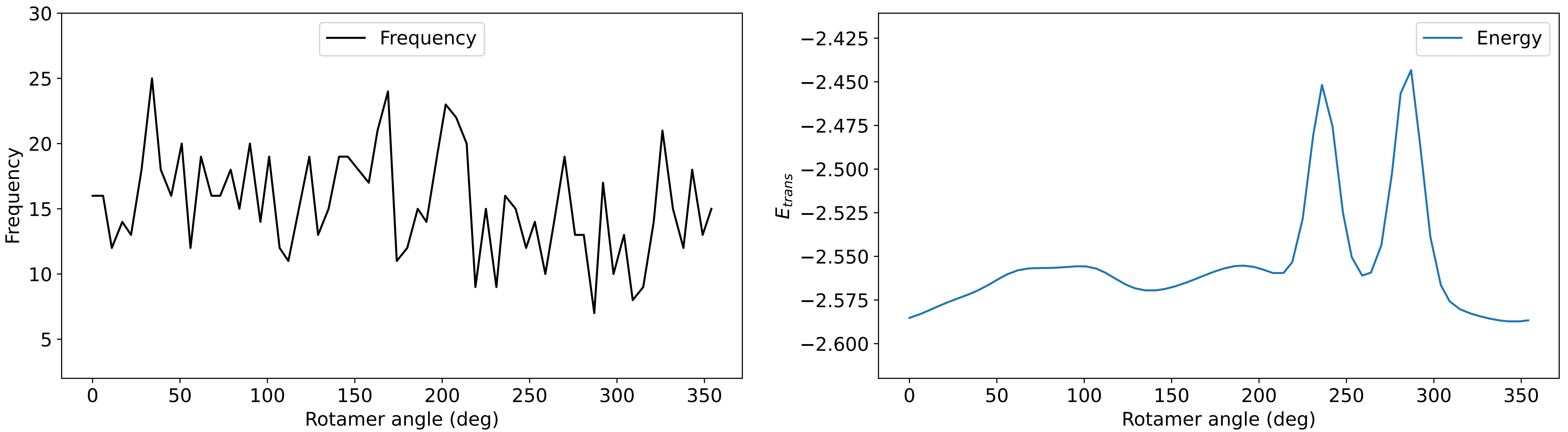

### Profile_71.pdf

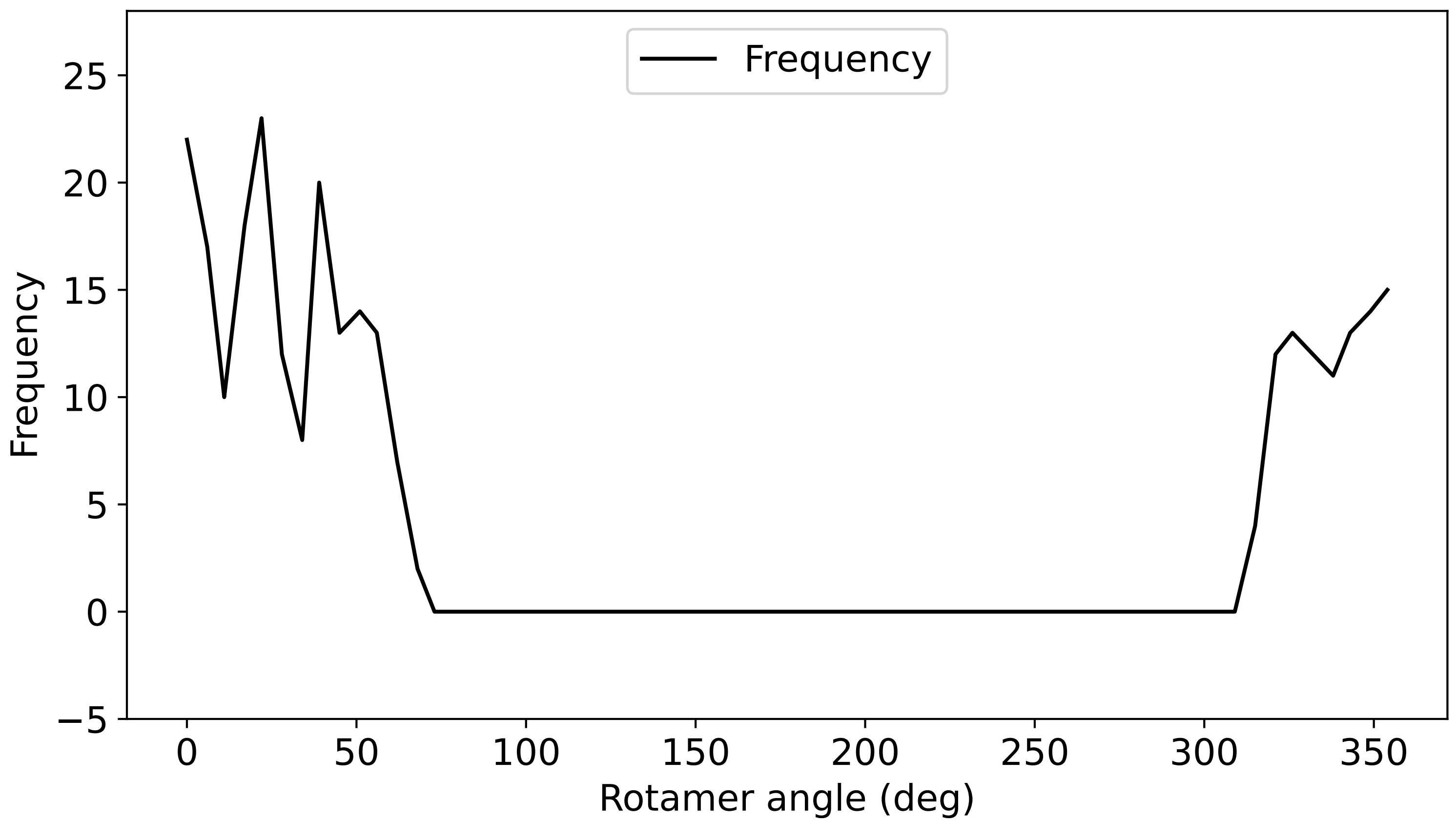

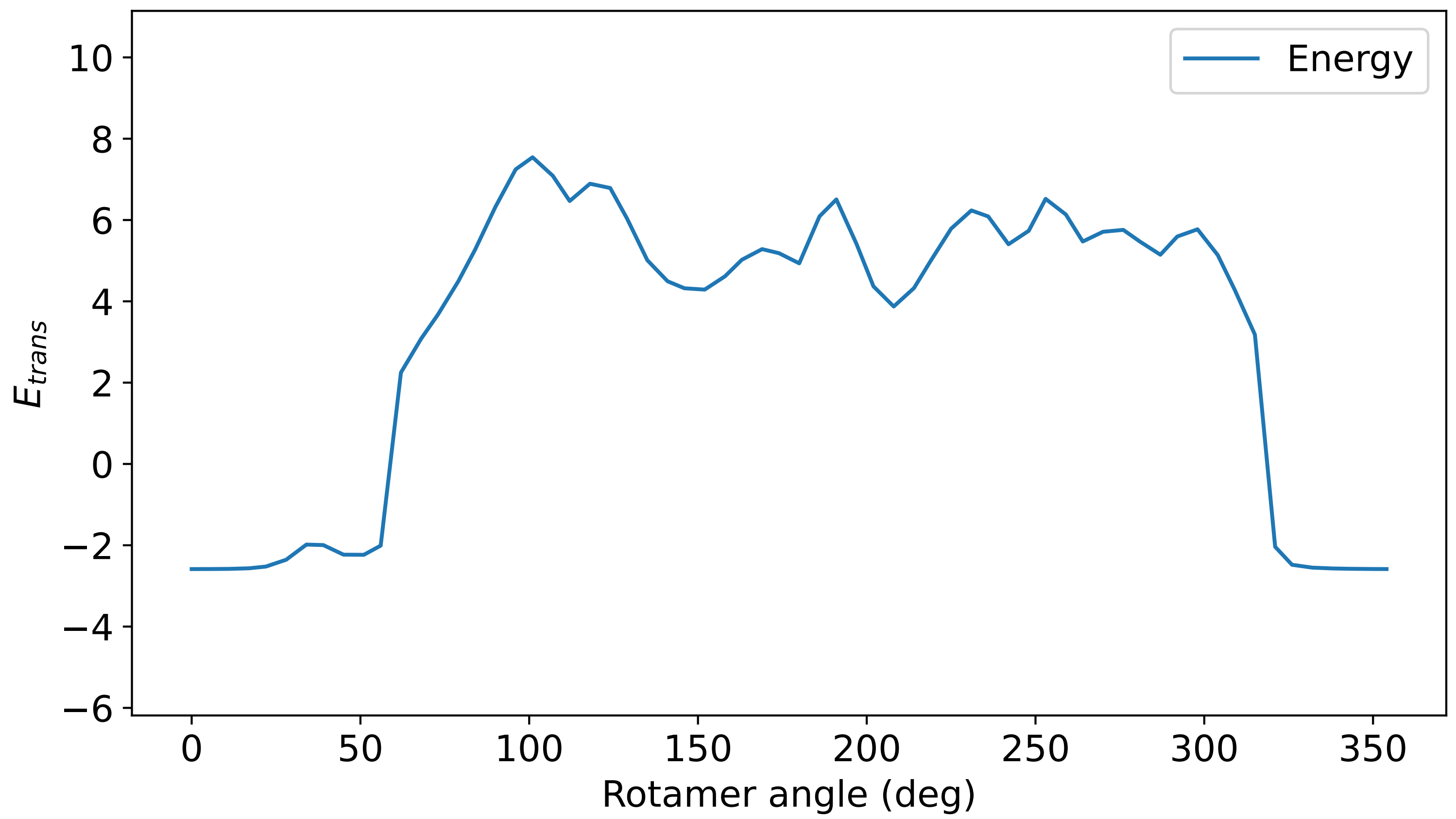

### Profile_72.pdf

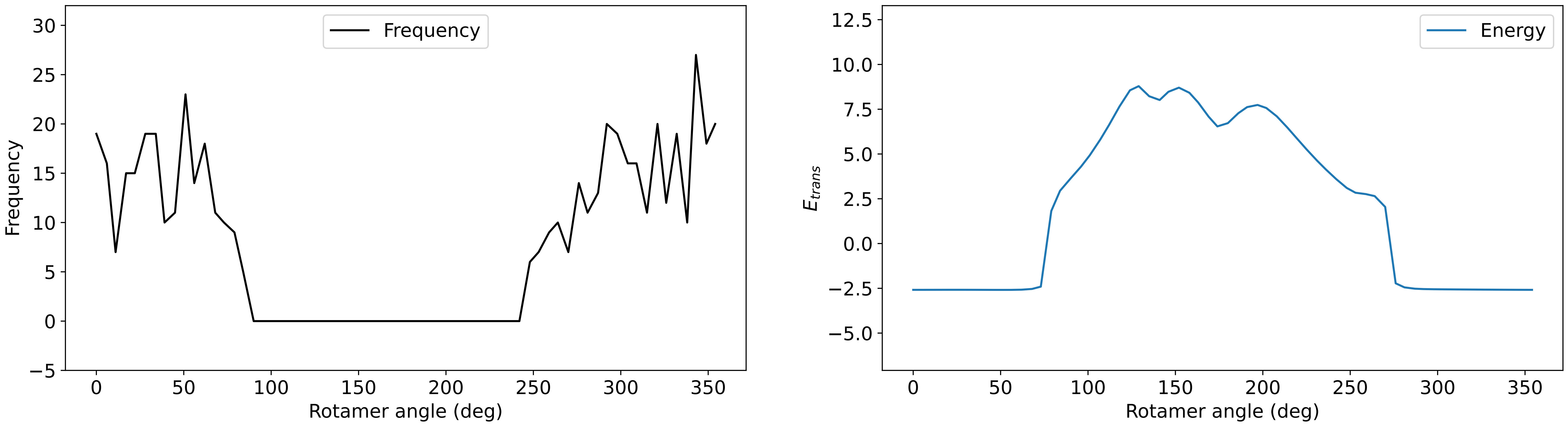

### Profile_73.pdf

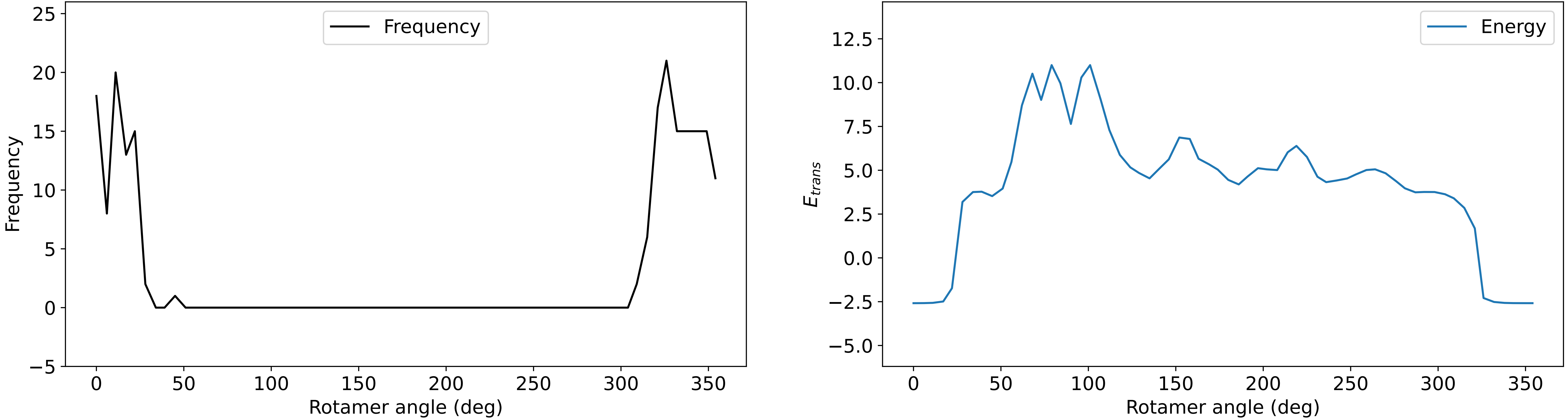

### Profile_74.pdf

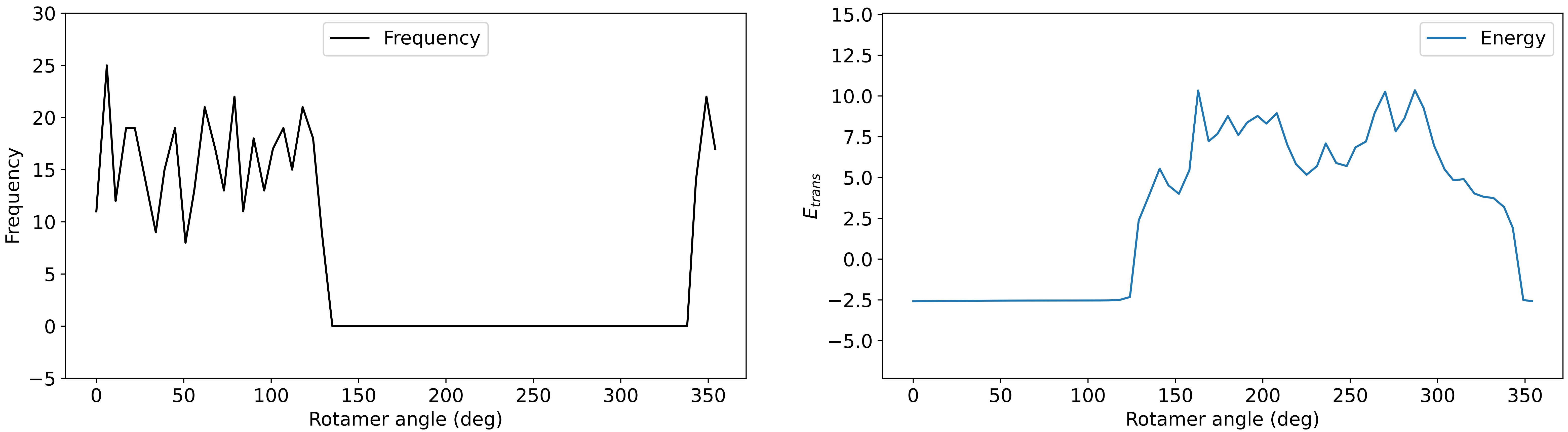

### Profile_75.pdf

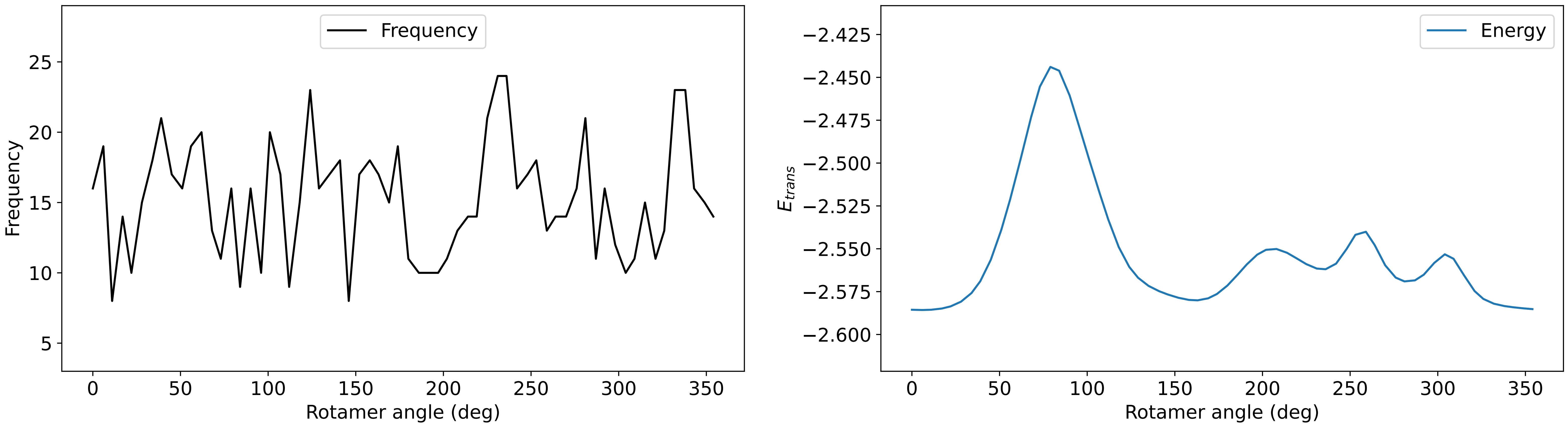

### Profile_76.pdf

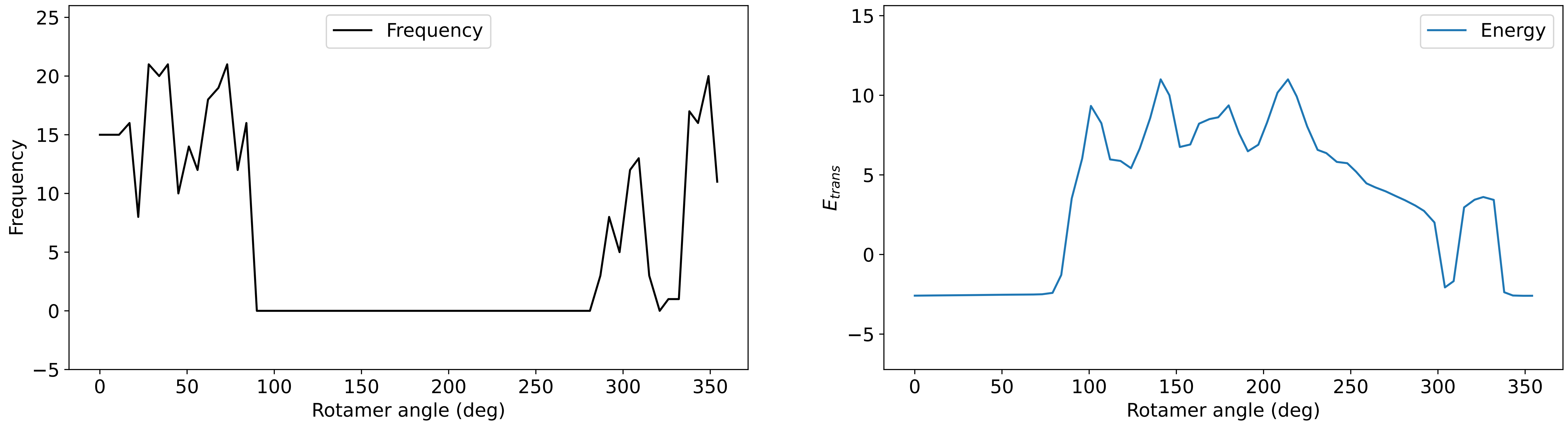

### Profile_77.pdf

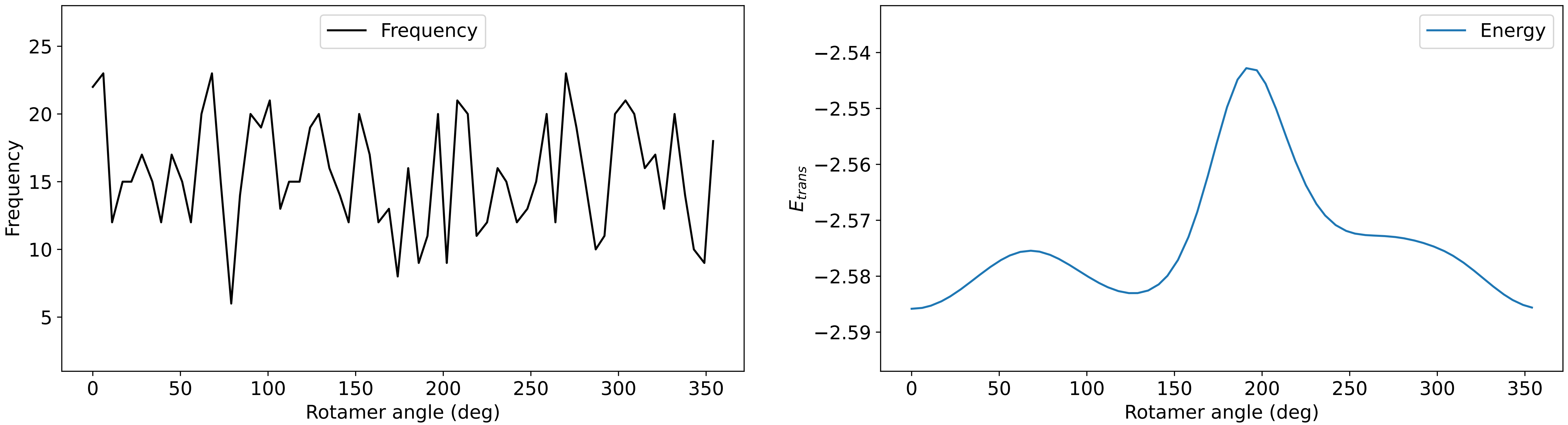

### Profile_78.pdf

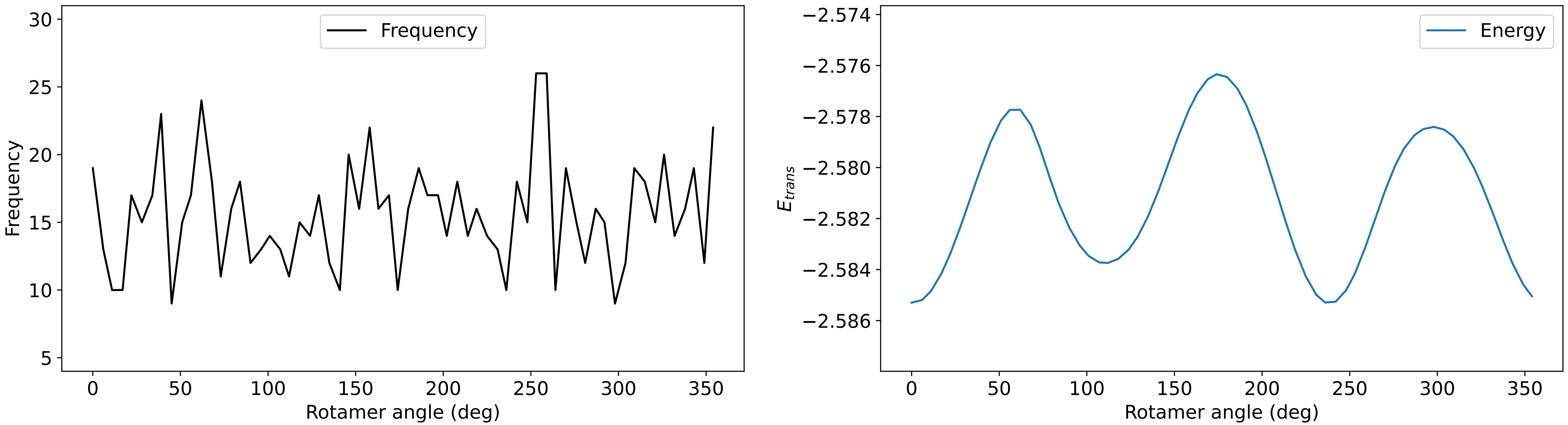

### Profile_80.pdf

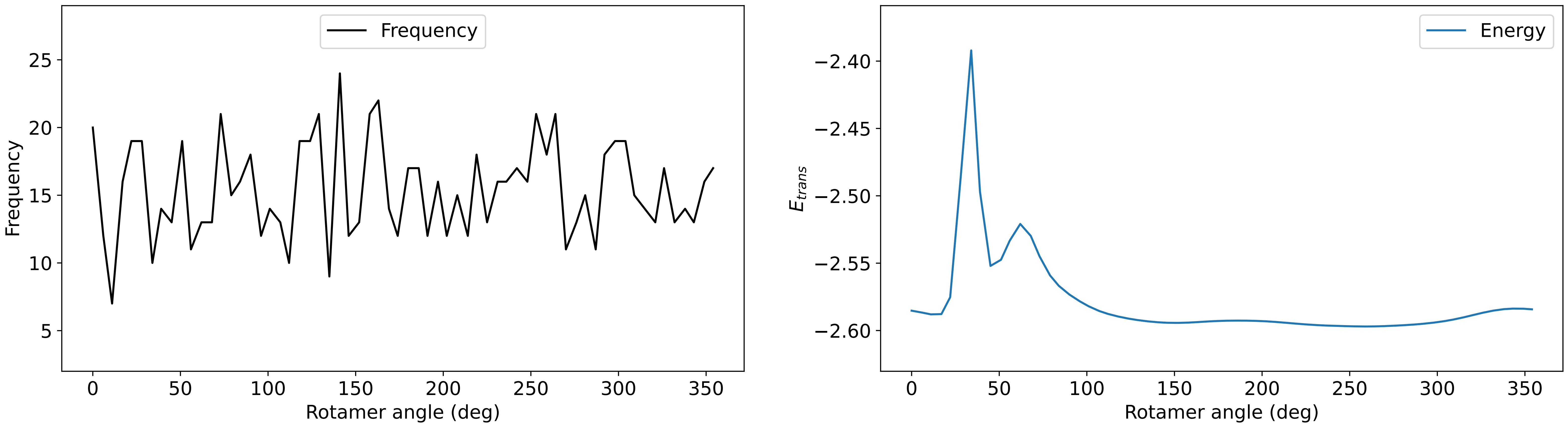

### Profile_82.pdf

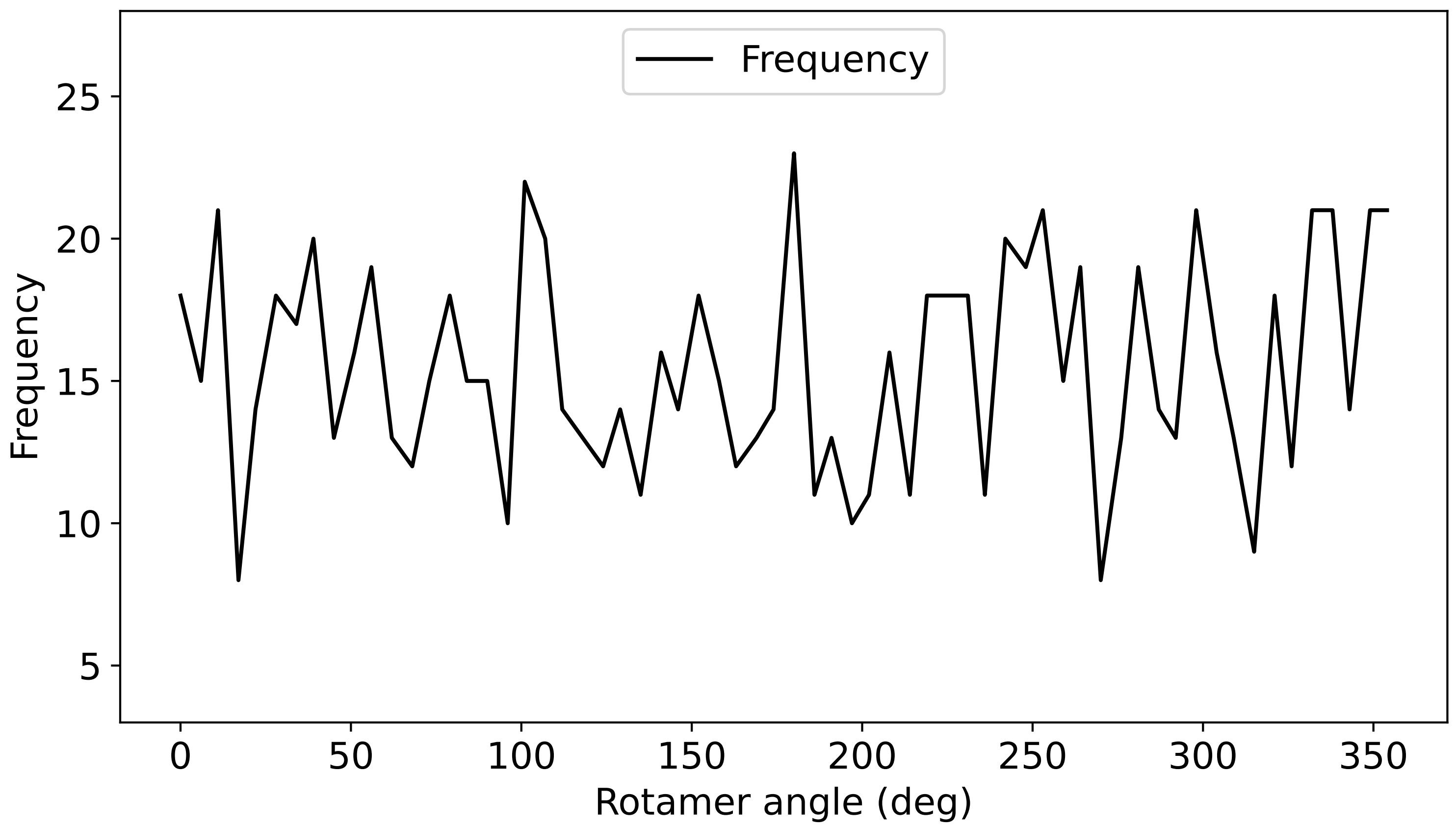

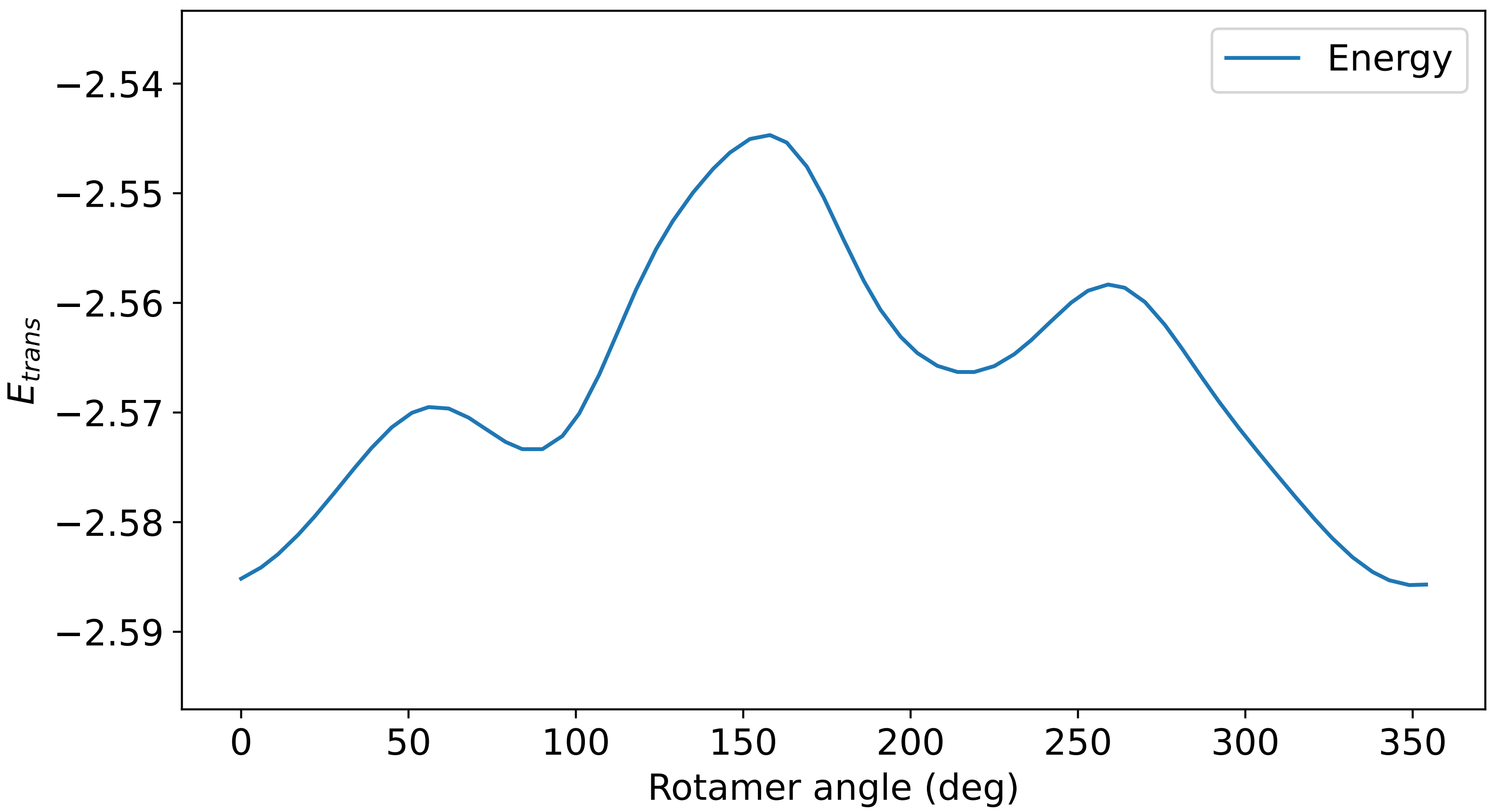
